# Ruthenium Coordination Architecture Redirects Imiquimod Antiviral Activity

**DOI:** 10.64898/2026.11.16.765078

**Authors:** Pon Ganish Prakash, Ana C. Carrasco, Greta Bajetto, Naveen Kumar, Thomas Iellici, Maria Jesus Moran Plata, Riccardo Moro, Andrea Bileck, Christopher Gerner, Matteo Biolatti, Hans-Joachim Mollenkopf, Rajendra Kumar Gurumurthy, Miguel A. Huertos, Samuel M. Meier-Menches, Valentina Dell’Oste, Cindrilla Chumduri, Luca Salassa

## Abstract

Immune agonists can reinforce antiviral defence, but therapeutic efficacy requires control over the magnitude and quality of the host response. Here we show that metal coordination can reprogramme the pharmacology of imiquimod, a clinically used TLR7 agonist. A series of Ru(II)-arene-imiquimod complexes identified the trinuclear compound **7** as a lead with activity against Herpes simplex virus types 1 and 2 (HSV-1 and HSV-2), including clinical isolates and antiviral-resistant strains. Compound **7** directly reduced HSV-1 infectivity and broadly suppressed viral protein accumulation in infected fibroblasts. In patient-derived human ectocervical organoids, it reduced epithelial infection and the recovery of infectious virus. Transcriptomic and proteomic profiling showed that **7** selectively restored infection-suppressed interferon-responsive and antiviral restriction programmes. Thus, Ru coordination does not simply enhance imiquimod activity but redirects its pharmacology, coupling virus-directed activity to restoration of epithelial antiviral defence. These findings establish a broader principle: coordination chemistry can engineer the response profile of immune-active anti-infective agents and expand the design space of host-directed antiviral therapeutics.

## Introduction

Immune agonists offer an attractive route to antiviral therapy because they can reinforce host defence rather than acting exclusively on a viral target. Imiquimod, a synthetic imidazoquinoline and clinically used TLR7 agonist, provides a well-established example of this strategy. It is approved for topical treatment of external genital warts, actinic keratosis and superficial basal cell carcinoma, and activates innate and adaptive immune responses through induction of interferons, inflammatory mediators and cellular immunity^1^. Imidazoquinoline derivatives have consequently been explored as immune-active therapeutic and vaccine-adjuvant scaffolds. However, imiquimod commonly induces dose-dependent local inflammatory reactions, and broader immune stimulation can also produce mucosal or systemic adverse effects^1,2^. This raises a broader pharmacological challenge: how to harness antiviral immunity while controlling the magnitude and quality of the host response.

Coordination chemistry provides a potential route to such pharmacological reprogramming. Metal complexes occupy chemical space distinct from that of conventional organic molecules through their three-dimensional geometries, charge states, redox properties, ligand-exchange behaviour and modes of biomolecular interaction^3^. Direct coordination of bioactive ligands to metal centres, as well as conjugation to organometallic fragments, can alter stability, cellular uptake, distribution and target engagement and generate biological activities distinct from those of the parent molecules^4^. Metal-containing compounds have shown considerable promise in antibacterial and broader anti-infective drug discovery, including through screening of metal-complex libraries and incorporation of organometallic fragments into established pharmacophores^5,6^. Coordination compounds with antiviral activity have also been reported, but the field remains comparatively limited^7^. A recent Pt(IV) oxaliplatin prodrug carrying the related imidazoquinoline gardiquimod illustrates the broader potential of incorporating immune agonists into metal-containing therapeutic architectures, although this system relies on covalent prodrug conjugation rather than direct metal coordination^8^. Direct metal coordination of an immune-active imidazoquinoline as a strategy to reprogramme antiviral pharmacology has not, to our knowledge, previously been explored.

Imiquimod is particularly suited to testing this concept because its heterocyclic scaffold contains multiple nitrogen donors capable of metal coordination. We therefore investigated whether direct coordination of imiquimod to organometallic fragments could generate compounds whose pharmacology differs from that of the parent immune agonist rather than simply increasing its activity. Herpes simplex viruses provide a stringent system in which to test this possibility. Herpes simplex virus types 1 and 2 (HSV-1 and HSV-2) establish lifelong latent infections and periodically reactivate, causing recurrent mucocutaneous disease and severe complications in neonates and immunocompromised individuals. Standard therapy remains centred on acyclovir and related nucleoside analogues that inhibit viral DNA replication; these drugs suppress productive infection but do not eliminate latency, and resistance is an important clinical problem in immunocompromised patients^9,10^. New viral targets, particularly the helicase–primase complex, are expanding the therapeutic landscape, but resistance and incomplete control of recurrent infection continue to motivate mechanistically distinct antiviral approaches^11^. Imiquimod has shown anti-HSV activity in experimental models and selected clinical studies, but its efficacy in recurrent HSV has been inconsistent and can be accompanied by substantial local inflammation^12,13,14,15^. Together, these observations make HSV an informative system in which to determine whether direct metal coordination can redirect the antiviral pharmacology of an immune-active scaffold.

Here we coordinated imiquimod directly to Ru(II)-arene scaffolds and generated a focused series of Ru-imiquimod complexes with distinct chemical and biological properties. We identify the trinuclear compound **7** as a lead derivative with activity against HSV-1 and HSV-2, including clinical isolates and antiviral-resistant strains. Compound **7** directly reduces HSV-1 infectivity and broadly suppresses viral protein accumulation in infected fibroblasts. In patient-derived human ectocervical organoids, it reduces epithelial infection and infectious-virus recovery while selectively restoring infection-suppressed interferon-responsive and antiviral restriction programmes. Together, these findings show that Ru coordination reprogrammes imiquimod pharmacology, generating a distinct antiviral profile characterised by direct antiviral activity and infection-dependent restoration of epithelial antiviral programmes.

## Results

### Ru coordination generates distinct imiquimod architectures

We prepared seven Ru(II)-arene imiquimod complexes of general formula [Ru(η^6^-p-cym)(imiquimod)(Cl/Z)]^n+^ (**Fig. 1a**), in which imiquimod acts as a bidentate ligand and the remaining coordination site is occupied by chloride (**1**), or a monodentate Z ligand such as pyridine (**2**), 1,3,5-triaza-7-phosphaadamantane (**3**), imidazole (**4**), indazole (**5**) or 5-fluorouracil (**6**). Complexes **1**–**6** were obtained by adapting established Ru-arene substitution chemistry^16–19^. Attempts to introduce 2-aminoadamantane instead yielded the trinuclear derivative **7**, in which the Z site is occupied by the quinoline nitrogen of a neighbouring Ru-imiquimod unit following deprotonation of the imiquimod amino group and bridging coordination, consistent with related higher-nuclearity Rh-arene assemblies^20^.

**Figure 1:**
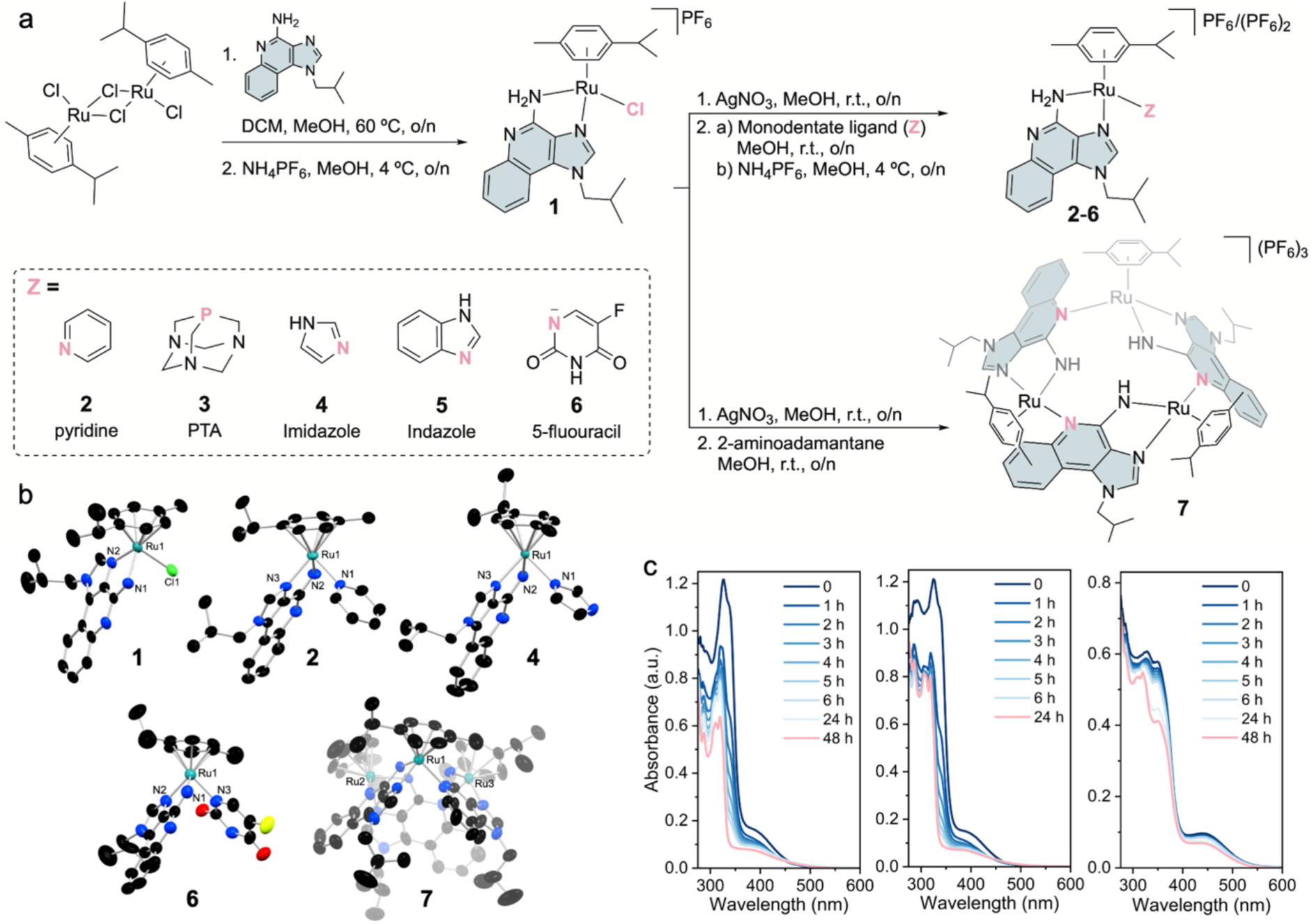
Ru coordination generates distinct imiquimod architectures and solution behaviour. **a**, Synthetic scheme for complexes **1**–**7**, isolated as PF_6_ salts with mono-(**1**, **6**), di-(**2**–**5**) or tricationic (**7**) charge states. **b**, X-ray structures of **1**, **2**, **4**, **6** and **7**. Displacement ellipsoids are shown at 50% probability; H atoms, PF ^−^ anions and solvent molecules are omitted for clarity. **c**, Evaluation of the stability of **1**, **6**, (100 μM) and **7** (25 μM) in DMEM cell culture media (5% DMSO) by UV-vis spectroscopy.

All compounds were characterised by multinuclear NMR spectroscopy, mass spectrometry and elemental analysis (**Figs**. **S1–S17**), and single-crystal X-ray diffraction confirmed the structures of **1**, **2**, **4**, **6** and **7** (**Fig. 1b** and **Table S1**). The complexes were stable as solids under ambient conditions for several weeks. Their solution behaviour was examined by ^1^H NMR spectroscopy in methanol and DMSO (**Figs**. **S18–S30**). Complexes **1**–**6** remained unchanged in methanol, whereas **7** was insufficiently soluble for analysis. In DMSO, **1** showed partial conversion after 24 h (∼30%), while **7** was largely inert (∼7% conversion), with no detectable release of free imiquimod. Complexes **2**–**6** showed variable Z-ligand lability, with **5** converting rapidly upon dissolution and **3** remaining unchanged over 48 h.

Solution studies in cell-culture medium were focused on the most biologically active derivatives, **1**, **6**, and **7** (*vide infra*). In DMEM containing 5% DMSO, **1** and **6** slowly evolved to species consistent with aquation and Z-ligand release, whereas **7** retained its characteristic 450-nm absorption band with minimal degradation over 48 h (**Fig. 1c**). These findings establish substantial differences in coordination stability and ligand lability across the Ru-imiquimod series, with **7** displaying a distinct and comparatively stable trinuclear architecture.

### Ru–imiquimod complexes inhibit HSV replication

We first assessed the cytotoxicity of the Ru-imiquimod series in HFF, HeLa and VERO cells by MTT assay, defining as non-cytotoxic the concentrations that preserved at least 70% viability after 48 h. All metalated derivatives were less cytotoxic than free imiquimod across the three cell lines (**Table S2** and **Extended Data Fig. 1a**). HSV-1 activity was then quantified by virus-yield reduction assays and expressed as plaque-forming units per millilitre (PFU ml⁻¹), reflecting infectious viral particles capable of forming plaques in a susceptible cell monolayer. Compounds were initially tested at non-cytotoxic concentrations of 12.5 μM (HeLa) or 25 μM (HFF and VERO) for **7**, and 25 μM across all cell lines for the remaining complexes. Compounds **1**, **2**, **4**, **5**, **6** and **7** showed significant antiviral activity in HeLa cells, whereas **7** was the only compound active in HFF and VERO cells, indicating cell-type-dependent efficacy. At 48 h post-infection, **7** reduced HSV-1 titres by at least 2.5 log units in both HFF and HeLa cells, while **6** produced a comparable reduction in HeLa cells. Compounds **1**, **6** and **7** were therefore prioritized for further analysis because they showed the strongest activity and broadest cell-line profiles, whereas **2–5** were active predominantly in HeLa cells (**Fig. 2a** and **Extended Data Fig. 1a**). The Ru-*p*-cymene precursor showed no comparable antiviral activity, indicating that the Ru fragment alone was insufficient for the observed effect (**Extended Data Fig. 1b,c**).

**Figure 2:**
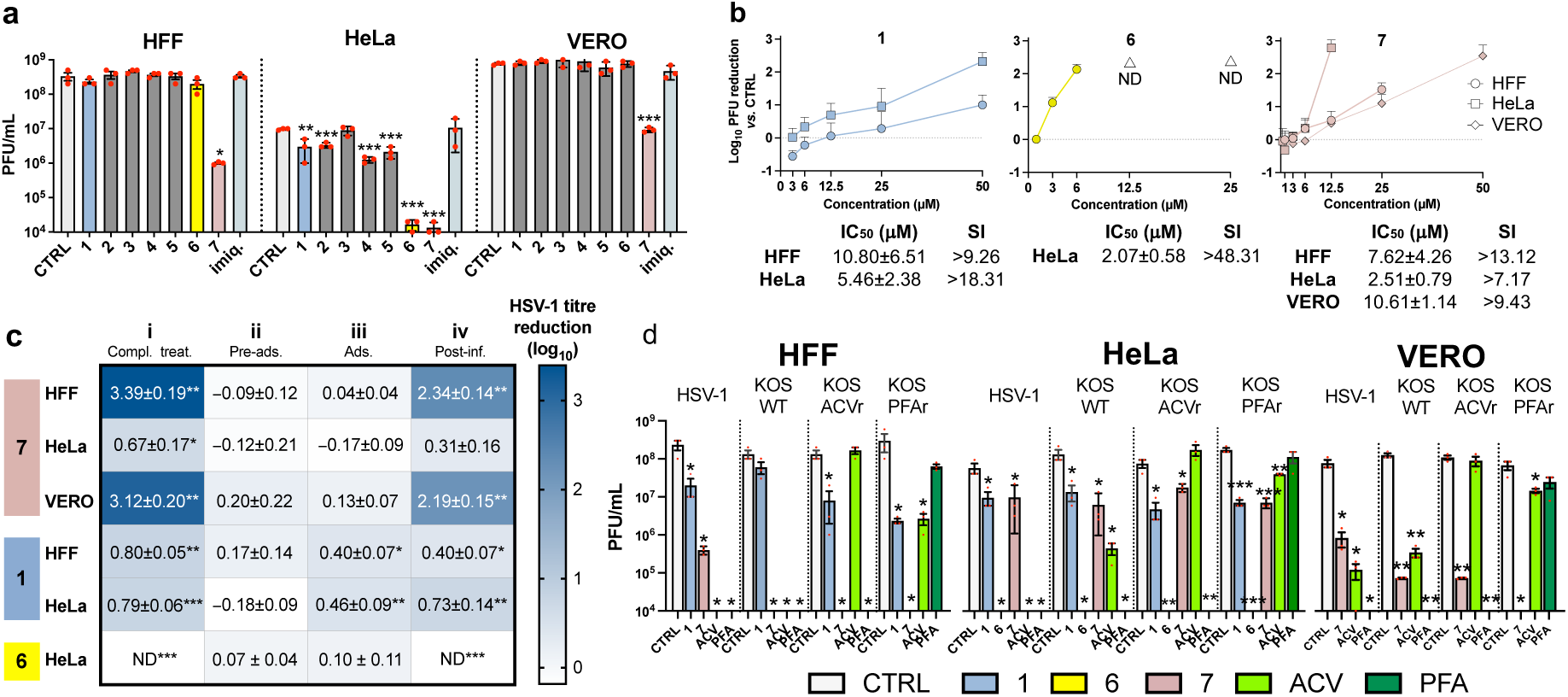
Ru–imiquimod complexes show broad antiviral activity against HSV-1. **a**, Virus-yield reduction assay of compounds **1**–**7** in HFF, HeLa and VERO cells infected with HSV-1 (MOI 0.1, 48 h). Cells were treated with the indicated compounds or vehicle control (CTRL); **7** was used at 12.5 μM in HeLa cells, whereas all other treatments were performed at 25 μM. **b**, Concentration-dependent antiviral activity of **1**, **6** and **7** in the indicated cell lines (MOI 0.1, 48 h), expressed as log_10_ reduction in viral titre relative to the paired CTRL. IC_50_ and selectivity index (SI) values are indicated. **c**, Heatmap summarizing the time-of-addition assays for **1**, **6** and **7**. Cells were treated with **1** (50 μM in HFF, 25 μM in HeLa), **6** (12.5 μM in HeLa) or **7** (25 μM in HFF, 12.5 μM in HeLa, 50 μM in VERO) according to four schedules: (i) complete treatment, comprising 2 h of pre-treatment, 2 h of virus adsorption in the presence of compound and 48 h of post-infection treatment; (ii) pre-treatment for 2 h only; (iii) treatment during virus adsorption for 2 h only; or (iv) post-infection treatment for 48 h only. In all conditions, HSV-1 infection was performed at MOI 0.1, and compounds were removed between treatment phases as appropriate. Heatmap colours represent the mean log_10_ reduction in viral titre relative to the paired CTRL, with positive and negative values indicating decreased and increased viral yields, respectively. **d**, Antiviral activity against HSV-1, the KOS reference strain (KOS WT), and acyclovir- or phosphonoformate-resistant KOS strains (KOS ACVr and KOS PFAr). Cells were treated with **1** (50 μM in HFF, 12.5 μM in HeLa), **6** (12.5 μM in HeLa), **7** (25 μM in HFF, 12.5 μM in HeLa, 50 μM in VERO), acyclovir (ACV, 8.9 μM), phosphonoformate (PFA, 1 mM) or CTRL. Viral yield was quantified by plaque assay on VERO cells using combined cell-associated and supernatant fractions. Data represent three independent experiments and are shown as mean ± SEM where applicable. Panels **a** and **d** report viral titres in plaque-forming units per millilitre (PFU/mL), whereas **b** and **c** report log_10_ viral titre reductions relative to CTRL. ND indicates that no infectious virus was detected by plaque assay (0 PFU); such results are not assigned numerical log_10_ reduction values. Where indicated, statistical significance was determined using unpaired t-tests comparing treatments with CTRL (p < 0.05 *, p < 0.01 **, p < 0.001 ***).

Dose–response analysis confirmed antiviral activity of **1**, **6** and **7**. Compound **1** inhibited HSV-1 replication with IC_50_ values of 10.80 ± 6.51 μM in HFF cells and 5.46 ± 2.38 μM in HeLa cells, corresponding to selectivity indices of 7.49 and 19.06, respectively. Compound **6** showed an IC_50_ of 2.07 ± 0.58 μM in HeLa cells, whereas **7** showed IC_50_ values of 7.62 ± 4.26, 2.51 ± 0.79 and 10.61 ± 1.14 μM in HFF, HeLa and VERO cells, respectively. Selectivity indices exceeded 7 in all responsive conditions and reached 48.31 for **6** in HeLa cells (**Fig. 2b** and **Extended Data Fig. 1d**). Time-of-addition experiments showed that pre-treatment alone was ineffective, whereas activity was retained when compounds were present after viral adsorption. Compounds **6** and **7** were most effective when maintained throughout infection or added after adsorption, with **6** nearly abolishing viral production in HeLa cells and **7** in HFF cells (**Fig. 2c**). These findings indicate that sustained exposure during post-entry stages of infection is important for antiviral activity.

ICP-MS analysis revealed marked differences in cellular Ru accumulation. After 24 h, mononuclear complexes **1** and **6** accumulated at 5.7–10.4 ng Ru per 10⁶ cells in HFF and HeLa cultures^21^, whereas trinuclear **7** reached 819 ± 14 ng Ru per 10⁶ HFF cells and 442 ± 15 ng Ru per 10⁶ HeLa cells (**Table S3**)^22^. Thus, enhanced uptake likely contributes to the higher activity of **7**, together with additional cellular effects examined below.

The activity of **1**, **6** and **7** extended to HSV-2, a major cause of genital herpes. All three compounds inhibited HSV-2 replication in a dose-dependent manner, with selectivity indices above 5 and reduced viral protein expression (**Extended Data Fig. 2**). We next tested HSV-1 variants resistant to acyclovir (ACV) or phosphonoformate (PFA), compared to our clinical HSV-1 isolate and the HSV-1 KOS reference strain (**Fig. 2d**). Compound **1** was particularly active in HFF cells against the clinical isolate, KOS and KOS-ACVr, whereas **7** showed broad activity across cell types, including against resistant variants in HFF and VERO cells. In HeLa cells, all three derivatives reduced replication of wild-type and resistant strains, with **6** showing the broadest efficacy. ACV and PFA lost activity against their corresponding resistant viruses, confirming the resistance phenotypes. Together, these data identify **1**, **6** and particularly **7** as Ru-imiquimod derivatives with activity against HSV-1, HSV-2 and drug-resistant HSV-1 variants.

Virucidal assays showed that **7** directly reduced HSV-1 infectivity at 37 °C, whereas no comparable effect was observed against non-enveloped adenovirus (**Extended Data Fig. 3a**), supporting an additional virus-directed component that may involve the viral envelope. The Ru-imiquimod complexes also bound and damaged DNA *in vitro*, with **7** showing the strongest effect (**Extended Data Fig. 3b,c**), although the contribution of this activity to antiviral efficacy remains unclear.

### Proteomic profiling reveals distinct host and viral responses to Ru–imiquimod treatment

We used label-free quantification proteomics to assess how HSV-1 infection and Ru-imiquimod treatment remodel the proteome of HFF^23^. Across 36 samples comprising vehicle-, **1**- and **7**-treated cells with or without HSV-1 infection, 5,078 proteins were identified, of which 4,327 were shared across all conditions, indicating robust measurements (**Fig. 3a**). Protein coverage was consistent across samples, with most proteins detected in all six biological replicates and strong within-condition correlations. Lower correlations between infected and uninfected samples reflected the presence of viral proteins (**Extended Data Fig. 4a–d**). Principal component analysis separated infected from non-infected cells and further distinguished **7**-treated samples from vehicle- and **1**-treated conditions, indicating distinct proteomic states induced by the viral infection and metallodrug treatment (**Fig. 3b**). HSV-1 infection was readily captured at the proteome level: 54 curated HSV-1 proteins were detected and hierarchically clustered across the three infected conditions (**Fig. 3c** and **Supplementary Table 1**). At 24 h post-infection, these included capsid proteins (UL18, UL35 and UL38), DNA-packaging factors (UL6, UL15, UL28 and UL33) and envelope glycoproteins (UL1, UL10, UL22, UL27, UL53 and US8). Viral protein intensities were broadly similar in vehicle- and **1**-treated infected cells, whereas **7** markedly reduced HSV-1 protein abundance.

**Figure 3:**
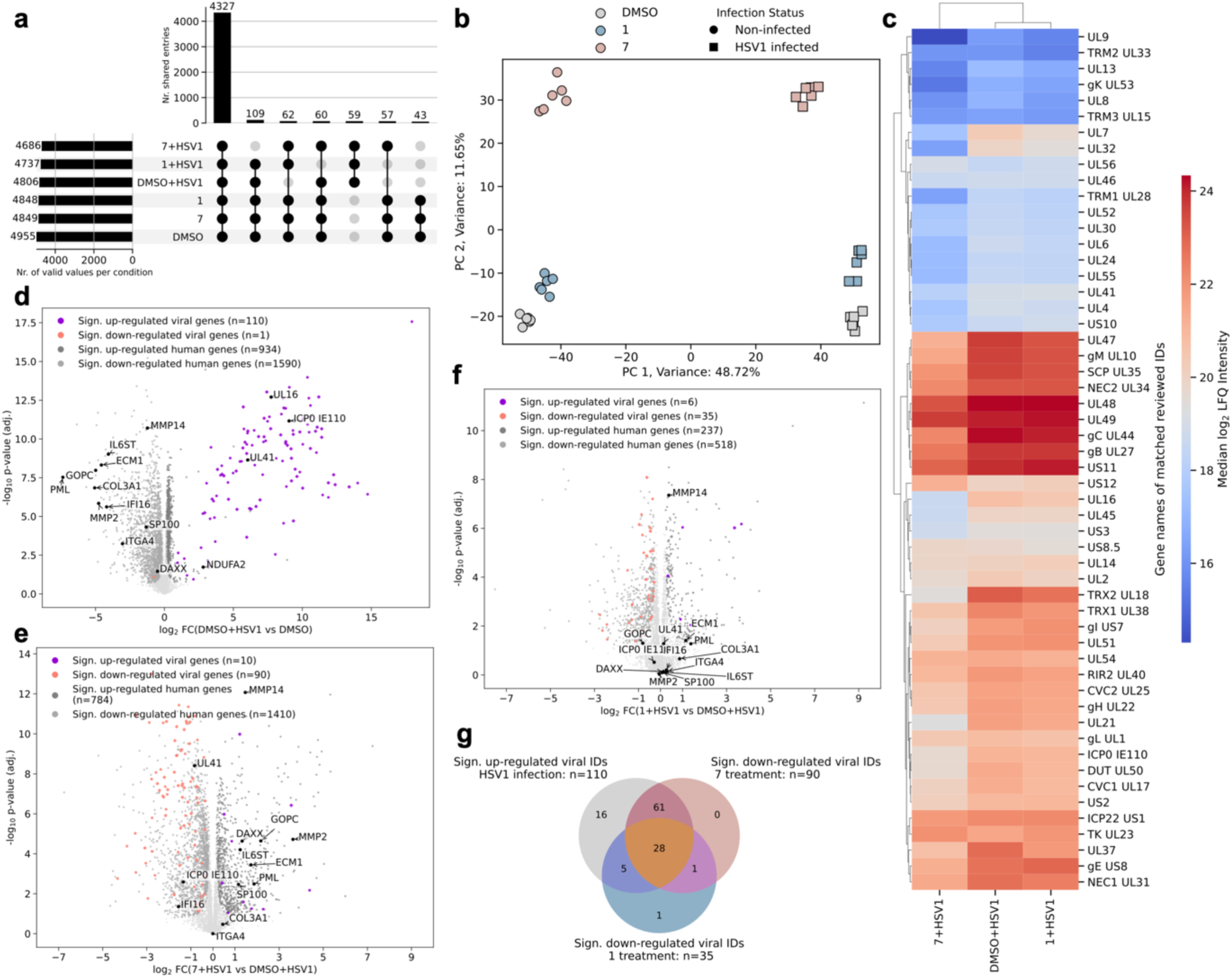
Compound 7 suppresses the HSV-1 proteome and remodels host antiviral responses. **a**, UpSet plot to visualise shared proteins across experimental conditions based on the total of 5078 identified proteins in the dataset. The majority of proteins is shared across all conditions. **b**, Principal component analysis (PCA) of the samples based on their proteome profile, highlighting the distinct impact of HSV-1 infection and treatment with **1** or **7**. **c**, Hierarchical clustering of the 54 reviewed HSV-1 proteins identified in the dataset and their intensity distribution across the experimental conditions. The associated colormap displays the median log_2_ LFQ intensity of the six biological replicates of each viral protein. **d**, Volcano plot highlighting the effects of HSV-1 infection in HFF. Purple dots correspond to up-regulated viral proteins above the significance threshold. Viral proteins represent the majority of regulated proteins in this comparison. The proteins of viral origin exhibit the highest fold changes of the proteins present in the dataset. Viral entries that were matched to Swiss-Prot reviewed proteins are shown as diamonds, while non-reviewed TrEMBL entries are shown as circles. Salmon colored dots depict significantly down-regulated viral proteins. **e** and **f** respectively represent the treatments of HSV-1 infected HFF cells with **7** and **1**. The color code employed in **e** and **f** is equal to **d**. **g**, Venn diagram highlighting the overlap of statistically significant viral protein regulations across HSV-1 infected conditions. HSV-1 infection = up-regulated viral proteins from DMSO+HSV-1 vs. DMSO; 7 treatment = down-regulated viral proteins from 7+HSV-1 *vs*. DMSO+HSV-1 and 1 treatment = down-regulated viral proteins from 1+HSV-1 vs. DMSO+HSV-1.

Differential proteome analysis of HSV-1-infected versus non-infected HFF-1 cells revealed a dual signature of viral hijacking and host defence (**Fig. 3d**). Infection was dominated by viral protein accumulation, consistent with extensive HSV-1 replication and the ability of the virus to redirect host metabolism towards virion production while suppressing host gene expression through mRNA degradation^24,25,26^. In parallel, infected cells activated antiviral defence programmes, including interferon-stimulated genes, host restriction factors and pathways linked to translational arrest^25^.

Two virus-related effects were evident in host cells. UL41, the virion host-shutoff protein that degrades host mRNA, was strongly upregulated^27^. Consistent with its reported effects on integrins, extracellular-matrix components and the adhesome^28^, several related proteins were downregulated, including ECM1, COL3A1, ITGA4, MMP2, MMP14, and IL6ST (**Fig. 3d**). Their reduced abundance may therefore reflect mRNA degradation by the viral nuclease. Second, ICP0, a tegument protein with E3-like activity, was strongly upregulated. ICP0 promotes the degradation of host restriction factors, including PML nuclear bodies and interferon-associated antiviral proteins^25,29^. Consistently, infected cells showed reduced levels of PML, SP100, IFI16, GOPC and, to a lesser extent, DAXX (**Fig. 3d**). GDF15 was also downregulated (**Extended Data Fig. 4e**); as this stress-response factor can be secreted, its reduced intracellular abundance may reflect release into the extracellular space^30^.

Beyond direct viral effects, regulated host proteins revealed defence mechanisms and extensive cellular remodelling. Fibroblasts typically induce a type I interferon response upon HSV-1 infection^24,31,32^, which promotes heterochromatin formation, mRNA degradation and translation arrest through eIF2α phosphorylation. Several components of this antiviral state were upregulated, including IFIT1–3 (**Extended Data Fig. 4f**) as well as ISG15, MX1 and the viral RNA sensor RIG-I/DDX58 (**Extended Data Fig. 4e**)^25,33^. Heat-shock and apoptosis-related proteins were also increased, indicating protective stress responses and apoptotic priming (**Extended Data Fig. 4j**).

Ru-imiquimod treatment reshaped both components of this response in a compound-dependent manner. During infection, **7** regulated 2,194 host proteins and reduced 90 viral proteins, compared with 755 host proteins and 35 viral proteins for **1** (**Fig. 3e,f**). Both compounds led to a significant down-regulation of 28 shared viral proteins, whereas **7** decreased further 61 unique proteins, consistent with its broader suppression of HSV-1 protein accumulation (**Fig. 3g**). Treatment with **7** also partially reversed infection-associated losses of host proteins targeted by the UL41 and ICP0 pathways: MMP2, MMP14, ECM1, DAXX, IL6ST, PML and SP100 were significantly up-regulated in infected cells treated with **7** compared with infected vehicle-treated cells (**Fig. 3e**). In addition, **7** induced NRF2-associated proteins, including HMOX1 and SRXN1, consistent with activation of a cellular redox capacity (**Extended Data Fig. 4g**).

In uninfected HFF, pre-ranked gene-set enrichment analysis showed that **7** primarily affected inflammatory-signalling and unfolded-protein-response pathways (**Extended Data Fig. 4h**). Notably, **7** also increased antiviral proteins including ISG15 and MX1 (**Extended Data Fig. 4i**), suggesting that the compound can modify the basal antiviral state of fibroblasts. EIF2AK2/PKR, the kinase that phosphorylates eIF2α, was also increased, consistent with a potential contribution of translational control to the reduction in viral protein synthesis^34^.

Targeted immunoblotting independently supported the proteomic signature of impaired HSV-1 protein accumulation. ICP4, VP16 and gD were selected to monitor progression from immediate-early regulation to late structural-protein production. Notably, **7** strongly reduced all three viral proteins, with gD becoming almost undetectable in each cell line examined. By contrast, **1** reduced these proteins only in HeLa cells, consistent with its narrower antiviral profile, whereas **6** produced a more modest decrease in HeLa cells (**Extended Data Fig. 5**).

Together, these data show that **7** acts at both viral and host-proteome levels: it broadly suppresses HSV-1 protein accumulation while reshaping host programmes associated with viral immune evasion, intrinsic antiviral restriction and cellular stress responses.

### Patient-derived ectocervical organoids reveal antiviral activity of Ru-imiquimod compounds

To assess antiviral activity in a physiologically relevant epithelial model, we used patient-derived human ectocervical organoids that reproduce the architecture and molecular organization of the stratified ectocervix^35,36^. The organoids retained CDH1-positive epithelial organization, KRT5- and P63-positive basal/parabasal compartments, and KRT6-positive suprabasal differentiated cells (**Extended Data Fig. 6a,b**). Compound tolerability was assessed by Hoechst and propidium iodide staining combined with automated fluorescence imaging. Imiquimod, **1** and **6** were well tolerated across the tested concentration range, whereas **7** reduced epithelial viability at concentrations above 50 μM (**Extended Data Fig. 6c,d**). These data defined non-cytotoxic concentrations for subsequent infection studies.

We next established a quantitative HSV-1-GFP infection assay using day-6 organoids infected with HSV-1 (17+) Lox-pMCMV-GFP and analysed 48 h later. Increasing multiplicities of infection produced progressively broader GFP-positive epithelial regions, whereas lower viral doses yielded graded, heterogeneous infection patterns (**Extended Data Fig. 6e**). An MOI of 0.006 was selected because it generated reproducible, non-saturating infection, enabling quantitative detection of treatment-dependent changes in HSV-GFP signal. Uninfected organoids showed minimal background fluorescence, and higher-magnification images revealed infection-associated disruption of epithelial boundaries and protrusion-like peripheral structures in GFP-positive regions (**Extended Data Fig. 6f**).

Using this model, organoids infected at MOI 0.006 were treated for 48 h with imiquimod, **1**, **6** or **7**. Representative images showed compound-dependent reductions in HSV-GFP infection, with the strongest effect observed with **7** (**Fig. 4a,b**). Quantification of GFP signal normalized to nuclear area confirmed a robust, concentration-dependent reduction by **7** at concentrations that preserved epithelial viability (**Fig. 4c**). Imiquimod, **1** and **6** produced partial inhibition at higher concentrations.

**Figure 4:**
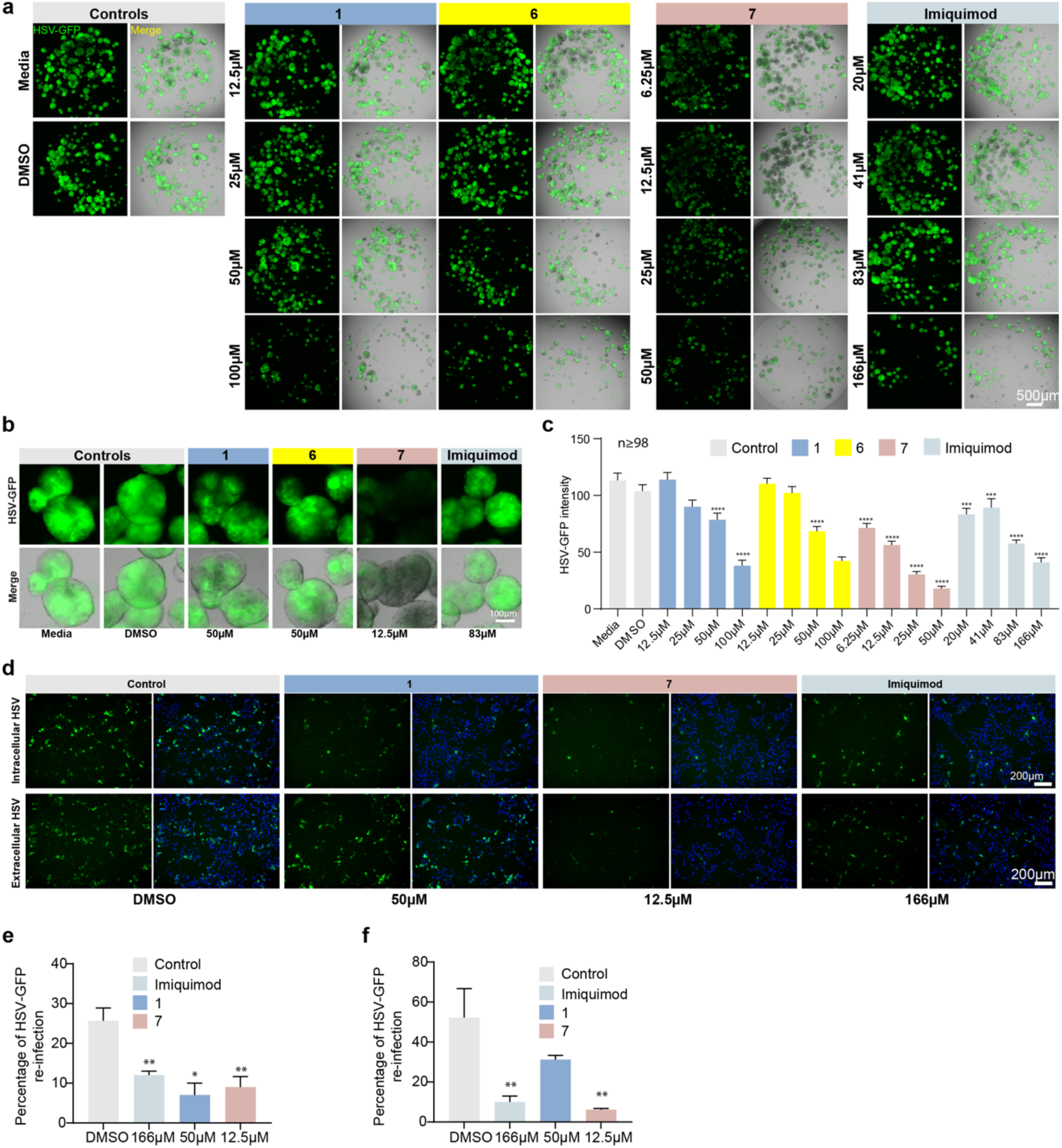
Compound 7 suppresses HSV-1 infection and reduces infectious virus recovery in patient-derived ectocervical organoids. **a**, Representative fluorescence and merged images of ectocervical organoids infected with HSV-1 (17+) Lox-pMCMV-GFP at MOI 0.006 and treated with the indicated concentrations of **1**, **6**, **7**, imiquimod, DMSO or medium control for 48 h. **b**, Higher-magnification images of selected conditions from panel a, showing compound-dependent changes in HSV-GFP signal and organoid morphology. **c**, Quantification of HSV-GFP signal normalized to nuclear area and plotted relative to DMSO control. Bars show mean ± SEM and statistical comparisons were performed using the Kruskal–Wallis test followed by Dunn’s multiple-comparison test against the media control. ***P = 0.0001; ****P < 0.0001. **d**, Representative fluorescence images of HeLa cells re-infected with cell-associated HSV-1 recovered from organoid lysates (top) or with released HSV-1 recovered from organoid culture supernatants (bottom), after treatment of organoids for 48 h with DMSO, **1**, **7** or imiquimod at the indicated concentrations. GFP marks HSV-1-infected cells; nuclei were counterstained with DAPI. Scale bars, 200 µm. **e–f**. Quantification of HSV-GFP-positive HeLa cells re-infected with cell-associated virus from organoid lysates (**e**) and with released virus from organoid supernatants (**f**). Data are presented as mean ± s.d.; statistical significance was assessed using an unpaired two-tailed Welch’s t-test against the DMSO control. *P < 0.05; **P < 0.01. Representative images in **a** and **d**, and data in **c**, **e**, and **f**, are derived from three independent biological replicates.

We then tested whether reduced epithelial infection was accompanied by reduced production of infectious virus. Cell-associated virus from organoid lysates and released virus from culture supernatants were transferred to permissive HeLa cells as secondary infection readouts (**Fig. 4d-f**). Lysates from **1**-, **7**-, and imiquimod-treated organoids generated fewer HSV-GFP-positive HeLa cells than vehicle-treated controls (**Fig. 4d**, upper panels, **Fig. 4e**). Supernatants from all three treatment groups similarly showed reduced infectivity, with **7** producing the strongest effect despite being used at a lower concentration than imiquimod and **1** (**Fig. 4d**, lower panels, **Fig. 4f**).

Together, these findings show that **7** retains strong antiviral activity in a patient-derived stratified epithelial model, reducing both the extent of epithelial infection and the recovery of infectious virus. The organoids therefore provide a physiologically relevant human epithelial model in which to examine how Ru-imiquimod treatment influences not only viral burden but also the epithelial response to infection.

### Compound 7 selectively restores antiviral programmes in HSV-1-infected cervical epithelium

To distinguish direct antiviral activity from modulation of epithelial responses, we profiled patient-derived ectocervical organoids treated with vehicle (DMSO), imiquimod, **1** or **7**, with or without HSV-1 infection. Microarray analysis of eight groups (n = 5) enabled independent assessment of treatment, infection, and drug-infection interaction effects (**Fig. 5a**). After background correction and normalization, expression profiles were comparable and showed no systematic batch effects (**Extended Data Fig. 7a**). Principal component analysis separated samples by infection status and treatment, with tight clustering of replicates, confirming robust and reproducible responses (**Fig. 5b**).

**Figure 5:**
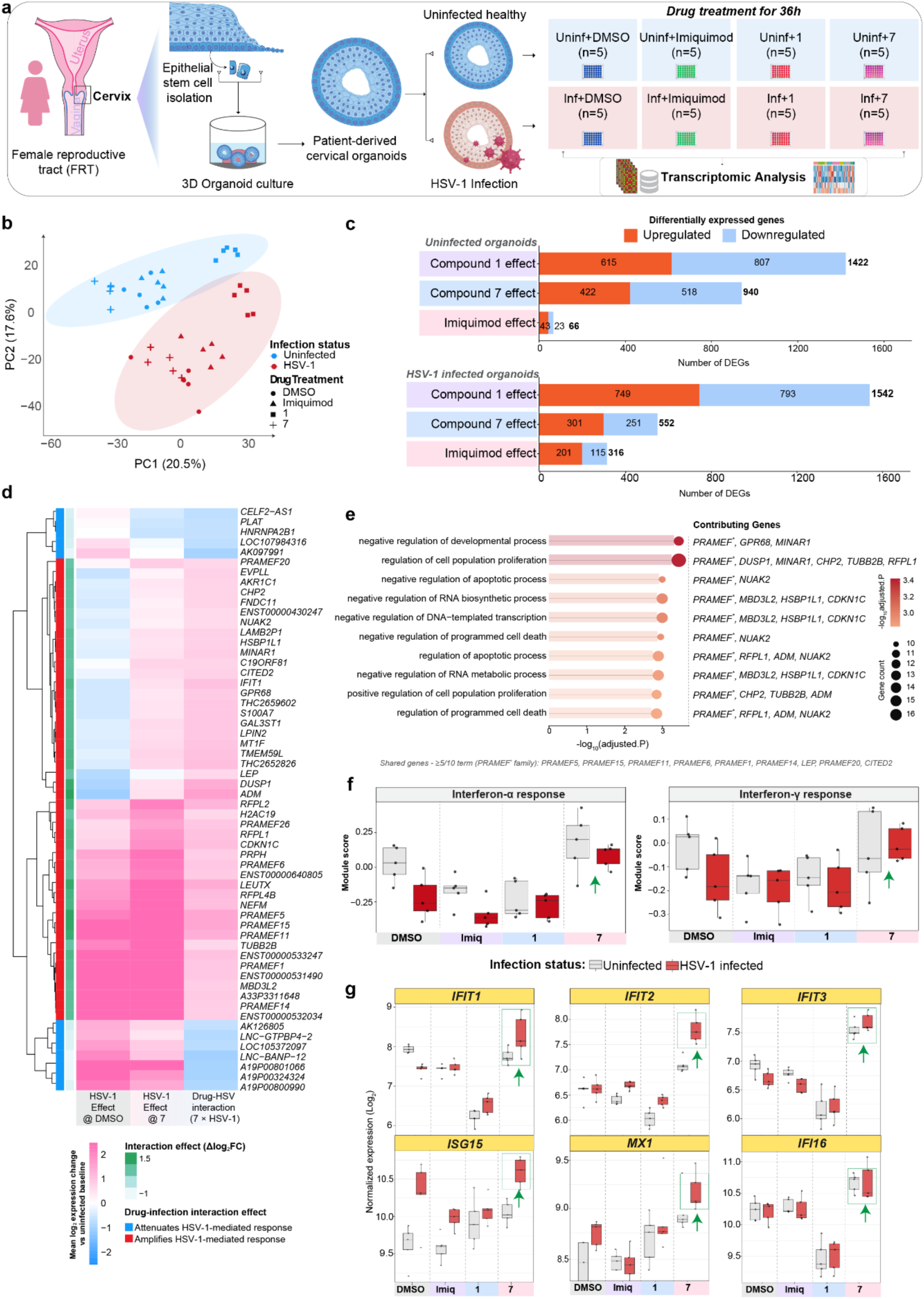
Compound 7 selectively restores antiviral programmes in HSV-1-infected cervical epithelium. **a**, Schematic overview of experimental design. Patient-derived cervical organoids were treated with DMSO, Imiquimod, **1**, or **7** for 36 hours in the presence or absence of HSV-1 infection (n = 5 per group) and profiled by microarray. **b**, Principal component analysis (PCA) of the 2,000 most variable genes across all samples. Points are coloured by infection status and shaped by treatment. Variance explained by each principal component is indicated on the respective axis. **c**, Bar plot showing the total number of differentially expressed genes (DEGs) for each compound relative to DMSO in uninfected (top) and HSV-1-infected (bottom) organoids (Benjamini–Hochberg-adjusted P < 0.05, log2FC > 0.6), separated by direction. **d**, Heatmap showing genes whose HSV-1-induced expression is modified by compound **7**. Columns show the HSV-1 effect in the vehicle background, the HSV-1 effect in the **7** background, and the **7** x HSV-1 interaction, defined as the difference between the two. The main color scale shows the mean log2 expression change relative to the uninfected baseline; the adjacent scale shows the interaction effect (Δlog2FC), and the leftmost annotation indicates whether **7** amplifies or attenuates the HSV-1-mediated response. **e**, GO biological-process terms enriched among genes with amplified HSV-1 responses under **7** (over-representation analysis; adjusted P < 0.05). Bar length and color show -log10(adjusted P), point size the number of contributing genes. Representative genes are listed for each term; genes shared by at least five of the ten terms are given below the panel. **f**, Hallmark interferon-α and interferon-γ module scores across all conditions, calculated as the mean z-scored expression of gene-set members per sample. **g**, Box plots showing the normalized expression (log2) of interferon-stimulated genes across all conditions. In **f** and **g**, box color indicates infection status (grey: uninfected; red: HSV-1 infected).

In uninfected organoids, each compound induced a distinct transcriptional baseline. Compound **1** regulated 1,422 genes, **7** regulated 940 and imiquimod only 66 (**Fig. 5c**, **Supplementary Data 1**). The responses were largely non-overlapping, with moderate concordance between imiquimod and either Ru complex but weak concordance between **1** and **7** (**Extended Data Fig. 7b–d**), indicating that the Ru complexes generated distinct regulatory states rather than simply amplifying imiquimod activity. These baseline responses were also functionally distinct. Compound **1** broadly altered metabolic, structural and signalling genes, including *HOTAIRM1*, *GDF15* and *ALDH3A1*, while suppressing innate immune and defence programmes. Compound **7** increased *H19*, *CXCL17* and *TRIM22* and reduced *TUBB3*, *MMP1*, *IL1A* and *IL1B* (**Extended Data Fig. 7e–g**, **Supplementary Data 1**). Enrichment analyses linked **1** to lipid and organic-acid metabolism and KRAS suppression, whereas **7** activated DNA-replication, cell-cycle and MYC-target programmes while suppressing hypoxia, TNF-NF-κB and inflammatory signatures (**Extended Data Fig. 7h,i**). Imiquimod produced a narrower response associated with interferon suppression, establishing distinct baselines for interpreting infection-dependent effects.

HSV-1 infection substantially reshaped the magnitude and composition of these responses. Within infected organoids, **1** again produced the largest transcriptional perturbation, regulating 1,542 genes, compared with 552 for **7** and 316 for imiquimod (**Fig. 5c**, **Supplementary Data 1**). However, the regulated gene sets remained largely compound-specific, with minimal overlap and low Jaccard similarity across treatments (**Extended Data Fig. 8a–c**). Thus, transcriptional breadth did not correlate directly with antiviral activity: **1** altered the largest number of genes, whereas **7** produced a more selective antiviral response.

In infected organoids, **1** broadly suppressed epithelial differentiation, keratinization, ribosome biogenesis and interferon-associated programmes. Compound **7** more selectively reduced epithelial differentiation and proliferation while inducing stress-adaptive and antiviral genes, including the E3 ubiquitin ligases *TRIM22*, *HERC5*, *TRIM31* and *TRIM49* (**Extended Data Fig. 8d–h**, **Supplementary Data 2**). This enrichment was not observed in uninfected organoids, indicating that these antiviral restriction programmes were preferentially engaged by **7** in the context of HSV-1 infection. Hallmark analysis further linked **7** to suppression of E2F, G2/M-checkpoint and mitotic-spindle programmes, whereas **1** suppressed MYC targets and imiquimod mainly affected estrogen-response pathways (**Extended Data Fig. 8i**, **Supplementary Data 2**).

We next defined the core HSV-1 transcriptional programme to distinguish virus-driven changes from drug-specific effects. In vehicle-treated organoids, infection regulated 758 genes, predominantly through induction, and 343 genes were consistently altered across all treatment backgrounds, defining a conserved epithelial infection signature (**Extended Data Fig. 9a-c**). This signature combined induction of stress- and antiviral-associated genes, including *TRIM43*, *TRIM49*, *PRAMEF*-family members and *RASD1*^37^, with suppression of metabolic, proliferative and chromatin-related processes (**Extended Data Fig. 9d-h**, **Supplementary Data 1**,**2**). Selected markers showed how this conserved response was modified by treatment. *TRIM49* and *RASD1* were induced across all backgrounds^38,39^, whereas *NQO1* remained high only under **1**, indicating compound-specific antioxidant engagement (**Extended Data Fig. 9i**)^40^. All three treatments increased *SQSTM1* in infected organoids^41^, while **7** partially restored the HSV-suppressed restriction factor *SP100*, with a weaker effect from imiquimod^42^. *IFI16* and *TBK1* remained comparatively stable, whereas *IRF7* was modestly reduced, consistent with viral attenuation of interferon amplification (**Extended Data Fig. 9j**)^43^. Thus, this conserved HSV-1 signature provided the reference needed to identify virus-induced changes selectively reversed or reinforced by each compound.

We next used drug × HSV-1 interaction analysis to identify transcriptional responses that could not be explained by the independent effects of drug treatment or infection alone. Here, the interaction quantifies how the HSV-1-induced expression change differs between a drug-treated background and the corresponding DMSO background. Thus, a positive interaction indicates that the HSV-1-associated expression change becomes more positive under drug treatment, whereas a negative interaction indicates a shift in the opposite direction. Further, the biological interpretation depends on the direction of the underlying HSV-1 response. Imiquimod produced a limited interaction profile, whereas **1** induced a broader but heterogeneous attenuation of selected HSV-responsive genes (**Extended Data Fig. 10a**,**b**). Compound **7** generated a more coherent interaction response, enhancing infection-dependent induction of PRAMEF-family genes, *LEUTX*, *NUAK2*, *CITED2*, *RFPL1* and *CDKN1C*, with enrichment in transcriptional regulation, RNA biosynthesis, developmental control and programmed cell death (**Fig. 5d,e** and **Extended Data Fig. 10c, Supplementary Data 1, 2**).

Most notably, **7** restored HSV-suppressed interferon-α and interferon-γ programmes, with module scores reaching or exceeding uninfected levels, whereas **1** produced the weakest responses (**Fig. 5f** and **Extended Data Fig. 10d**). Consistently, **7** increased *IFIT1-3*, *ISG15*, *MX1* and *IFI16*, particularly *IFIT2*, *ISG15* and *MX1*, while **1** maintained low IFIT expression (**Fig. 5g**). Increased *IFI16* transcription may partially compensate for its ICP0-mediated degradation^44^. These findings indicate that **7** does not simply increase transcriptional activity globally, but preferentially re-establishes antiviral programmes suppressed during HSV-1 infection. Compound **7** also reduced proliferation-associated transcripts, including *MKI67*, *TOP2A*, *CCNB1* and *CDK1*, whereas imiquimod and **1** maintained or increased them (**Extended Data Fig. 10e**). *GMNN* displayed a distinct pattern, suggesting that the effects of compound **7** do not represent uniform suppression of all cell-cycle-associated genes^45,46^.

Regulon-based transcription factor (TF) analysis provided a complementary view of the regulatory programmes underlying these divergent responses. HSV-driven contrasts clustered across drug backgrounds, confirming a conserved infection programme, whereas drug-only and interaction contrasts formed a separate, compound-dependent group (**Fig. 6a**, **Supplementary Data 3**). Interaction signatures were not simple sums of the independent drug and infection effects, indicating emergent regulation in infected cells.

**Figure 6:**
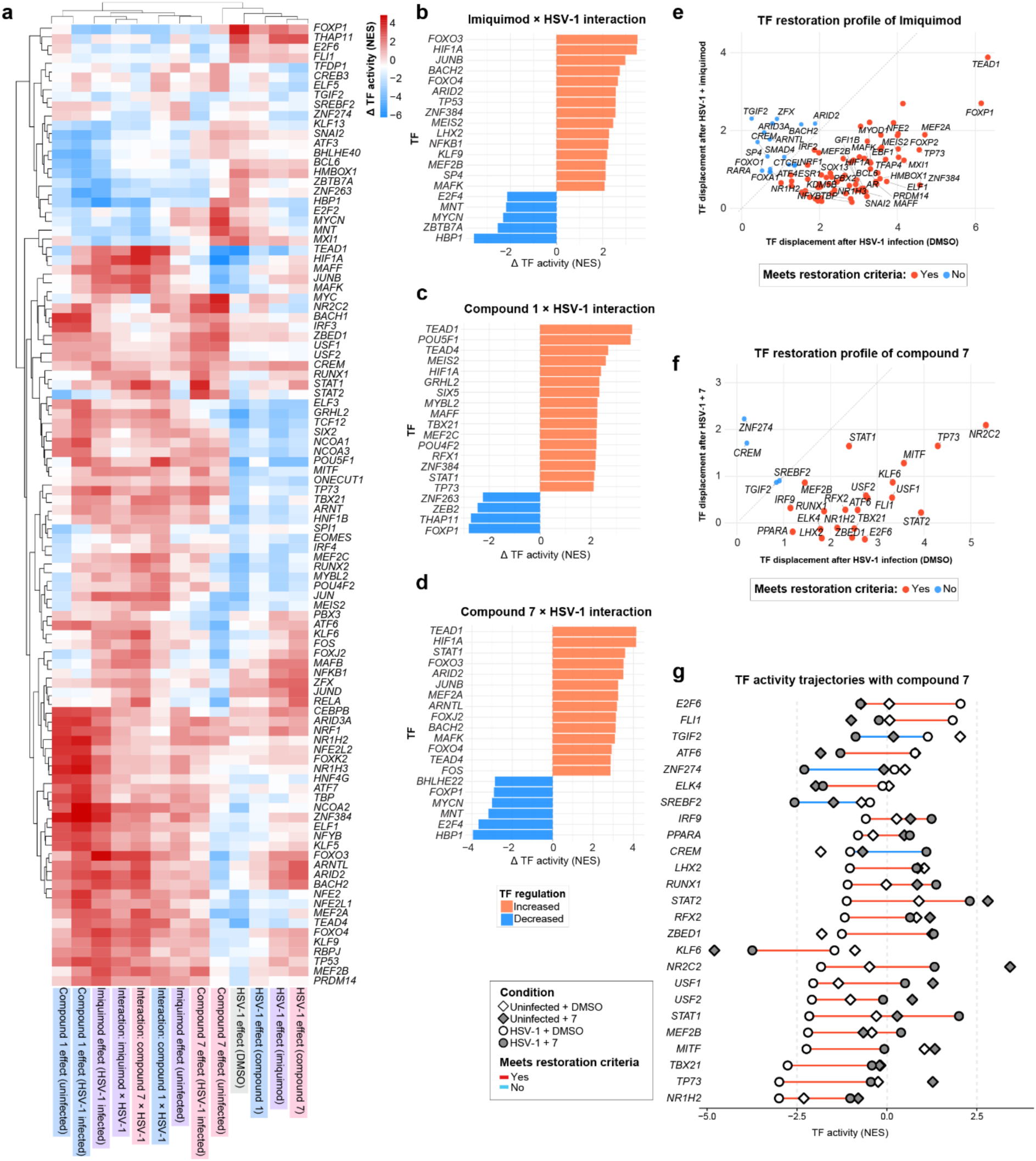
Regulatory reprogramming under compound 7 defines the infection-adapted antiviral state. **a**, Heatmap of inferred transcription factor (TF) activity changes for the top differentially active TFs across key contrasts. Rows represent individual TFs; columns represent contrasts ordered by biological context. Color indicates direction and magnitude of TF activity (Δ normalized enrichment score, NES) change. TFs shown are the top 20 with the largest absolute differential activity among significant TFs (adjusted P < 0.05) in each displayed contrast. **b–d**, Bar plots showing the top strongly altered TFs in the drug x HSV-1 interaction contrast conditions for Imiquimod (**b**), **1** (**c**), and **7** (**d**), ranked by absolute Δ TF activity. Positive values (orange bars) indicate increased TF activity, whereas negative values (blue bars) indicate decreased activity. **e–f**, Scatter plots highlighting the TF activity changes under HSV infection for Imiquimod (**e**) and **7** (**f**). Each dot represents a TF, positioned by its absolute displacement from the drug-matched uninfected baseline under DMSO-treated infection (x-axis; HSV_DMSO - Uninfected_Drug) versus drug-treated infection (y-axis; HSV_Drug - Uninfected_Drug). Dots below the diagonal indicate TF activity restored toward the drug-specific uninfected baseline. Color indicates whether the movement met the restoration criteria (reduction in displacement > 0.3 NES and > 20% of the infection-induced displacement), to distinguish substantial from small shifts. **g**, Strip plot showing distribution of TF activity changes for **7**-treated HSV-1 infected organoids. For each TF, the line connects the HSV-1 + DMSO and HSV-1 + **7** conditions and is colored by whether the restoration criteria were met; symbols mark the four experimental conditions. TFs represent those with the strongest interaction effects; values are limma group means.

In vehicle-treated organoids, HSV-1 suppressed regulators of epithelial identity and differentiation, including *GRHL2*, *TEAD1*, *POU5F1* and *ARNT*, while activating selected stress-responsive factors. Similar patterns were observed with imiquimod and **1**, although **1** caused broader transcription-factor suppression. By contrast, HSV-1 infection in the presence of **7** was associated with increased activity of a coherent regulatory network comprising *ARNTL*, *ARID2*, *CEBPB*, *RELA*, *FOS*, *JUNB* and *BACH2* (**Fig. 6a**, HSV-1 effect columns)^47,48,49,50^. Several of these factors are linked to antiviral defence, stress adaptation and control of inflammatory responses, suggesting that **7** engages a regulatory state distinct from simple inflammatory activation.

Drug × HSV-1 interaction analysis further distinguished the compounds. Imiquimod activated few stress-responsive regulators, including *HIF1A*, *JUNB* and *FOXO3*, whereas **1** activated epithelial-identity factors and *STAT1* without restoring downstream interferon effectors (**Fig. 6b,c**). Compound **7** showed the broadest inferred antiviral regulatory signature, with increased *STAT1*, *STAT2*, *HIF1A*, *ARNTL*, *JUNB* and *BACH2* activity (**Fig. 6a,d**, **Supplementary Data 3**). These factors encompass canonical interferon signalling together with stress-adaptive and homeostatic regulatory programmes. We then asked whether treatment returned infection-displaced regulators toward the uninfected state. Restoration was measured relative to each compound’s own uninfected baseline, ensuring that compound-induced changes in uninfected cells were not themselves interpreted as restoration. **7** restored several HSV-displaced regulators, including *STAT1*, *STAT2*, *TBX21*, *RUNX1* and *FLI1*, towards their uninfected state (**Fig. 6e-g**)^51^. Imiquimod showed limited restoration, whereas **1** combined NRF2-associated stress responses with persistent interferon attenuation (**Extended Data Fig. 10f-h**, **Supplementary Data 3**) ^38,40^.

Thus, transcriptional breadth did not predict antiviral efficacy. Compound **1** produced the largest overall perturbation but broadly suppressed interferon-associated programmes, whereas imiquimod had comparatively limited effects. Compound **7**, despite regulating fewer genes, generated a more coherent infection-dependent antiviral response, restoring interferon-responsive programmes, increasing antiviral restriction factors and re-establishing STAT1- and STAT2-centred regulatory networks displaced by HSV-1 infection. Rather than broadly amplifying epithelial transcription, **7** therefore appears to recalibrate the infected epithelial state towards antiviral competence. These findings provide a mechanistic basis for its distinct antiviral activity in patient-derived cervical epithelium.

## Discussion

Here we show that direct coordination of imiquimod to Ru(II)-arene fragments reprogrammes the pharmacology of an established immune-active scaffold. The resulting complexes did not function as progressively stronger versions of imiquimod; instead, they acquired distinct chemical, antiviral, and host-response properties. This effect was most pronounced for the trinuclear complex **7**, which exhibited broad activity against HSV-1 and HSV-2, suppressed productive infection, and elicited a distinctive response in patient-derived cervical epithelium. Therefore, Ru coordination altered not only antiviral potency but also the manner in which the biological activity of imiquimod was manifested.

The observed diversity across the Ru-imiquimod series supports this conclusion. Complexes **1**, **6**, and **7** differed in solution behaviour, cellular accumulation, cell-type dependence, and antiviral activity. Notably, complexes **1** and **7** induced markedly different transcriptional states despite sharing the same immune-active ligand. The relatively high stability and cellular accumulation of complex **7** likely contribute to its distinct behaviour, although its trinuclear architecture prevents a direct molecule-for-molecule comparison of total Ru accumulation with the mononuclear complexes. More broadly, these differences indicate that coordination architecture is an essential determinant of biological response. This observation is consistent with established organoruthenium pharmacology, where coordination environment, ligand exchange, and activation state can substantially alter cellular engagement and biological activity^52,53^.

The comparison between complexes **1** and **7** provides clear evidence that the observed effects are not solely quantitative. Compound **1** induced the largest transcriptional perturbation in both uninfected and infected organoids, yet exhibited weaker antiviral activity and persistent attenuation of interferon-associated programmes. In contrast, **7** regulated fewer genes but generated a more coherent antiviral response in the infected state. Thus, transcriptional breadth did not predict antiviral efficacy; rather, the identity and infection dependence of the engaged programmes were more informative. This distinction is particularly relevant for immune-active scaffolds such as imiquimod, where stronger transcriptional activation does not necessarily result in more effective antiviral defence. Imiquimod can inhibit HSV-1 independently of canonical TLR7 signalling through an adenosine A1 receptor–PKA–cystatin A pathway^54^, further demonstrating that its antiviral and canonical immunostimulatory activities are separable. Therefore, the phenotype of complex **7** cannot be explained by simple amplification of native imiquimod signalling.

The organoid data highlight the biological significance of this distinction in human epithelium. Patient-derived cervical organoids retain regional epithelial identities and reveal infection-associated cellular programmes that are challenging to resolve in conventional monolayer cultures^35,55^. Single-cell profiling has further demonstrated that cervical epithelial defence is heterogeneous and distributed across both infected and bystander populations^36^. In this study, complex **7** did not broadly amplify epithelial transcription. Its most pronounced effects emerged during HSV-1 infection, where interferon-α and interferon-γ response programmes were restored, along with *IFIT1-3*, *ISG15*, *MX1*, *IFI16*, and *TRIM*-family restriction factors. These responses define an infection-adapted antiviral state rather than generalized immune stimulation.

This infection dependence is significant because HSV-1 does not globally suppress innate immunity. Instead, it selectively remodels interferon signalling and intrinsic restriction, while antiviral, inflammatory, and stress programmes remain active. HSV-mediated host shutoff and immune-evasion mechanisms disrupt components of JAK–STAT signalling, including JAK1 and STAT2^56^, as well as other intrinsic defence pathways. Restoration of interferon-responsive programmes by complex **7** is therefore consistent with counteracting virus-associated attenuation of antiviral defence. The inferred regulatory landscape supports this interpretation: *STAT1*- and *STAT2*-centred activity was re-established, accompanied by changes in HIF1A, *ARNTL*, *JUNB*, and *BACH2*, and by movement of several regulatory states toward the uninfected condition. Rather than implicating a single transcription factor as the mechanism of action, the convergence of interferon-module restoration, antiviral effector induction, and regulon activity indicates a coherent reorganization of the infected epithelial state.

Proteomic data provide an independent and complementary perspective on this pharmacology. In HFF fibroblasts, **7** broadly reduced HSV-1 protein accumulation across multiple stages of the viral programme while remodelling host proteins associated with intrinsic restriction, stress responses, and viral immune evasion. Although proteomics and organoid transcriptomics interrogate different molecular layers and cellular contexts, both approaches converge on a shared biological interpretation: productive HSV-1 infection is suppressed while host antiviral programmes are reorganized. Such pathway-level convergence is characteristic of contemporary virus–host multi-omics studies, where transcriptomic, proteomic, phosphoproteomic, and genetic measurements capture complementary dimensions of infection rather than requiring molecule-for-molecule agreement^57^.

The pharmacological activity of **7** extends beyond host-response modulation. Activity was retained against clinical HSV-1 strains and variants resistant to established antivirals, remained evident when treatment was initiated after viral adsorption, and included a direct reduction in HSV-1 infectivity in virucidal assays at 37 °C. These observations exclude host pre-activation as the sole basis of efficacy and establish a complementary virus-directed component. The distinction of **7** therefore lies in the coexistence of both activities within the same coordination scaffold.

Functional studies in cervical organoids link these molecular responses to infection control. Compound **7** produced the most pronounced concentration-dependent reduction in HSV infection and decreased recovery of infectious virus from both organoid lysates and supernatants. The antiviral and regulatory phenotypes of complex **7** are therefore preserved in organized human cervical epithelium, establishing a tissue-level connection between the infection-dependent host response and suppression of productive HSV infection.

Collectively, these findings show a principle that extends beyond the specific Ru-imiquimod complexes examined in this study. The key advance is not merely the generation of a more active imiquimod derivative, but the demonstration that coordination chemistry can redirect the response profile of an immune-active scaffold. In complex **7**, this results in a pharmacological profile characterised by strong restriction of productive HSV-1 infection, selective restoration of antiviral programmes in infected epithelium, and an additional virus-directed component. The comparison with complex **1** further demonstrates that the magnitude of host perturbation does not predict antiviral efficacy.

More broadly, this work expands the role of coordination chemistry in anti-infective drug design. Metal-based compounds occupy a chemical space in which geometry, ligand environment, exchange kinetics, and reactivity can be manipulated to alter cellular engagement, and their biological activities increasingly extend beyond classical DNA-targeting paradigms^58^. These findings demonstrate that such chemical flexibility can be applied to an immune-active scaffold to reshape antiviral pharmacology. Coordination chemistry may therefore offer a means not only to alter potency or target engagement, but also to control how antiviral activity is configured across both virus and infected tissue.

## Acknowledgements

The authors thank Nanna Søndberg Lindberg Breum Nielsen for her valuable technical assistance with the antiviral experiments and Ina Wagner for support with the microarray experiments. SGIKER (UPV/EHU/ERDF, EU) is acknowledged for technical and human support. S.M.M-M. thanks Hans Paper for assistance in data interpretation. The authors are grateful to Core Facility for Mass Spectrometry and the Joint Metabolome Facility (University of Vienna and Medical University of Vienna), which are members of the Vienna Life-Science Instruments (VSLI).

## Author contributions

L.S. conceived the design of Ru-imiquimod complexes and the overall metal-based antiviral strategy. L.S., V.D.O., C.C., S.M.M.-M., and R.K.G. contributed to the overall study design. A.C.C., M.J.M., R.M., and M.A.H. carried out the synthesis, characterization, and reactivity studies of the complexes (Fig. 1 and the associated Supplementary Information). V.D.O., G.B. and M.B. performed the antiviral studies in 2D cell models and the immunoblotting experiments (Figs. 2, 3 and the associated Supplementary Information and Data). T.I., A.B., C.G. and S.M.M.-M. performed the proteomics studies (Fig. 3 and the associated Supplementary Information and Data). T.I. and A.B. performed data curation. T.I. and S.M.M.-M performed data interpretation on the proteomics data.

C.C. and R.K.G. conceived and designed the organoid and transcriptomic studies presented in Figs. 4–6, Extended Data Figs. 6–10, and the associated Supplementary Information and Data. N.K. contributed to experimental design and performed the organoid experiments, including cytotoxicity, HSV-1 infection, infectious-virus recovery, and RNA preparation for microarray analysis. P.G.P., together with C.C. and R.K.G., designed the transcriptomic analysis strategy, and P.G.P. performed the bioinformatic analyses. P.G.P., R.K.G. and C.C. interpreted the transcriptomic and drug–infection interaction analyses. H.-J.M. performed the microarray processing.

C.C., S.M.M.-M., M.A.H., V.D.O. and L.S. provided funding and resources and supervised the work. L.S. and C.C. wrote the manuscript. P.G.P., N.K., R.K.G., V.D.O. and S.M.M.-M. contributed to manuscript writing. R.K.G., C.C., V.D.O., S.M.M.-M., M.A.H. and L.S. critically revised the manuscript. All authors reviewed and approved the final version of the manuscript.

## Data availability

Chemical synthesis, compound characterization and associated analytical data are provided in the Supplementary Information. Crystallographic data for complexes **1**, **2**, **4**, **6** and **7** have been deposited with the Cambridge Crystallographic Data Centre (CCDC) under deposition numbers 2583749–2583753. The mass spectrometry proteomics data have been deposited to the ProteomeXchange Consortium via the PRIDE partner repository with the dataset identifier PXD086827. Microarray data from this publication have been deposited in the National Center for Biotechnology Information Gene Expression Omnibus (GEO) under accession code GSE352174.

Source data underlying the main figures, Extended Data figures and Supporting Information figures are provided with this paper. Additional data supporting the findings of this study are available from the corresponding authors upon reasonable request.

## Extended Data

**Extended Data Fig. 1:**
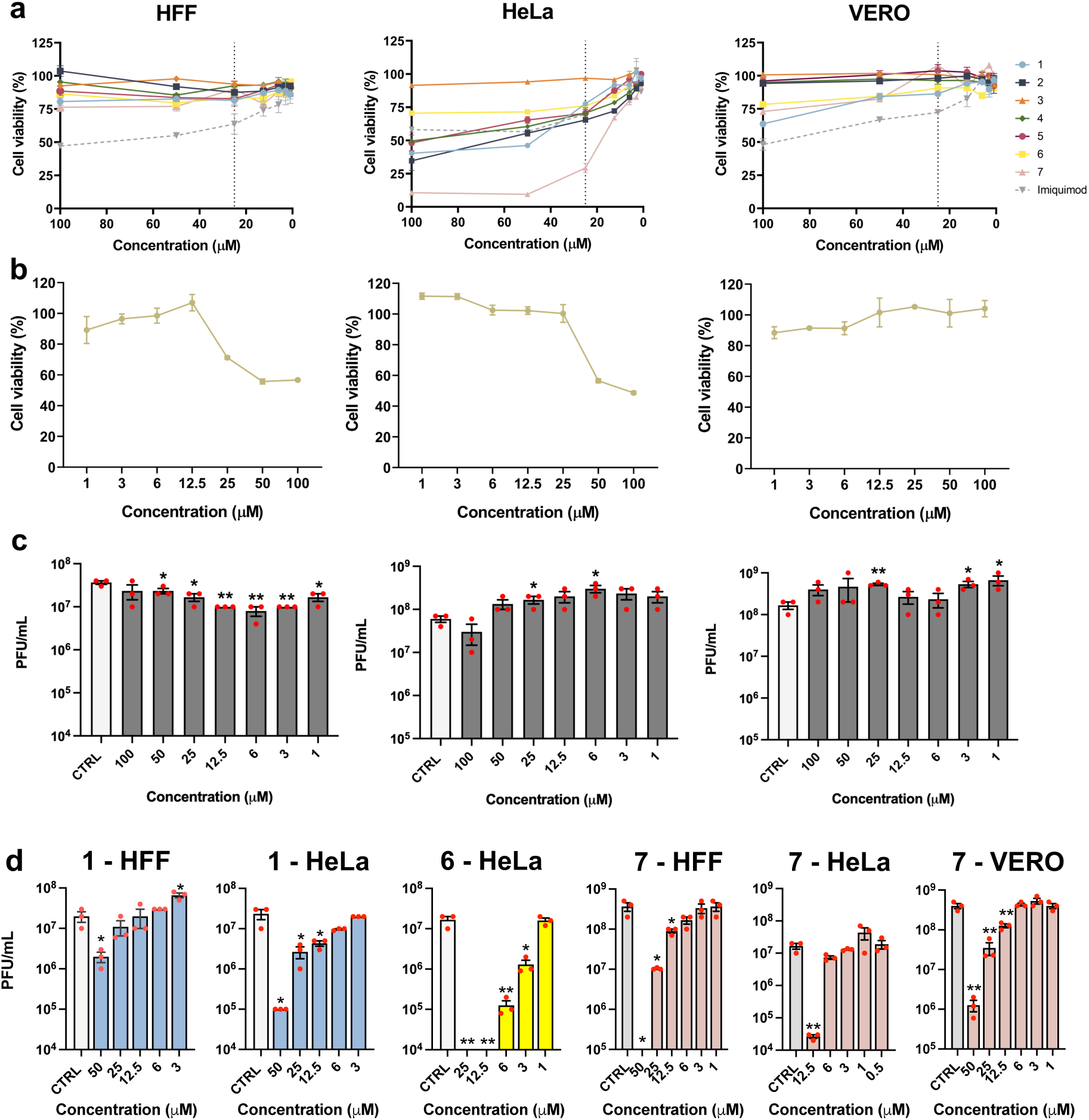
Cytotoxicity and antiviral activity of Ru-imiquimod compounds and the Ru-p-cymene precursor. **a,b**, Cell viability of confluent HFF, HeLa and VERO cultures treated with increasing concentrations of the indicated Ru-imiquimod compounds (**a**) or the Ru-p-cymene precursor (Ru-p-cym, **b**), or vehicle control (CTRL). Viability was assessed by MTT assay after 48 h. **c**, Antiviral activity of Ru-p-cym against HSV-1 in HFF, HeLa and VERO cells. Cells were infected with HSV-1 (MOI 0.1) and treated with the indicated concentrations of Ru-p-cym or CTRL. Viral yield was determined by plaque assay on VERO cells using combined cell-associated and supernatant fractions. **d**, Concentration-dependent antiviral activity of **1**, **6** and **7** in the indicated cell lines following HSV-1 infection (MOI 0.1, 48 h). Viral yield was quantified by plaque assay on VERO cells using combined cell-associated and supernatant fractions. Data represent mean ± SEM from three independent experiments and are expressed as cell viability (**a**,**b**), PFU/mL (**c**,**d**). Statistical significance was determined by unpaired *t*-test versus CTRL: *P* < 0.05 (*), *P* < 0.01 (**).

**Extended Data Fig. 2:**
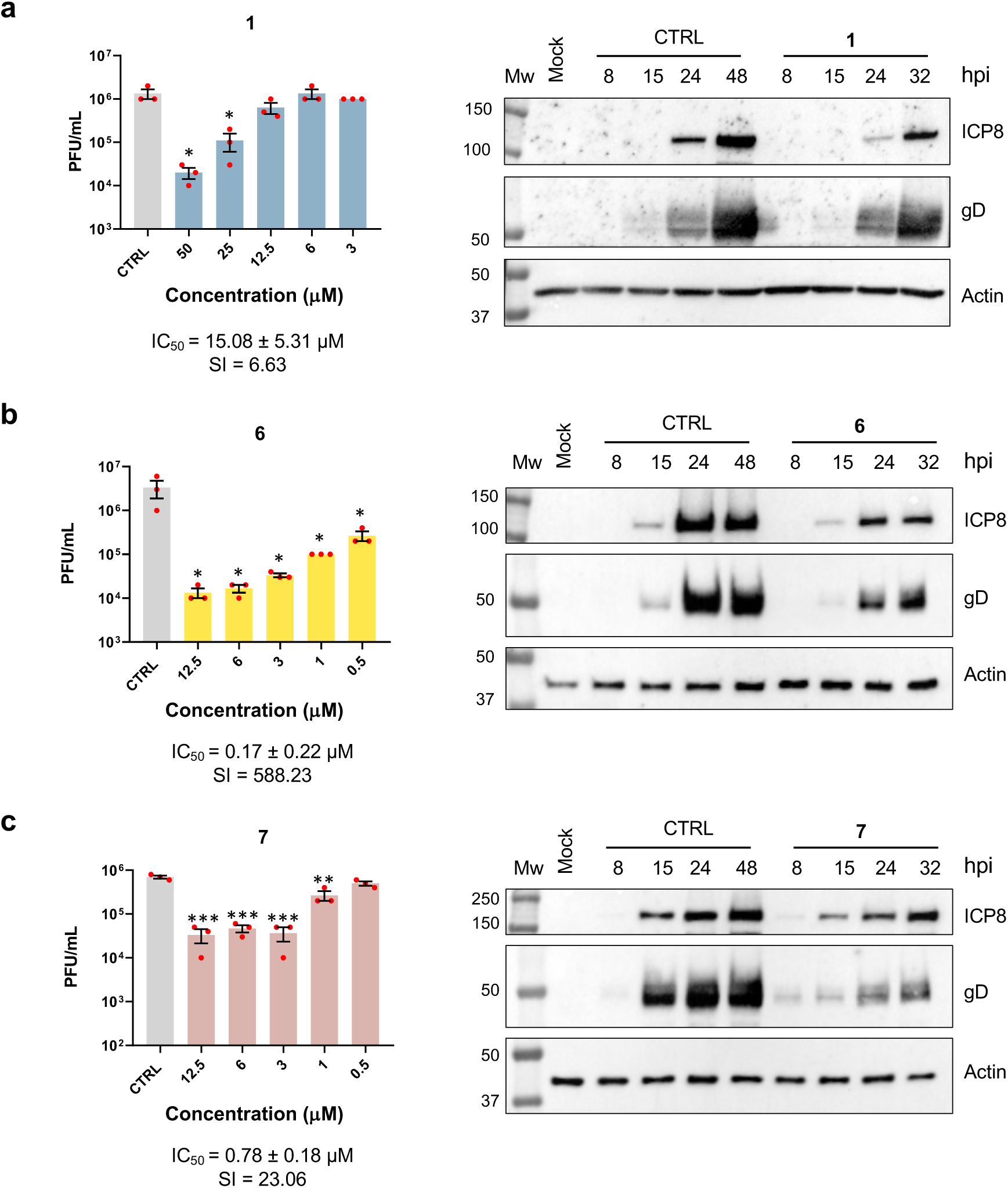
Antiviral activity of lead compounds 1, 6 and 7 against HSV-2. Left panels. Confluent HeLa cells were pre-treated with varying concentrations of **1** (**a**), **6** (**b**), **7** (**c**) or an equivalent volume of CTRL, infected with HSV-2 (MOI = 0.1) for 2 h at 37°C in the presence of the compounds. The inoculum was then removed and replaced by fresh compound for an additional 48 h. The extent of HSV-2 replication was then quantified by titrating the infectivity of supernatants and cell-associated viruses using a standard plaque assay on VERO cells. Plaques from three independent experiments were microscopically counted, and results are expressed as mean plaque-forming units per milliliter (PFU/mL) ± SEM for the entire process (p < 0.05 *, p < 0.01 **, p < 0.001 *** unpaired t-test compound dilutions *vs.* CTRL). Right panels. Effect of **1** (25 μM) (**a**), **6** (12.5 μM) (**b**), and **7** (25 μM) (**c**) on the expression of viral proteins in HSV-2-infected HeLa cells. Protein levels of HSV-2 ICP8 and gD were assessed at different times post-infection (hpi) by Western blotting. Actin was included as a loading control.

**Extended Data Fig. 3:**
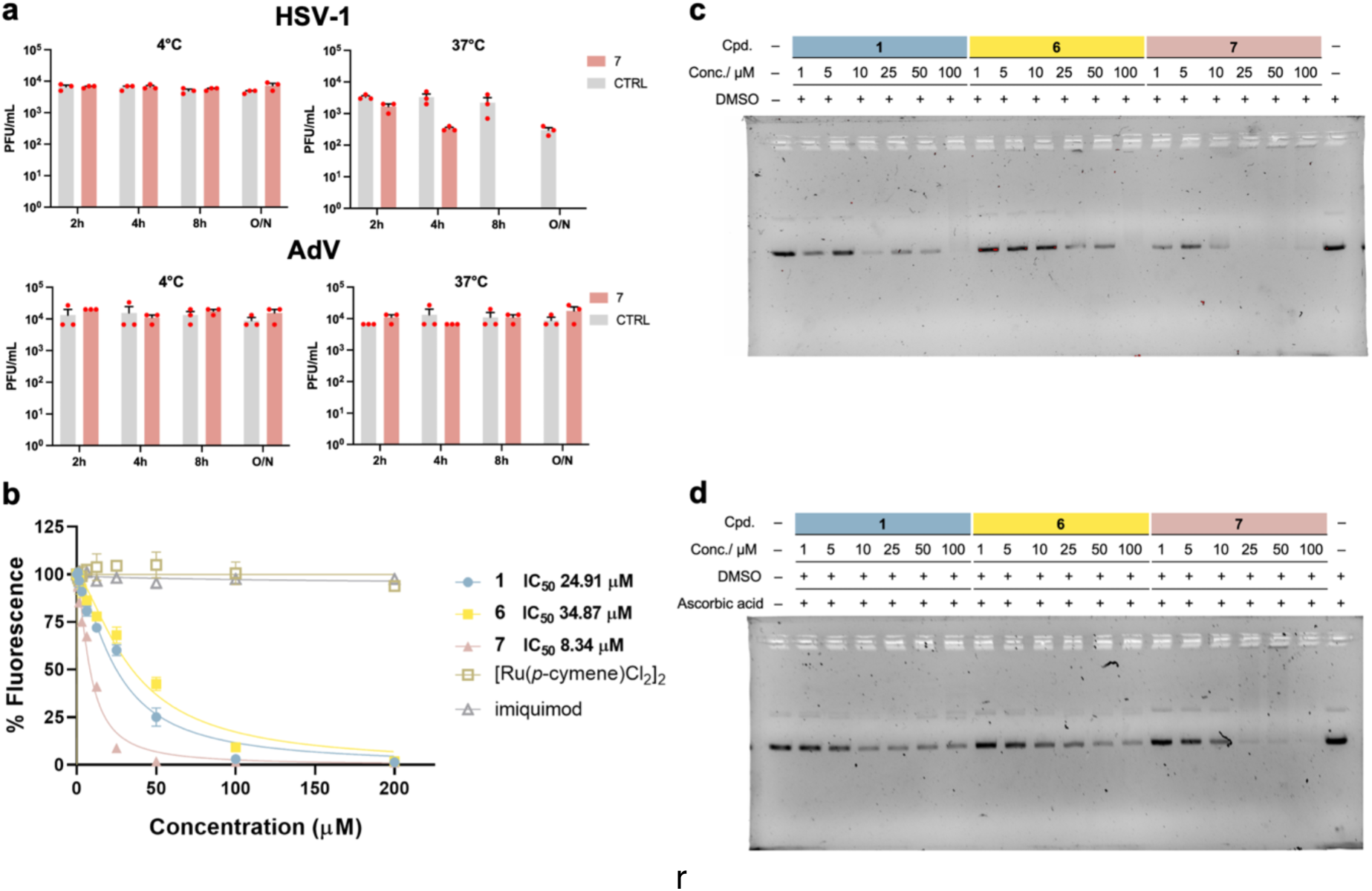
Additional antiviral breadth and virus-directed effects. **a**, Virucidal assays. 10⁶ viral particles of HSV-1 or Adenovirus were incubated with **7** (50 μM) at 4 °C or 37 °C for the indicated times. The mixture was then diluted 1:100 to abolish the antiviral activity of the compound, and residual viral infectivity was quantified by a standard plaque assay. Statistical significance was determined by one-way ANOVA followed by a Bonferroni post hoc test (*ns*, not significant). **b**, Estimation of drug-DNA binding constants. Inhibition of thiazole orange binding to calf thymus DNA was evaluated by the addition of different concentrations of **1**, **6** and **7**. The binding constants of these derivatives to DNA can be estimated and compared by measuring the loss of thiazole orange as a function of added complex. The concentration of Ru-imiquimod complex that produces 50% inhibition of fluorescence, is approximately inversely proportional to the binding constant. Based on this method, the concentrations producing 50% inhibition were determined to be 24.91, 34.87 and 8.34 µM for **1**, **6** and **7**, respectively. Control experiments confirmed that neither the Ru-p-cymene scaffold nor imiquimod alone bind to DNA. **c**, Agarose gel electrophoresis experiments: plasmid pUC19 in the presence of **1**, **6** and **7**. **d**, plasmid pUC19 in the presence of ruthenium complexes **1**, **6** and **7** with addition of excess ascorbic acid (1 mM).

**Extended Data Fig. 4:**
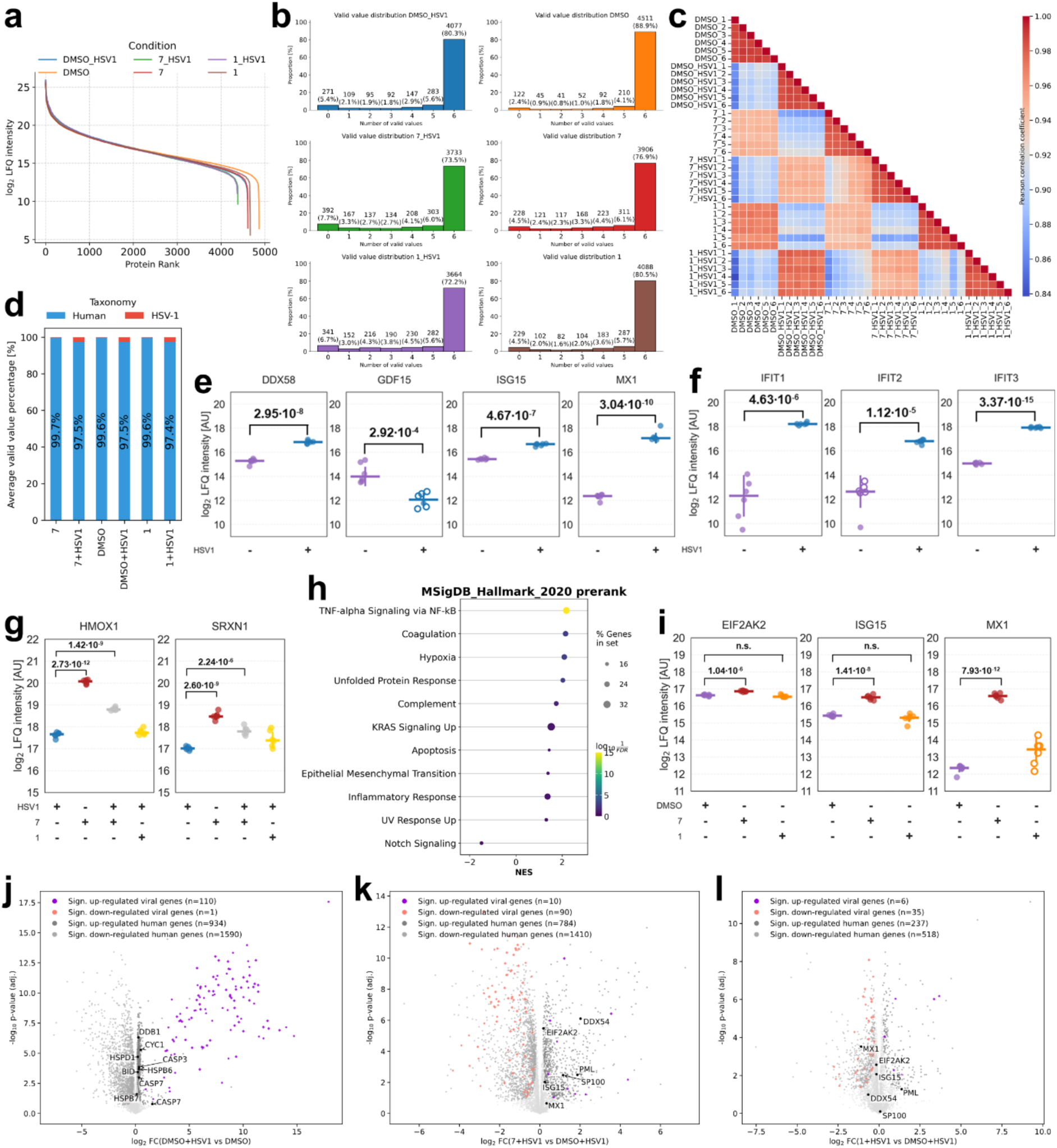
Proteomic quality control and pathway responses. **a**, Protein rank plots show a homogeneous log_2_ LFQ intensity distribution across the different conditions. **b**, Histograms highlighting the valid value distribution of the detected proteins according to biological replicates in each condition. The majority of proteins was identified in six out of six biological replicates. **c**, Correlation heatmap of protein intensity distribution across biological replicates and conditions, confirming a strong impact of HSV-1 infection on the proteome. **d**, Valid value percentage of human and HSV-1 proteins in each experimental condition. Percentages represent the averages of six biological replicates. **e,f**, Point plots of the host type I interferon response upon infection with HSV-1. Adjusted P-values are shown. Hollow circles represent imputed values. **g**, Point plots of NRF2 target genes in the infected condition and **7** treatment alone. Adjusted P-values are shown. **h**, Log_2_ LFQ pre-ranked gene set enrichment analysis performed on the molecular signature database 2020 hallmark gene set of the perturbation of 7 for 24 h in uninfected HFF-1 cells. FDR=0.25. **i**, Point plots of direct **7** antiviral effects in uninfected cells compared to **1** treatment. Adjusted P-values are shown. Hollow circles represent imputed values. **j**, Volcano plot of host cell response to viral infection *via* apoptosis and heat shock proteins 24 h post infection. **k**, Volcano plot of the 24 h perturbation with **7** in uninfected HFF-1 cells. **l**, Volcano plot of the 24 h perturbation with **1** in uninfected HFF-1 cells.

**Extended Data Fig. 5:**
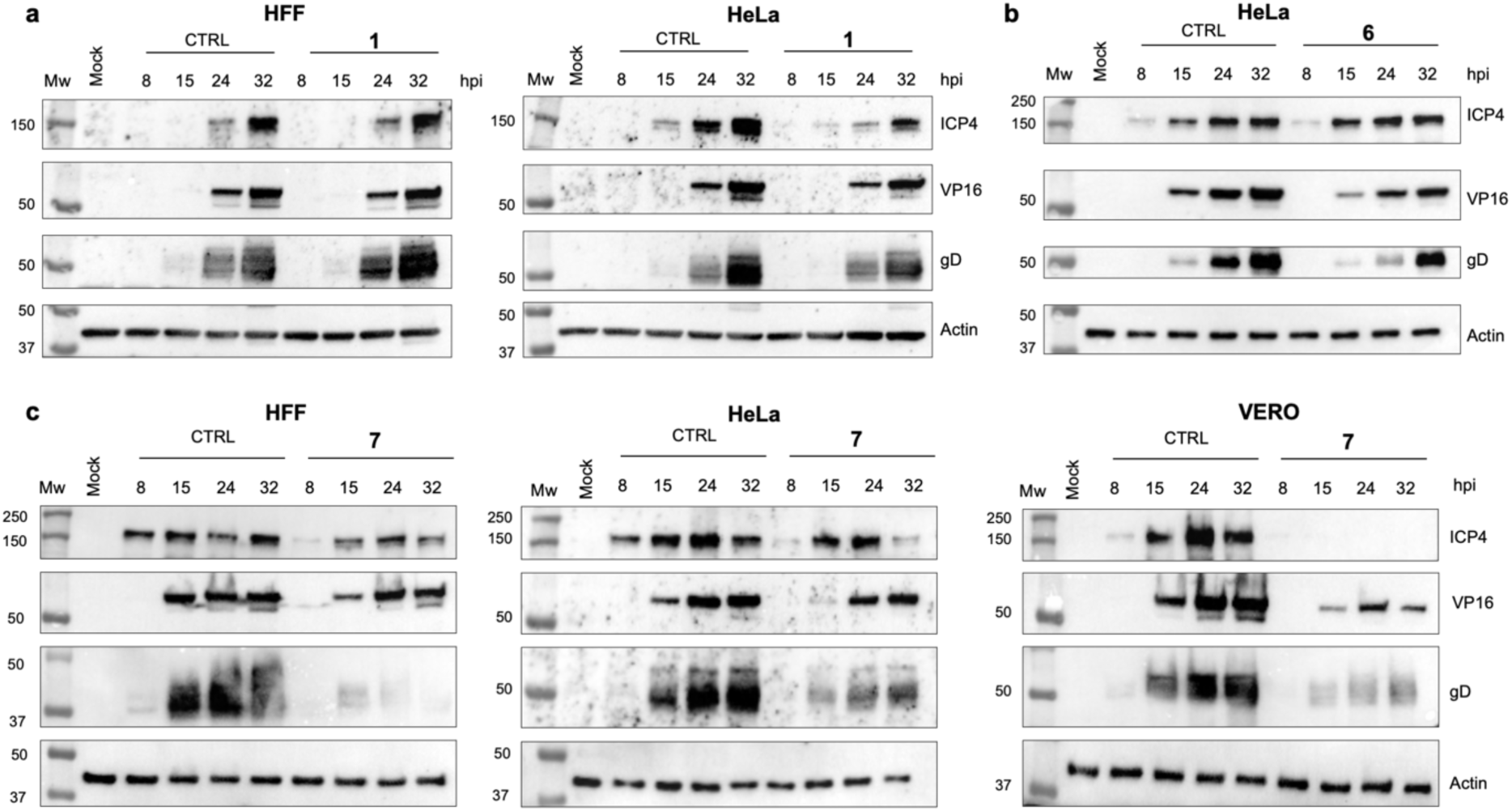
Effects of Ru-imiquimod complexes on HSV-1 protein expression. Western blot analysis of viral proteins ICP4, VP16, and gD at the indicated hours post-infection (hpi) in cells infected with HSV-1 (MOI 1) and treated with the indicated compounds or vehicle control (CTRL). **a**, **1** (50 μM in HFF and 25 μM in HeLa cells); **b**, **6** (12.5 μM in HeLa cells); c, **7** (25 μM in HFF, 12.5 μM in HeLa and 50 μM in VERO cells). Actin was used as a loading control.

**Extended Data Fig. 6:**
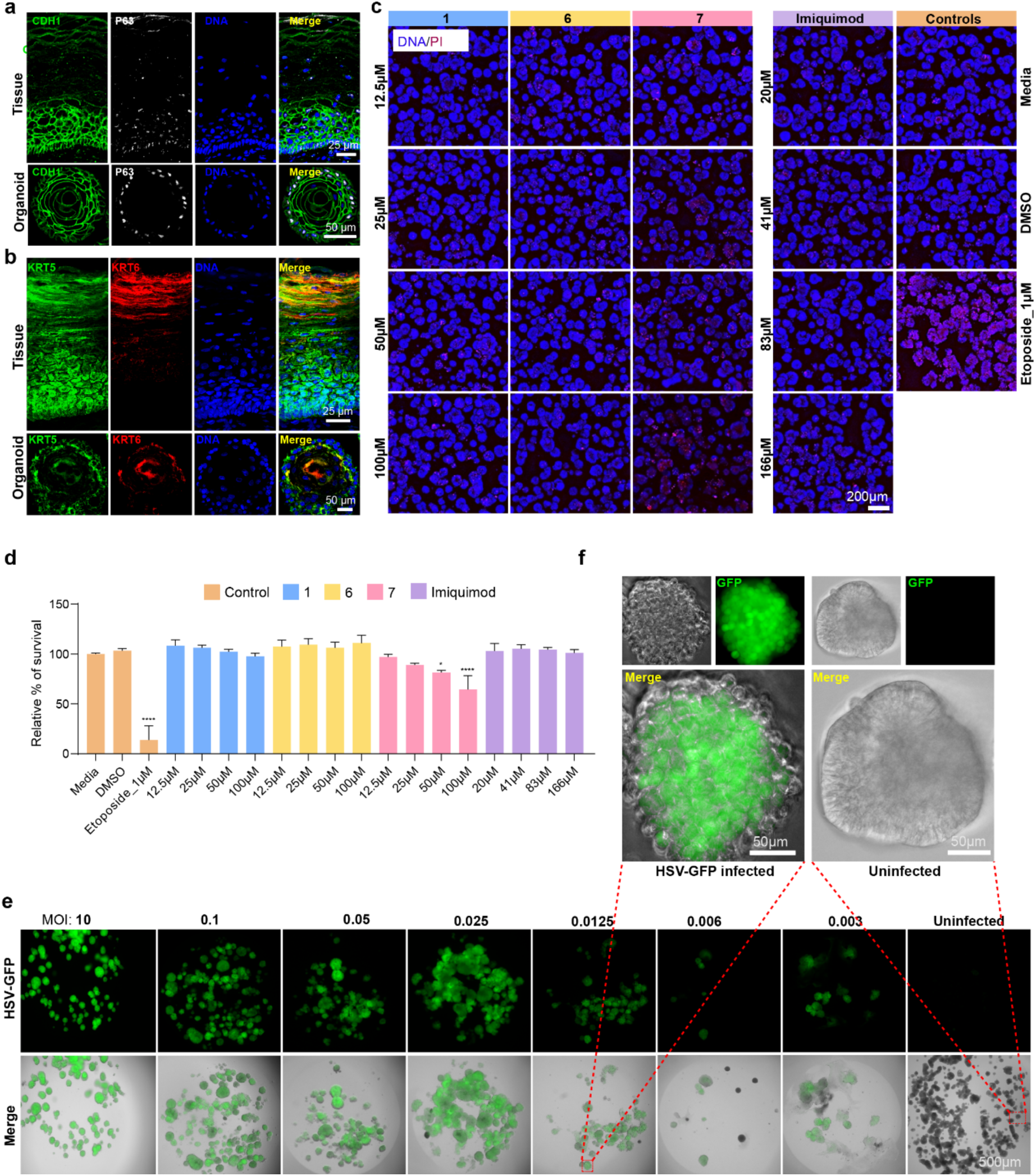
Characterization, viability, and HSV-1 infection of patient-derived ectocervical organoids. **a**,**b**, Immunofluorescence staining of human ectocervical tissue and patient-derived ectocervical organoids showing epithelial architecture and lineage organization. **a**, CDH1 marks epithelial cells, and p63 marks the basal/parabasal compartment. **b**, KRT5 and KRT6 mark basal and suprabasal epithelial compartments, respectively. Nuclei were counterstained with DAPI. **c**,**d**, Assessment of organoid viability following compound treatment. Organoids were stained with Hoechst (blue) to label nuclei and propidium iodide (PI; red) to identify dead cells. **c**, Representative fluorescence images of ectocervical organoids treated with the indicated concentrations of **1**, **6**, **7** or imiquimod, together with medium and DMSO controls and etoposide (1 µM) as a positive cytotoxicity control. **d**, Quantification of relative organoid survival following treatment, normalized to the media control. Bars show mean ± SEM. Statistical comparisons were performed using one-way ANOVA followed by Dunnett’s multiple-comparisons test against the media control. *P* = 0.0101; \*\*\*\**P* < 0.0001. **e**, Representative fluorescence and merged images of ectocervical organoids infected with HSV-1 (17+) Lox-pMCMV-GFP at the indicated multiplicities of infection (MOI). An uninfected organoid control is shown for comparison. **f**, Higher-magnification images of an HSV-1-GFP-infected organoid and an uninfected control from panel **e**, showing GFP-positive epithelial regions in the infected organoid. Scale bars, **a**,**b**, 25 µm (tissue) and 50 µm (organoids); **c**, 200 µm; **e**, 500 µm; **f**, 50 µm. Representative images in **a**–**c** and **e**, **f**, and data in **d**, are derived from three independent biological replicates.

**Extended Data Fig. 7:**
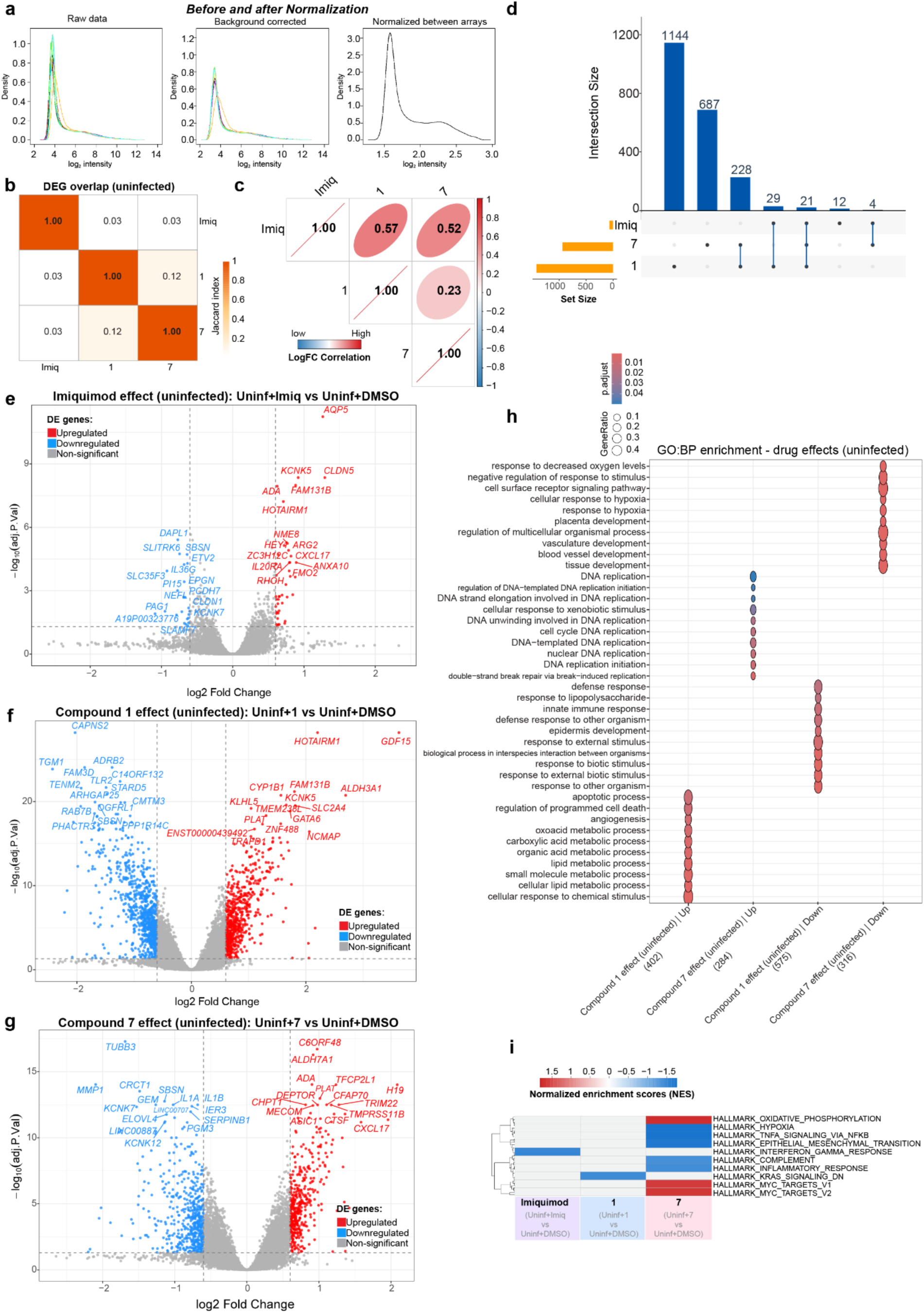
Compound-specific transcriptional responses in uninfected patient-derived ectocervical organoids. **a**, Probe-intensity density distributions for individual arrays before normalization, after background correction, and after between-array normalization; each line represents one array. **b**, Pairwise Jaccard similarity of differentially expressed gene (DEG) sets following treatment with imiquimod, **1** or **7**, relative to DMSO. Color indicate Jaccard index. **c**, Pairwise Pearson correlation of log_2_ fold changes between treatments. Numbers indicate Pearson correlation coefficients; ellipse shape and color indicate correlation magnitude and direction. **d**, UpSet plot showing the intersections between DEG sets for the three compounds in uninfected organoids. Bar heights indicate intersection size; filled circles indicate contributing sets. **e**–**g**, Volcano plots of differential gene expression following treatment with imiquimod (**e**), **1** (**f**), or **7** (**g**) relative to DMSO. Red and blue denote significantly upregulated and downregulated genes, respectively (BH-adjusted *P* < 0.05, log_2_FC > 0.6); selected significant genes are labelled. **h**, Gene Ontology biological process (GO:BP) enrichment among genes significantly upregulated or downregulated by **1** and **7**. Dot size indicates gene ratio and color indicates BH-adjusted *P* value. **i**, Hallmark gene-set enrichment analysis following treatment with imiquimod, **1** or **7** relative to DMSO. Color indicates normalized enrichment score (NES), with positive and negative enrichment shown in red and blue, respectively; white denotes non-significant enrichment (BH-adjusted *P* > 0.05). Data in **a**–**i** are derived from *n* = 5 independent biological replicates per condition.

**Extended Data Fig. 8:**
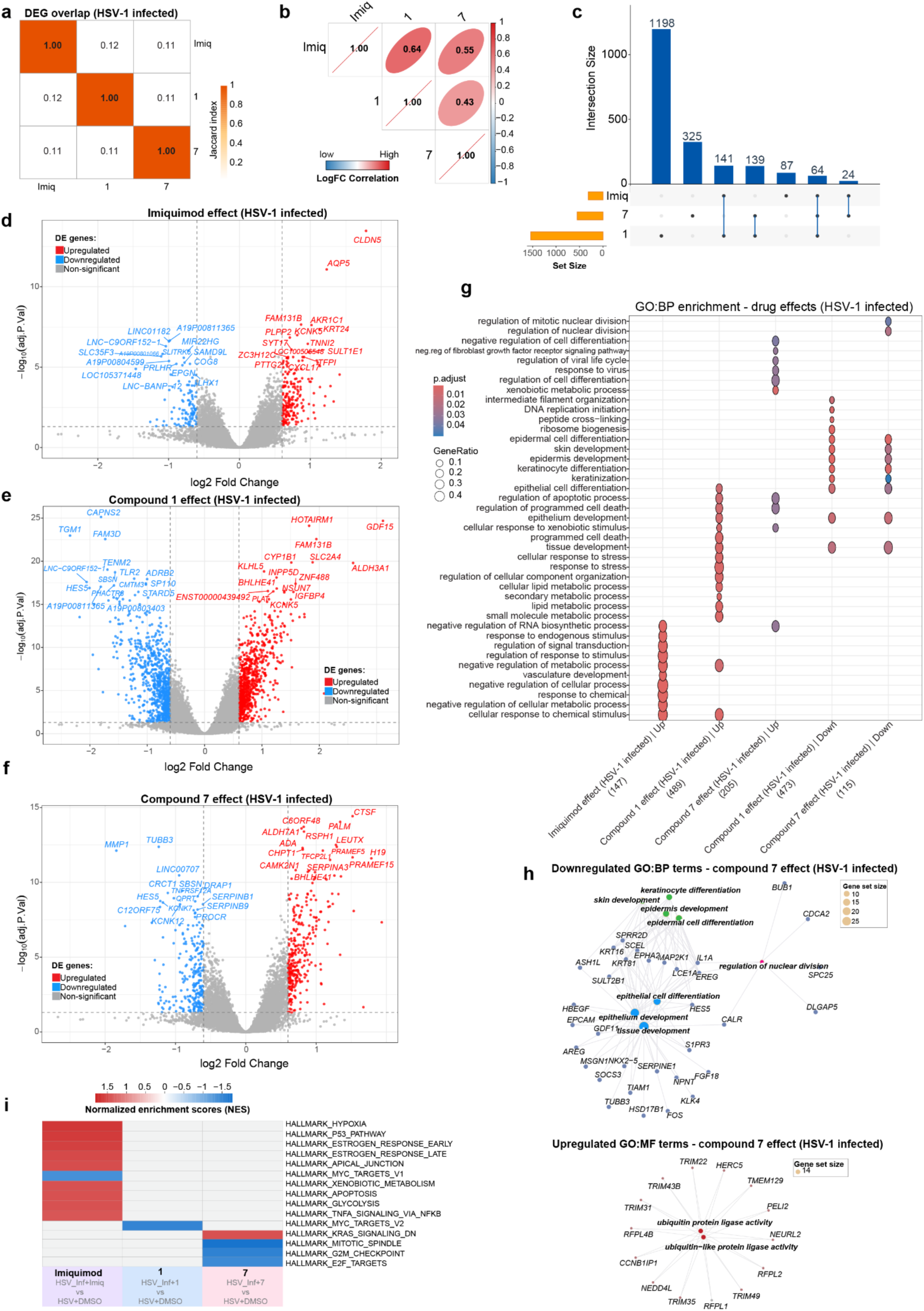
Compound-specific transcriptional responses during HSV-1 infection. **a**, Pairwise Jaccard similarity of DEG sets following treatment with imiquimod, **1** or **7** in HSV-1-infected organoids relative to DMSO. Color indicates Jaccard index. **b**, Pairwise Pearson correlation of log_2_ fold changes between treatment responses in HSV-1-infected organoids. Numbers indicate Pearson correlation coefficients; ellipse shape and color indicate correlation magnitude and direction. **c**, UpSet plot showing overlap among DEG sets induced by imiquimod, **1** and **7**. Horizontal bars indicate total DEG-set sizes, vertical bars indicate exclusive intersection sizes and filled circles denote the sets contributing to each intersection. **d–f**, Volcano plots showing differential gene expression following treatment with imiquimod (**d**), **1** (**e**) or **7** (**f**) relative to DMSO in HSV-1-infected organoids. Red and blue indicate significantly upregulated and downregulated genes, respectively (BH-adjusted *P* < 0.05, log_2_FC > 0.6); selected significant genes are labelled. **g**, Gene Ontology biological-process (GO) enrichment among genes significantly upregulated or downregulated following treatment with imiquimod, **1** or **7** in HSV-1-infected organoids. Dot size indicates gene ratio and color indicates BH-adjusted *P* value. **h**, Gene–term network plots showing enriched GO terms among genes downregulated by **7** (top) and enriched GO molecular-function (GO) terms among genes upregulated by **7** (bottom) in HSV-1-infected organoids. Term-node size indicates gene-set size, and edges connect genes with the enriched terms to which they contribute. **i**, Hallmark gene-set enrichment analysis following treatment with imiquimod, **1** or **7** relative to DMSO in HSV-1-infected organoids. Color indicates normalized enrichment score (NES), with positive and negative enrichment shown in red and blue, respectively; white denotes non-significant enrichment (BH-adjusted *P* > 0.05). Data in **a**–**i** are derived from *n* = 5 independent biological replicates per condition.

**Extended Data Fig. 9:**
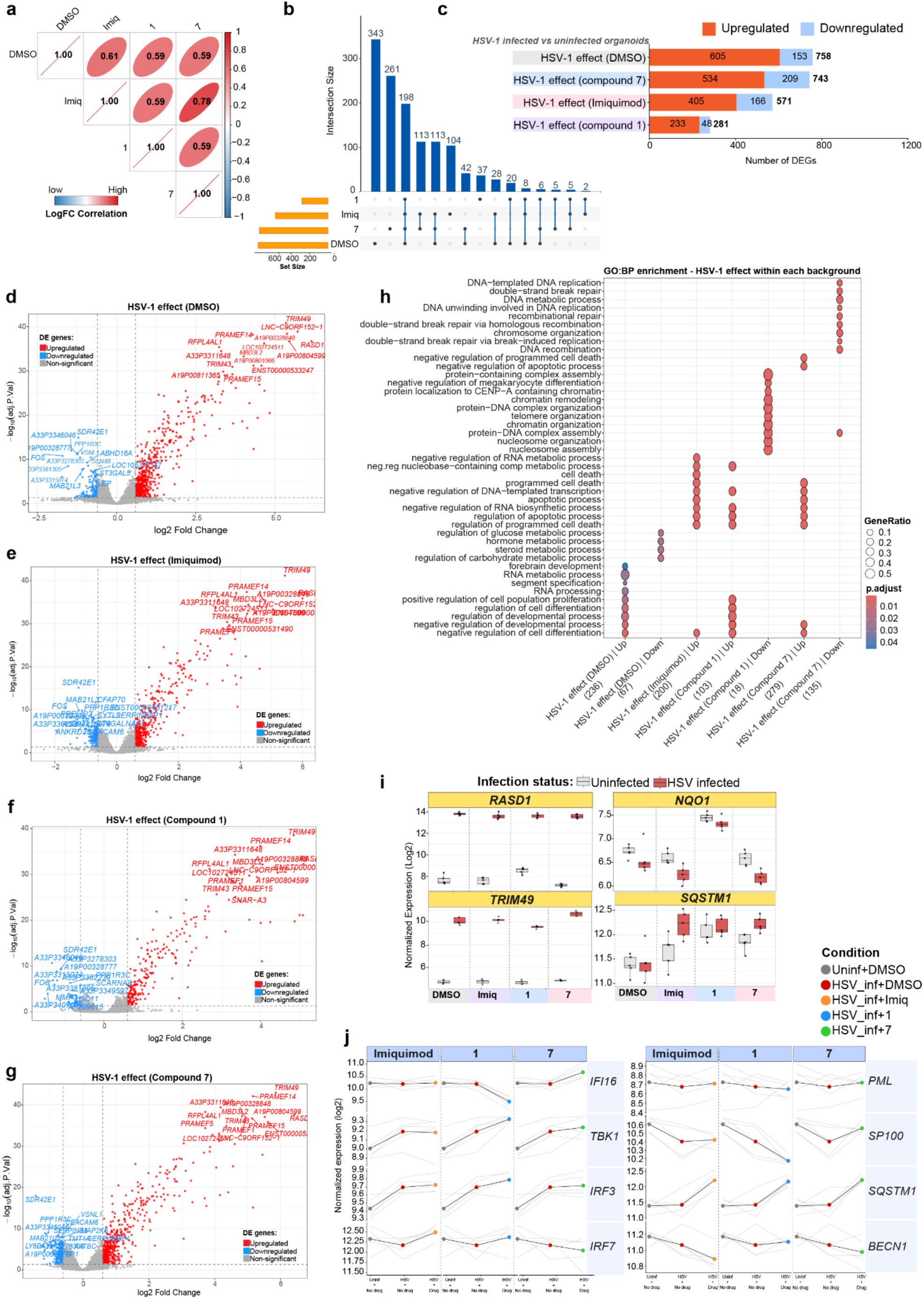
HSV-1 transcriptional responses across treatment backgrounds. **a**, Pairwise Pearson correlation of log_2_ fold changes for the HSV-1 effect across DMSO, imiquimod, **1**, and **7** treatment backgrounds. Numbers indicate Pearson correlation coefficients; ellipse shape and color indicate correlation magnitude and direction. **b**, UpSet plot showing overlap among HSV-1-responsive DEG sets across treatment backgrounds. Horizontal bars indicate total DEG-set sizes, vertical bars indicate exclusive intersection sizes, and filled circles denote contributing sets. **c**, Numbers of significantly upregulated and downregulated DEGs induced by HSV-1 within each treatment background (BH-adjusted *P* < 0.05, log_2_FC > 0.6). **d**–**g**, Volcano plots showing differential gene expression between HSV-1-infected and uninfected organoids within the DMSO (**d**), imiquimod (**e**), **1** (**f**), and **7** (**g**) backgrounds. Red and blue indicate significantly upregulated and downregulated genes, respectively; selected significant genes are labeled. **h**, GO biological process (GO: BP) enrichment among genes upregulated or downregulated by HSV-1 within each treatment background. Dot size indicates gene ratio, and color indicates BH-adjusted *P* value. **i**, Normalized log_2_ expression of selected infection-responsive genes across the indicated treatment backgrounds; color indicates infection status. **j**, Gene expression trajectory plot for DNA-sensing pathway-associated genes across uninfected + DMSO, HSV-1 + DMSO, and HSV-1 + compound conditions. Thin grey lines represent individual biological replicates; the black line represents the group means, and colored points indicate group means. Data in **a**–**j** are derived from 5 independent biological replicates per condition.

**Extended Data Fig. 10:**
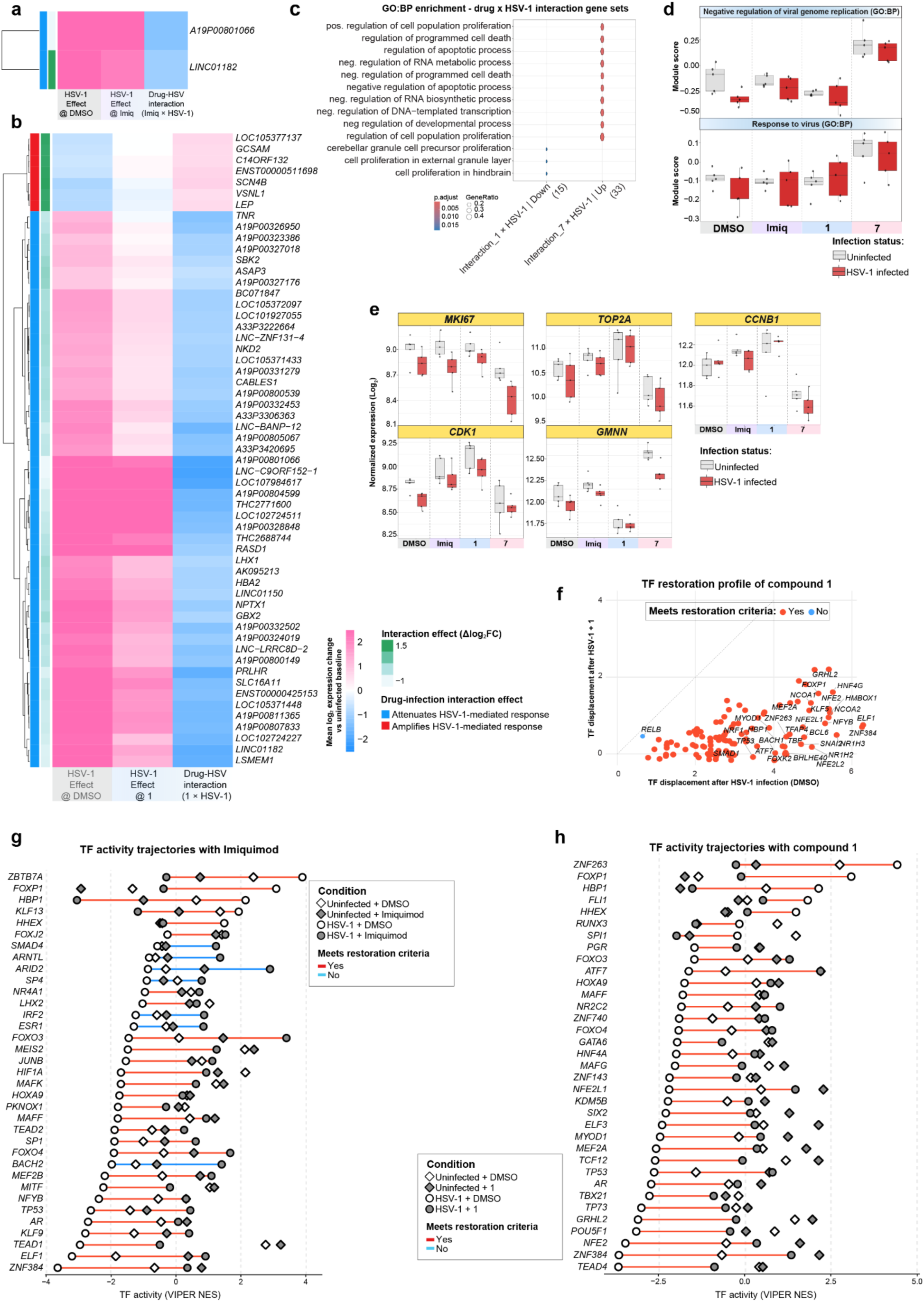
Drug–HSV-1 interaction effects and transcription-factor responses to imiquimod and 1. **a**,**b**, Heatmaps showing genes with drug–HSV-1 interaction effects for imiquimod (**a**) or **1** (**b**). Columns, color scales, and annotations are as in Fig. 5d. **c**, GO biological-process (GO:BP) terms enriched among genes showing positive or negative interaction effects between **1** and HSV-1. Dot size indicates gene ratio, and color indicates BH-adjusted *P* value. **d**, Module scores for the GO:BP gene sets ‘negative regulation of viral genome replication’ and ‘response to virus’ across the indicated conditions, calculated as in Fig. 5f. **e**, Normalized log_2_ expression of the cell-cycle genes *MKI67, TOP2A, CCNB1, CDK1* and *GMNN* across the indicated conditions. In **d**,**e**, box color indicates infection status and points represent individual biological replicates. **f**, Transcription-factor (TF) restoration profile following treatment with **1**; axes, colors, and restoration criteria are as in Fig. 6e,**f**.**g**,**h**. TF activity profiles following treatment with imiquimod (**g**) or **1** (**h**); symbols, colors, and TF-selection criteria are as in Fig. 6g. Data in **a**–**h** are derived from five independent biological replicates per condition.

