## Supplementary Information for "Ruthenium Coordination Architecture Redirects Imiquimod Antiviral Activity"

### 1. Experimental section

#### 1.1. Materials

All reactants and solvents were purchased at the highest purity available and used without further purification. Dichloro(p-cymene)ruthenium(II) dimer (p-cym) and 5-fluorouracil (5-fu) were obtained from Acros Organics. Pyridine (py) from Scharlau; imidazol (im), caffeine (caf), ammonium hexafluorophosphate ($NH_4PF_6$) and silver nitrate ($AgNO_3$) from Sigma-Aldrich; imiquimod (imq) and indazol (in) from Indagoo; 1, 3, 5-triaza-7-phosphadmantane (pta) from Alfa Aesar. Methanol (MeOH) and dimethylsulfoxide (DMSO) were obtained from Scharlau and dichloromethane (DCM) from Fisher Scientific. DMEM medium for stability experiments was obtained from Fisher Scientific. Ultrapure water was deionized and filtered with a Millipore MilliQ system.

#### 1.2. Instrumentation and methods

**Nuclear magnetic resonance (NMR)**
$^{1}H$ and $^{13}C$ NMR spectra were recorded in 5 mm NMR tubes using standard pulse programs and a Fourier TM Bruker 300 or 400 NMR spectrometer at room temperature. The solvents used were deuterium oxide ($D_2O$), chloroform-$d_1$ ($CDCl_3$), methanol-$d_4$ ($CD_3OD$), dimethylsulfoxide-$d_6$ ($(CD_3)_2SO$) and acetone-$d_6$ ($(CD_3)_2CO$). These were obtained from Euroisotop. The chemical shifts (δ, quoted in parts per million, ppm) in $^{1}H$ and $^{13}C$ NMR were internally referenced to the residual peak of the corresponding solvents: HDO ($^{1}H$: 4.79 ppm), $CD_2HOD$ ($^{1}H$: 3.31 ppm and $^{13}C$: 49.00 ppm), $(CD_5H)_2SO$ ($^{1}H$: 2.50 ppm and $^{13}C$: 39.52 ppm). Coupling constants, *J*, are reported to the nearest 0.1 Hz. Data were processed using MestreNova, version 11.0 (Mestrelab Research, S.L).

**Mass spectrometry (MS)**
The analysis consisted of a chromatographic separation in an ultrahigh-performance liquid chromatography (UPLC, Acquity system from Waters Cromatografia S.A.) coupled to a high-resolution mass spectrometer (Synapt G2 from Waters Cromatografia S.A., time-of-flight analyser (TOF) by an electrospray ionization source in positive mode (ESI+). The chromatographic separation was achieved using an Acquity UPLC BEH C18 column (1.7 µm, 2.1×50 mm i.d.) with an Acquity UPLC BEH C18 1.7 µm VanGuard pre-column (2.1×5 mm) (Waters Cromatografia S.A.) and a binary A/B gradient (solvent A: water with 0.1 % formic acid and solvent B: methanol). The gradient program was established as follows: initial conditions were 5 % B, raised to 100 % B over 2.5 min, held at 100 % B until 4 min, decreased to 5 % B over the next 0.1 min, and held at 5 % B until 5 min for re-equilibration of the system prior to the next injection. A flow rate of 0.25 mL/min was used, the column temperature was 30 °C, the autosampler temperature 4 °C and the injection volume 5 µL.
High-resolution mass data were acquired in SCAN mode, using a mass range of 50–1300 u in resolution mode (FWHM ≈ 20,000) and a scan time of 0.1 s. The source temperature was set to 120 °C and the desolvation temperature to 350 °C. The capillary voltage was 0.7 kV and the cone voltage 15 V. Nitrogen was used as the desolvation and cone gas at flow rates of 600 L/h and 10 L/h, respectively. Before analysis, the mass spectrometer was calibrated with a sodium formate solution. A leucine-enkephalin solution was used for the lock mass correction, monitoring the ions at mass-to-charge ratios (m/z) 556.2771 and 278.1141. All of the acquired spectra were automatically corrected during acquisition based on the lock mass.

Samples were dissolved in methanol or dimethyl sulfoxide at a concentration of 1 mg/ml and diluted to 20 µg/mL for the analysis.

**UV-vis spectroscopy.**
UV-vis absorption data were collected using a JASCO V-730 spectrophotometer (JASCO Corp., Tokyo, Japan) with 1 cm path length quartz cuvettes (800 µL). Spectra were recorded at different times in aqueous solutions or buffers from 600 to 200 nm unless otherwise stated. Data processing was carried out using Spectra Manager™.

**Elemental analysis.**
Elemental analysis was performed on a TruSpec Micro elemental (CHNS) analyzer (LECO Corp.). Samples were weighed on a microbalance, using 1-2 mg per sample. For calibration, sulfamethazine ($C_{12}H_{14}N_4O_2S$), a NIST-certified reference material, was used. The presented values are the average of determinations in duplicate.

**Stability of compounds in solution by NMR spectroscopy.**
Methanol and dimethyl sulfoxide solutions of the Ru(II) complexes were prepared at ca. 4-5 mM by dissolving them in methanol-*d*$_4$ ($CD_3OD$) or dimethylsulfoxide-*d*$_6$ ($(CD_3)_2SO$). The $^1$H NMR was recorded in situ after sample preparation (data collection at ca. 5 min after dissolution) and at different times up to 48h (24h intervals). Speciation of each complex was evaluated following the changes in the $^1$H NMR signals.

**Stability of compounds in DMEM cell culture media by UV-vis spectroscopy.**
Dimethyl sulfoxide stock solutions of the Ru(II) complexes **1**, **6,** and **7** at 0.5-2 mM were prepared by dissolving them in DMSO (HPLC grade). Then, samples were diluted to 100 µM for complexes **1** and **6**, and 25 µM for complex **7** with cell culture media. The UV-vis absorption spectra were recorded *in situ* after sample preparation at 298 k and at 1 h intervals during 6 h, then 24 h and 48 h. Speciation of each complex was evaluated following the changes in the UV-visible spectra.

**Single crystals X-ray diffraction.**
Single crystals for **1**, **2**, **4**, **6,** and **7** were selected and mounted on a SuperNova, Dual, Cu at home/near, HyPix diffractometer. The crystal was kept at 170.00(10) K during data collection. Using Olex2,[1] the structure was solved with the SHELXT[2] structure solution program using Intrinsic Phasing and refined with the SHELXL[3] refinement package using Least Squares minimisation.
CCDC 2583749–2583753 contains the supplementary crystallographic data for compounds **1**, **2**, **4**, **6,** and **7**. These data can be obtained free of charge from The Cambridge Crystallographic Data Centre via www.ccdc.cam.ac.uk/data_request/cif

**Competitive thiazole orange displacement assay – interaction with calf thymus DNA**
*Competitive thiazole orange displacement assay*. A working solution containing 20.0 µM UltraPure calf thymus DNA (ct-DNA; Invitrogen, Cat. No. 15633019, $\varepsilon_{260}$ = 12,824 $M^{-1}(bp)^{-1}\cdot cm^{-1}$) and 25.2 µM thiazole orange (TO; TargetMol, T19065) was prepared in HEPES buffer (80 mM HEPES, 40 mM NaCl, pH 7.4). Stock solutions of metal complexes, metal salts, and free ligands were prepared at a concentration of 4.0 mM in DMSO and subsequently diluted with ultrapure water. For the assay, 50 µL of the DNA–TO working solution were placed into each well of a 96-well microplate, excluding the blank wells, where 100 µL of HEPES buffer alone were placed. Serial dilutions of the test compounds were added to the wells containing the DNA–TO solution, and the final volume in each well was adjusted to 100 µL. This resulted in final concentrations of 4.0 µM for ct-DNA and 12.6 µM for TO. The plate was incubated at room temperature for 1 hour prior to fluorescence measurement. Fluorescence

readings were obtained using a BioTek Synergy HT multi-mode microplate reader, with excitation and emission wavelengths set at 490 nm and 530 nm, respectively.
Each concentration of the test compounds was analyzed in quadruplicate. Apparent binding constants ($K_{app}$) were calculated using the equation:

$$\frac{K_{app} = K_e \times 12.6}{C_{50}}$$

where $K_e = 8.8 \times 10^6 M^{-1}(bp)^{-1}$ and $C_{50}$ is the concentration of the test compound required to displace 50% of bound TO.

**Drug interaction with DNA by agarose gel electrophoresis**
For the interaction of the compounds with plasmid DNA (pUC19, ThermoScientific, Cat. No. SD0061), samples of the different complexes at different concentrations (1, 5, 10, 25, 50 and 100 µM) were incubated with pUC19 plasmid (200 ng per sample, estimated spectrophotometrically) in HEPES buffer (80 mM HEPES, 40 mM NaCl, pH 7.4) for a final sample volume of 20 µL. The samples were incubated for 24 h at 37 °C and after that, 4 µL of loading buffer (10 mM Tris-HCl at pH 7.6, 60 mM EDTA, 60% glycerol, 0.03% xylene cyanol and 0.03% bromophenol blue) were added. A volume of 20 µL of each of these mixtures was loaded onto an agarose gel (1.2%). The gel was run for 90 min at 70 V in 1x TAE buffer (40mM Tris, 20 mM acetate and 1mM EDTA, pH 8.6). Next, bands were visualized under UV light and imaged digitally.

**Cell and virus culture**
Human foreskin fibroblasts (HFF; American Type Culture Collection, ATCC SCRC-1041), the cervix adenocarcinoma cell line HeLa (ATCC CCL-2), and the African green monkey kidney cells (VERO, ATCC CCL-81), were cultured in Dulbecco's modified Eagle's medium (Neo DMEM, NeBiotech Clinsciences Group), supplemented with 10% heat-inactivated fetal bovine serum (FBS), 2 mM glutamine, 1 mM sodium pyruvate, 100 U/mL penicillin, and 100 µg/mL streptomycin sulfate (Aurogene). The clinical isolates of human herpes simplex virus type 1 (HSV-1), human herpes simplex virus type 2 (HSV-2), human adenovirus (AdV) (kindly provided by Valeria Ghisetti, "Amedeo di Savoia" Hospital, Turin, Italy), the HSV-1($17^+$)$Lox_{pgMCMV}$mGFP (a generous gift from Beate Sodeik, Hannover Medical School, Hannover, Germany)[4] and the HSV-1 strain Kos (ATCC VR-1493) were used for the study. The viruses were propagated and titrated by plaque assay on VERO cells, as previously described.[5] The acyclovir (ACV) and foscarnet (PFA)-resistant HSV strains were provided by Graciela Andrei (Department of Microbiology, Immunology and Transplantation, Catholic University Leuven, Leuven, Belgium) and produced as previously described.[6–9]

**Cytotoxicity assay**
The cytotoxicity of the investigated compounds was evaluated by using a system based on the 3-(4,5-dimethylthiazol-2-yl)-2,5-diphenyltetrazolium bromide compound (MTT, Sigma-Aldrich). HFF, HeLa, and VERO cells were seeded in a 96-well culture plate at a density of about $4 \times 10^4$ cells/well in 100 µL of medium. Following overnight incubation at 37°C in a 5% $CO_2$ humidified atmosphere, the medium from the wells was removed, and cells were exposed to increasing concentrations of the compounds or vehicle (DMSO), in a final volume of 100 µl/well. After 48 h of incubation, the number of viable cells was determined using an MTT assay. MTT solution (500 µg/mL in serum-free medium) (Sigma-Aldrich) was added to each well, and the plates were incubated for 2 h. At the end of the incubation period, the medium was removed, and the formazan product was dissolved in 50 µL of DMSO. Cell viability was evaluated by measuring the absorbance at 570 nm, using a VICTOR3 1420 multilabel counter (Perkin–Elmer). All experiments were done in triplicate. The readings were converted to the percentage of control (% cell survival). $CC_{50}$ was defined as the concentration of each

compound that reduced cell viability to 50% compared to cells treated with vehicle (DMSO) alone, producing 50% cell death (AAT Bioquest, Inc., Sunnyvale, CA, USA, https://www.aatbio.com/tools/ic50-calculator).

**Virus yield reduction assay**

HFF, HeLa, and VERO cells were seeded in 24-well plates. Following 24 h of incubation, cells were pre-treated with different concentrations of the compounds or the vehicle control (DMSO) for 2 h at 37 °C. They were then infected with HSV-1 or HSV-2 at MOI of 0.1 PFU/cell in the presence of the compounds. Following virus adsorption (2 h at 37 °C), the viral inoculum was removed, and the cultures were maintained in a medium that contained the corresponding molecule for 48 h. The cells and supernatants were then harvested and disrupted using three freeze (liquid nitrogen)/thaw (37 °C) cycles. The extent of viral replication was subsequently assessed by titrating the infectivity of the supernatants of the cell suspensions in VERO cells by standard plaque assay, as previously described.[5] Plaques were microscopically counted, and the mean plaque counts for each drug concentration were expressed as PFU/mL. All experiments were done in triplicate. $IC_{50}$ was defined as the concentration of each compound that reduced plaque formation to 50% compared to untreated cells (AAT Bioquest, Inc., Sunnyvale, CA, USA, https://www.aatbio.com/tools/ic50-calculator).

**Time of addition assay.**

Cells were seeded in 24-well plates (~200,000 cells/well) and treated with the indicated compounds or vehicle solution according to the following schedules: i) throughout the experiment; ii) 2 h pre-infection as pre-treatment; iii) only during the 2 h of viral adsorption; iv) post-infection until sample collection. In all conditions, cells were infected with HSV-1 (MOI 0.1 PFU/cell). After 48 h, cells and supernatants were collected and disrupted using three freeze (liquid nitrogen)/thaw (37 °C) cycles. The level of virus replication was then assessed by titrating the infectivity of the samples by a plaque assay as described in the virus yield reduction assay section.

**Western blot analysis.**

Cells were seeded in 6-well plates (~$10^6$ cells/well), treated with the compounds as described in the virus yield reduction assay section, and infected with HSV-1 or HSV-2 at MOI 1. Cells were collected at different time points, and whole-cell protein extracts were prepared using Radioimmunoprecipitation Assay (RIPA) buffer. Thirty µg of protein extracts were subjected to electrophoresis on sodium dodecyl sulfate-polyacrylamide (SDS) gels and transferred to Immobilon-P membranes (Bio-Rad). Membranes were blocked with 5% non-fat dry milk in TBS-Tween 0.05% and incubated overnight at 4 °C with the primary antibodies. The following mouse monoclonal primary antibodies were used: anti-HSV-1 ICP4 (clone 10F1, H1A021), anti-HSV gD (clone 2C10, HA025), anti-ICP8 (clone H793, VS-P1115), anti-HSV-2 ICP8 (clone 4E6, VS-H2A024) (all from Virusys Corporation/SeraCare), anti-HSV-1 VP16 (clone 1-21, sc-7545, Santa Cruz Biotechnology), Anti-α-Actin-1 (ACTA1) rabbit polyclonal antibody (A2066, Sigma-Aldrich) (all at 1:1000 dilution in 5% non-fat dry milk, TBS-Tween 0.05%). After washing with TBST buffer (500 mM NaCl, 20 mM Tris pH 7.4, 0.05% Tween 20), the membranes were incubated with an HRP-conjugated anti-mouse or anti-rabbit secondary antibody (1:2000 diluted, 1706516 and 1706515, respectively, Bio-Rad Laboratories) for 2 h at room temperature, and immunocomplexes were visualized using an enhanced chemiluminescence detection kit (SuperSignal West Pico Chemiluminescent Substrate, ThermoFisher Scientific).

**Virucidal assay**

Virucidal activity was assessed by incubating approximately $10^6$ PFU of HSV-1 or AdV with **7** (50 µM) or with vehicle control, in DMEM, in a final volume of 100 µL. Virus–compound mixtures were incubated for the indicated times at either 4 °C or 37 °C. Following incubation, samples were serially diluted to reduce the compound concentration below its inhibitory range, and residual viral infectivity was quantified by plaque assay, as described above.

**Statistical Analysis**

The statistical tests were performed using GraphPad Prism version 8.0.2 for Windows (GraphPad, Boston, MA, USA. Data were presented as the mean value and standard error of the mean (SEM). Differences were considered statistically significant for $p < 0.05$ ($p < 0.05$ *; $p < 0.01$ **; $p < 0.001$ ***).

**Proteomic analysis**

**Whole cell lysate collection**

Control, treated (**1** and **7**) and the respective HSV-infected (MOI 1) HFF-1 cells were washed twice with 1 mL PBS. Next, 80 µL of 4% w/v sodium deoxycholate in Tris-HCl 2M (pH 8.5; SDC buffer) was added to each well and cells were scraped and transferred to Eppendorf tubes. Fifty microliters of SDC buffer were used to wash the wells, and this content was also transferred to the tubes, totaling 130 µL per replicate. The tubes were heated on a heating block and shaken for 5 min at 95 °C. Samples were stored at –20 °C until further processing.

**Protein quantification**

Whole-cell lysates stored at -20 °C were thawed at room temperature, briefly centrifuged at high rpm on a tabletop micro-centrifuge and subsequently heated to 95 °C in a pre-heated Thermomixer (Eppendorf) at 1400 rpm for five minutes. Samples were then sonicated three consecutive times for one second at 80% power with an ultrasonic homogenizer (UW 2070, BANDELIN electronic GmbH & Co. KG). The protein amount was quantified by means of the Bicinchoninic acid (BCA) assay. Albumin fraction V (Carl Roth GmbH + Co. KG) was used to prepare a ten-point calibration curve in a 96-wellplate. A photometric plate reader (Multiskan Go, ThermoFisher Scientific) was used to measure the extinction of the wells at 562 nm. The protein amount was quantified against a linear regression of a bovine serum albumin calibration curve.

**Protein digestion**

The protein amount was normalized to 20 µg for digestion according to a StageTip workflow.[10] Samples were adjusted to a final volume of 90 µL with SDC before the addition of 2 µL of 500 mM Bond-Breaker™ Tris(2-carboxyethyl)phosphine hydrochloride pH 7 (Sigma-Aldrich). Samples were briefly vortexed and incubated in a Thermomixer pre-heated to 45 °C at 1200 rpm over the course of five minutes. All samples were left to cool to room temperature, briefly centrifuged at high rpm before the addition of 5 µL of 800 mM 2-Chloroacetamide (Alfa-Aesar). Again, all samples were briefly vortexed and incubated for five minutes at 45 °C and 1200 rpm on a Thermomixer. Subsequently, samples were left to cool to room temperature, briefly centrifuged and placed on ice. Protein digestion was performed at an enzyme (Trypsin-LysC, Promega) to protein ratio of 1:20 over the course of 16 h on a Thermomixer at 37 °C and 1200 rpm.

Peptide desalting was performed using StageTips.[2] StageTips were prepared by punching out two disks of a polystyrenedivinylbenzene-reversed phase sulfonate material (Empore 2241 SDB-RPS, 12 µm particle size, 47mm; CDS Analytical LLC) which were stacked in 200 µL pipet tips shortly before use. The solid phase material was conditioned with 150 µL of loading solvent (99%vol/vol isopropanol, 1%vol/vol trifluoroacetic acid). Following the digestion, samples were nearly dried in a Vacuum concentrator (miVac DUO, Genevac) at 40 °C. The residual sample volume was loaded on StageTips by repeatedly pipetting up and down after

the addition of 120 µL loading solvent to each sample. Sample tubes were rinsed with 40 µL loading solvent. The solution was then quantitatively transferred to the StageTips. This was followed by a seven-minute centrifugation step at 700 g. The tips were centrifuged at 1500 g for five minutes after the addition of further 100 µL of loading solvent. Subsequently, 100 µL of wash solvent (94.8%vol/vol $H_2O$, 5%vol/vol acetonitrile, 0.2%vol/vol trifluoroacetic acid) were added to the tips. Tips were then centrifuged at 1500 g for eight minutes. Peptides were eluted from the solid phase material into LC-MS glass inlets with the addition of 30 µL elution solvent (59.7%vol/vol acetonitrile, 39.8%vol/vol $H_2O$, 0.5%vol/vol $NH_4OH$) followed by a centrifugation at 500 g for four minutes. Further 30 µL of elution solvent were added to the tips prior to the final seven-minute centrifugation step at 1500 g. Peptides were dried in the vacuum concentrator and stored at –20 °C until measurement.
Dried peptides were reconstituted in 5 µL of a 10 $fmol{\cdot}\mu L^{-1}$ synthetic peptide mix in 30% formic acid followed by 40 µL of LC loading solvent 98%vol/vol $H_2O$, 2%vol/vol acetonitrile, 0.05%vol/vol trifluoroacetic acid).

**Nano-flow liquid chromatography tandem mass spectrometry (nLC-MS/MS)**
A Dionex UltiMate™ 3000 RSLCnano system (Thermo Fisher Scientific) was employed for the chromatographic separation of the peptides. Solvent A (99.9% $H_2O$, 0.1% formic acid) was used to load 5 µL of reconstituted samples onto an Acclaim™ PepMap™ C18 100 (Thermo Fisher Scientific) pre-column at a 10 $\mu L{\cdot}min^{-1}$ flow rate. A gradient was run from 12% to 42% solvent B (79.9% acetonitrile, 20% $H_2O$, 0.1% FA) over 90 minutes at a flow rate of 300 $nL{\cdot}min^{-1}$ to achieve separation on the analytical column (Aurora Ultimate CSI UHPLC CoreJet™ emitter column, 25 cm × 75 µm, 1.7 µm, C18, Ionopticks). The inclusion of equilibration and washing phases resulted in a total runtime of 140 minutes.
The nanoLC system was hyphenated with a timsTOF Pro (Bruker Daltonics) mass spectrometer acquiring spectra in data-dependent mode using parallel accumulation serial fragmentation for label-free quantification of the proteins. Scan ranges for the acquisition of MS1 and MS2 spectra were both set from 100 to 1700 m/z. Ion mobility separation was achieved with an inverse reduced ion mobility (1/k0) scan ranging from 0.6 to 1.6 $Vs{\cdot}cm^2$, while the total cycle time for 10 PASEF MS/MS scans was 1.16 seconds. In total 321096 MS1 and 1794501 MS2 scans were acquired. Capillary voltage 1650 V, 3 L/min dry gas, dry temp 180 °C.

**Data processing and evaluation**
The raw data files resulting from the label free quantification proteomics measurements were processed by MaxQuant (version 1.6.17.0), which allows the identification and relative quantification of the proteins present in the samples. The raw data was searched against the SwissProt "homo sapiens" (14.12.2019 with 20380 canonical entries), as well as the "herpes simplex virus 1" database (03.06.2025 with 8969 both reviewed and unreviewed entries). The database search was performed with peptide-spectrum match and protein false discovery rates of 0.01. "Match between runs" was enabled with an alignment time window of 20 minutes and a matching time window of 0.7 minutes. A maximum of two missed cleavages were allowed. The acetylation of the protein N-terminus and Methionine oxidation were defined as variable modifications, while Cysteine carbamidomethylation was defined as a fixed modification for the database search. The minimal threshold for protein identifications was defined as the identification of two peptides, one of which had to be unique to the protein. The database search yielded a matrix consisting of 5681 protein groups. The number decreased to 5466 after removing common contaminants, entries that were matched to reversed sequences as well as proteins that were only identified by site. The resulting dataset was additionally filtered to contain only proteins that were present in at least five out of six replicates in at least one of the experimental conditions. This step removed further 388 proteins. LFQ values were $log_2$-transformed. The described filtering steps and the overall data processing

were performed using Pandas 2.2.2 in Python (version 3.12.7).[11] Perseus 1.6.14.0[12] was employed for missing value imputation from a normal distribution (downshift: 1.8, width: 0.3). Additionally, Perseus was also employed in the calculation of statistical significance of protein regulation by using two-sided *t* tests with FDR = 0.05, *s*0 = 0.1 and 250 permutations. Data visualization was performed using Matplotlib[13] or Seaborn (version 0.13.2).[14] Gene set enrichment analyses were performed using GSEApy (version 1.1.9).[15]

**Antibodies**

The following antibodies were used for immunofluorescence: Rabbit anti–KRT5–Alexa Fluor 488 (1:300, Abcam, ab193894), mouse-anti-KRT6 (1:50, Abcam, ab18586), donkey anti-mouse Cy3 (1:150, Dianova, 715-166-020), P63, CDH1and for labeling the DNA, 4′,6-diamidino-2-phenylindole (DAPI, Roche, 10236276001) was used.

**Patient material**

Human cervical samples were provided by the Department of Gynecology, Charité University Hospital, and Vivantes Auguste-Viktoria Klinikum, Berlin. Usage for scientific research was approved by the Ethics Commision of the Charité University Hospital, Berlin, Germany (EA1/059/15); informed consent to use their tissue for scientific usage was obtained from all donors. The study complies with all relevant ethical regulations regarding research involving human participants. Biopsies were sourced from standard surgical procedures for benign gynecological disease. Only anatomically normal tissue biopsies from anonymous female donors aged between 35 and 75 were processed within 2–3 h after removal.

**Human ectocervical primary organoid culture**

Normal human ectocervical organoids (derived from lab biobank) were cultured from single epithelial cells using Matrigel as described in our previously published protocol.[16] Briefly, single cell epithelial cells were maintained in 2D by co-culturing with mitotically inactive 3T3-J2 cell line in a tissue culture flasks until they reached 70–80% confluence in human ectocervical primary cell medium (ADF++ media (Advanced DMEM/F-12 (Invitrogen, 12634) supplemented with 12mM HEPES, 1% GlutaMax), supplemented with 1% B27, 1% N2, 10 ng/mL human EGF (Invitrogen, 15630-056, 35050-038, 17504-044, 17502048, PHG0311), 0.5 µg/mL hydrocortisone (Sigma, H0888-1G), 100 ng/mL human noggin (Peprotech, 120-10 C), 100 ng/mL human FGF-10 (Peprotech, 100-26-25), 100 ng/mL human FGF-7 (Peprotech, 100-19), 1.25 mM N-acetyl-L-cysteine (Sigma, A9165-5G), 2 µM TGF-β receptor kinase Inhibitor IV (SB431542, Calbiochem), 10 µM ROCK inhibitor (Y-27632) (Sigma, Y0503), 10mM nicotinamide (Sigma, N0636), 10 µM forskolin (Sigma, F6886).

For organoid generation, single epithelial cells were harvested from 2D co-culture by differential TrypLE treatment. 10,000 single epithelial cells were seeded per well of 24 well plate in 50 µL Matrigel (BD, #356231) and after polymerization of Matrigel, 500 µL of human ectocervical primary cell medium was overlaid and incubated at 37°C in a 5% $CO_2$ incubator for 6 days. Medium was changed every 2 to 3 days.

**Primary infection of organoids with HSV-GFP virus**

Day 6 grown organoids were harvested using ice-cold ADF++ medium. To determine the optimal MOI of HSV-1 (17+) Lox-pMCMV-GFP virus for automated fluorescence analysis, the viral stock was serially diluted in infection medium (ADF++ supplemented with 5% heat-inactivated FBS). Diluted virus was incubated with organoids for 2 hours at 37°C with gentle shaking.

Following incubation, organoids were centrifuged and the supernatant was discarded. The organoids were then reseeded in 50 µL Matrigel per well of a 24-well plate, overlaid with 500 µL human ectocervical primary cell medium, and incubated for 48 h at 37°C in 5% $CO_2$.

**Chemical compound treatment of organoids**
Chemical compounds were dissolved in DMSO at a stock concentration of 10 mM and further diluted in organoid medium to achieve the desired final concentrations. Day 6 organoids, either uninfected or HSV-infected, were cultured in the presence or absence of compounds for 48 hours.

**Re-infection of HSV-GFP in HeLa cells**
To assess viral re-infection ability following compound treatment, HeLa cells were infected with virus derived either from infected organoid lysates (intracellular virus) or culture supernatants (extracellular virus). For lysate preparation, organoid pellets were lysed using 3 mm glass beads by vortexing for 4 minutes in 1 mL infection medium in a 15 mL Falcon tube. Re-infected HeLa cells were incubated for 36 hours at 37°C in 5% $CO_2$.

**Microscopy**
For the analysis of cell cytotoxicity and virus infection, treated organoids were incubated with propidium Iodide (0.1 µg/mL, Sigma-Aldrich, P4170) and Hoechst 33342 (0.5 µg/mL, Merck, TA9H97BAECD2) in media for 45 minutes. Organoids were immediately imaged using an automated Keyence microscope. PI and GFP integrated intensities were normalized to DNA integrated area and expressed as percentage change relative to media-only controls.
For re-infection analysis in HeLa cells, cells were fixed overnight in methanol post-infection, stained with DAPI (1:1000) for 1 h, washed with PBS, and imaged using a Keyence microscope.

**Immunofluorescence analysis of Human tissue and organoids**
Immunofluorescence staining of paraffin-embedded tissue and organoid sections was carried out as previously described.[17,18]

**Human organoid culture, drug treatment and HSV infection**
Isolation of epithelial stem cells from the human cervix, cultivation, and maintenance in 2D and organoids were performed as described previously. Patient-derived cervical epithelial organoids were maintained in three-dimensional culture under defined conditions as previously described. For perturbation experiments, organoids were treated with DMSO (vehicle control), Imiquimod, **1** or **7** for 36 hours. For infection experiments, organoids were exposed to HSV under matched culture conditions. Uninfected organoids cultured in parallel served as controls. Biological replicates were generated for all experimental conditions.

**RNA extraction and microarray expression profiling**

**Microarray expression profiling and data analysis**
Untreated or treated organoids with or without HSV-1 (17+) Lox-pMCMVGFP virus infection were pelleted and resuspended in 1mL Trizol® reagent (Invitrogen™, 15596026) and snap-frozen in liquid nitrogen. Total RNA was measured using a NanoDrop 1000 UV–Vis spectrophotometer (Kisker), and quality was assessed by Agilent 2100 Bioanalyzer with RNA Nano 6000 microfluidics kit (Agilent Technologies, 5067-1511). All Microarray experiments were performed as single-color hybridizations. Total RNA was amplified and labeled with the Low Input Quick-Amp Labeling Kit (Agilent Technologies, 5190-2305). In brief, mRNA was reverse transcribed and amplified using an oligo-dT-T7 promoter primer and labeled with cyanine 3-CTP. After precipitation, purification, and quantification, 0.75 µg labeled cRNA was fragmented and hybridized to custom whole-genome human 8 × 60K microarrays (Agilent-048908) according to the supplier's protocol (Agilent Technologies). Microarrays were

scanned with 3 µm resolution using an Agilent G2505C microarray scanner (Agilent Technologies). Microarray image data were processed with Feature Extraction software v12.2.0.7 (Agilent Technologies) using default settings and the GE1_1200_Jun14 extraction protocol and the 048908_D_F_20200316 grid.

**Microarray preprocessing and normalization**

The extracted single-color raw data files were background corrected, and quantile normalized in R (v4.4.1) using the package limma (v.3.62.2).[19] Microarray probes were mapped to HGNC gene symbols and probes mapping to the same gene were averaged to obtain gene-level expression values using the 'avereps' function in limma. Genes lacking valid annotation were excluded from further analysis. Quality control was performed by comparison of signal intensity distributions and principal component analysis of normalized expression profiles.

**Differential expression and functional enrichment analysis**

Differential gene expression was analyzed using the R package limma.[19] Four families of comparisons were defined: (i) the effect of each compound relative to vehicle in uninfected organoids; (ii) the effect of each compound relative to vehicle in HSV-1-infected organoids; (iii) the effect of HSV-1 infection within each treatment background; and (iv) the drug x HSV-1 interaction, computed for each compound as: (HSV-1 + drug - uninfected + drug) - (HSV-1 + DMSO - uninfected + DMSO), that is, the difference between the HSV-1-induced $\log_2$ fold change in the drug background and in the vehicle background. Positive interaction values indicate that the compound amplifies the infection-driven change; negative values indicate attenuation. Differentially expressed genes (DEG) were defined at a Benjamini-Hochberg adjusted $p$-value < 0.05 and absolute log2 fold change > 0.6.

Gene Ontology over-representation analysis was performed separately for up- and downregulated genes using the R package clusterProfiler (v.4.14.6)[20] and the human annotation database org.Hs.eg.db (v.3.18.0). Significantly enriched biological processes and molecular functions were identified at FDR < 5%. Overlap between differentially expressed gene sets was assessed using the Jaccard similarity index and visualized using UpSetR (v.1.4.0).[21] Correlations between transcriptional responses across comparisons were determined using Pearson correlation of log2 fold changes.
Pre-ranked gene set enrichment analysis was performed using the R package fgsea (v.1.35.6) with gene sets obtained from the Molecular Signatures Database (MSigDB) using msigdbr (v.7.5.1) (https://igordot.github.io/msigdbr/). Genes were ranked according to the direction and magnitude of differential expression together with their statistical significance. Hallmark and selected MSigDB pathway collections, including Gene Ontology biological processes and regulatory gene sets, were used for enrichment analysis. Gene sets with an adjusted p-value < 0.05 were considered significantly enriched.
Additional gene set collections including GO biological process (C5:GO:BP), Reactome (C2:CP:REACTOME), KEGG (C2:CP:KEGG), and transcription factor target (C3:TFT) gene sets were also retrieved from MSigDB via the msigdbr package. For visualization of pathway activity across individual samples, gene set module scores were calculated as the mean z-scored expression of detected genes belonging to each gene set. Gene sets represented by fewer than five genes in the expression dataset were excluded.

**Transcription factor (TF) activity inference**

Transcription factor activity was inferred from normalized gene expression using DoRothEA regulons of confidence levels A-C and the VIPER algorithm. Human regulons were obtained using DoRothEA (v.1.18.0),[22] and TF activity scores were computed using VIPER (v.1.40.0) package.[23] VIPER normalized enrichment scores (NES) were calculated for each TF in each sample. Differential TF activity between experimental conditions was analyzed using the same linear modeling and contrast framework as for gene expression analysis, with adjusted p-value < 0.05 considered significant.

**TF activity restoration analysis**

To determine whether drug treatment restored HSV-induced TF activity toward the uninfected state, TF activity in HSV-infected drug-treated organoids was compared with the corresponding drug-matched uninfected condition. For each TF, displacement from the drug-matched uninfected baseline was calculated as the absolute difference in inferred VIPER NES between the HSV-infected vehicle condition and the uninfected drug-matched condition (x-axis displacement), and between the HSV-infected drug condition and the uninfected drug condition (y-axis displacement). Restoration was defined as movement toward the uninfected baseline and was considered biologically relevant when the reduction in displacement exceeded ($|\Delta| > 0.3$) and the fractional restoration exceeded 20% (> 0.20), enabling us to distinguish substantial from small TF shifts.

#### 1.3. Synthesis and characterization

***General details.*** Reactions were conducted in common laboratory glassware under magnetic stirring. Room temperature (RT) is defined as 20–25 °C. Reactions requiring heating were carried out using heating blocks, with temperatures monitored by a contact thermocouple. Operations under reduced pressure utilized a Büchi Rotavapor R-100, equipped with a Büchi V-491 heating bath and a Büchi V-800/850 vacuum controller.

##### Synthesis of complex 1

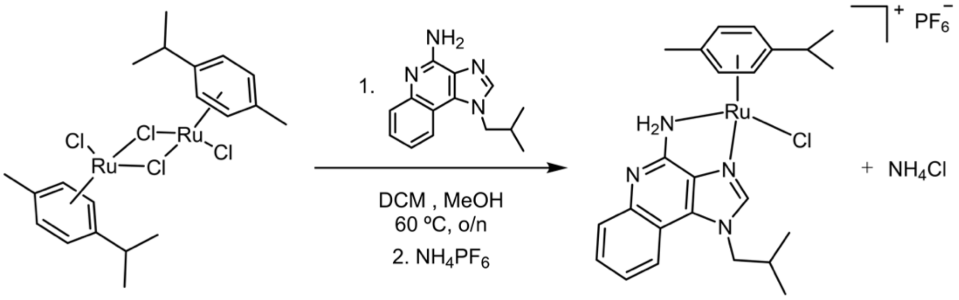


**Scheme 1.** Synthesis of complex **1**.

**[Ru(η⁶-p-cym)(imq)(Cl)]$PF_6$ (1).** (p-cymene)ruthenium(II) chloride dimer, [Ru(*p*-cymene)$Cl_2$]$_2$ (100.0 mg, 0.163 mmol, 1.0 eq) was dissolved in DCM (2.5 mL) under magnetic stirring. Then, a solution of imiquimod (78.72 mg, 0.328 mmol, 2.0 eq) in MeOH (20 mL) was added dropwise. The reaction mixture was left to stir at reflux (58–60 °C) overnight (16–24 h). After this time a solution colour change from dark red to bright orange was observed and an excess of $NH_4PF_6$ (140.1 mg, 0.859 mmol, 5.0 eq) was added before reducing to half-volume the amount of solvent under reduced pressure. Then, the solution was left to crystallize at 4 °C for approximately 24 h. The crystalline solid was isolated by decanting the supernatant, washed three times with diethyl ether, and dried under reduced pressure. The title was obtained as a crystalline dark orange solid (109.3 mg, 96.9%). Crystals suitable for X-ray diffraction were obtained by crystallization in methanol at 298 K.
**$^1$H NMR** (300 MHz, Methanol-*d*$_4$) δ 8.81 (s, 1H), 7.87 (d, *J* = 8.1 Hz, 1H), 7.54 – 7.45 (m, 1H), 7.40 – 7.28 (m, 2H), 5.89 (dd, *J* = 13.2, 6.0 Hz, 2H), 5.64 (dd, *J* = 8.4, 6.1 Hz, 2H), 4.42 (d, *J* = 7.4 Hz, 2H), 2.79 (hept, *J* = 6.8 Hz, 1H), 2.30 – 2.15 (m, 1H), 2.23 (s, 3H), 1.20 (dd, *J* = 9.1, 6.9 Hz, 6H), 1.03 (d, *J* = 6.7 Hz, 6H). **$^{13}$C NMR** (75 MHz, Methanol-*d*$_4$) δ 153.47, 146.81, 139.20, 136.46, 131.58, 130.83, 124.71, 123.19, 118.61, 113.71, 103.61, 100.77, 83.55, 83.26, 82.17, 81.05, 56.30, 32.47, 30.47, 22.62, 22.33, 19.70, 19.65, 19.03. **MS** (ESI+): m/z calcd para $C_{24}H_{30}ClN_4Ru$ [M]$^+$: 511.1197; found: 511.1166. **EA** calcd for $C_{24}H_{30}ClF_6N_4PRu \cdot H_2O$: C 42.77, H 4.79, N 2.59; found C 42.50, H 459, N 8.49.

##### Synthesis of complexes 2–6

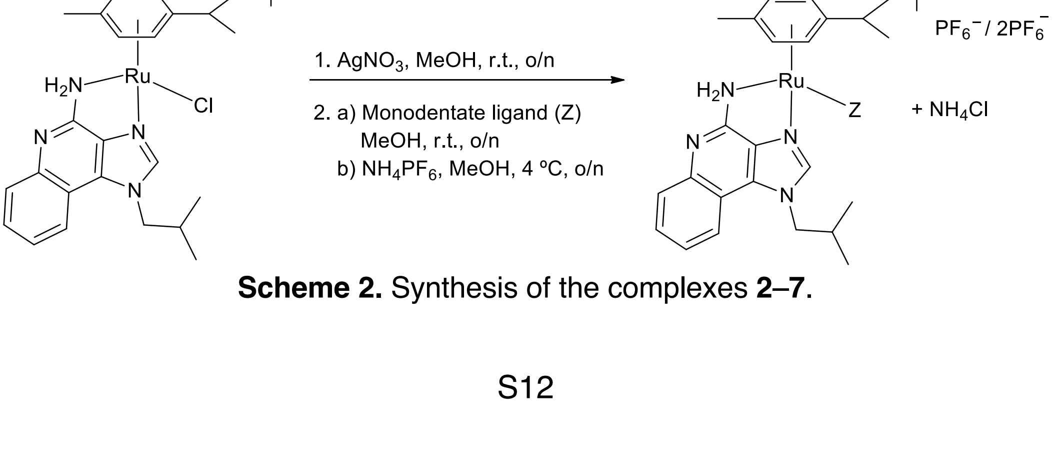


**Scheme 2.** Synthesis of the complexes **2–7**.

*General procedure.* A solution of compound **1** (1.0 eq) and $AgNO_3$ in methanol was stirred in a round-bottom flask, protected from light, at room temperature (rt) overnight. After that time, the reaction mixture was filtered using a syringe filter (0.2 µm) to remove the precipitated silver salts (AgCl). The resulting solution was transferred to another round-bottom flask, and the corresponding monodentate ligand (1.2-10.0 eq) was added. The reaction mixture was stirred overnight at rt. Subsequently, an excess of $NH_4PF_6$ (5.0 eq) was added, and the volume was reduced to one-fourth of its original volume under reduced pressure using a rotary evaporator. This solution was allowed to crystallize at 4°C for approximately 24 hours. The solid was isolated by decanting the supernatant, washed three times with diethyl ether, and dried under reduced pressure.

**[Ru(η$^6$-p-cym)(imq)(py)]($PF_6$)$_2$ (2).** Following the general procedure, the synthesis of complex **2** was carried out using compound **1** (29.5 mg, 0.045 mmol), $AgNO_3$ (9.3 mg, 0.055 mmol), pyridine (py) (37.0 µL, 0.450 mmol) as the monodentate ligand, and $NH_4PF_6$ (40.0 mg, 0.245 mmol). The compound was obtained as a yellow solid (30.0 mg, 78.9% yield). Crystals suitable for X-ray diffraction were obtained by crystallization in methanol at 298 K.
**$^1$H NMR** (300 MHz, Methanol-*d*$_4$) δ 9.12 (s, 1H), 8.59 (d, *J* = 5.0 Hz, 2H), 7.95 – 7.87 (m, 2H), 7.60 – 7.52 (m, 1H), 7.49 – 7.34 (m, 4H), 6.21 (dd, *J* = 27.0, 6.4 Hz, 2H), 5.88 (dd, *J* = 14.9, 6.0 Hz, 2H), 4.50 (dd, *J* = 7.6, 4.2 Hz, 2H), 2.63 (hept, *J* = 13.9, 6.7 Hz, 1H), 2.24 (hept, *J* = 13.3, 6.5 Hz, 1H), 1.86 (s, 3H), 1.13 – 1.03 (m, 9H), 0.97 (d, *J* = 6.6 Hz, 3H). **$^{13}$C NMR** (126 MHz, Methanol-*d*$_4$) δ 155.08, 154.02, 147.88, 140.92, 139.12, 135.70, 132.50, 131.48, 127.64, 125.35, 123.44, 119.12, 113.42, 105.15, 104.09, 87.29, 86.15, 83.94, 81.79, 56.72, 32.40, 30.25, 22.71, 22.37, 19.62, 18.06. **MS** (ESI+): m/z calcd for $C_{29}H_{35}N_5Ru$ [M – py – H]$^+$: 475.1430; found: 475.1440. **EA** calcd for $C_{29}H_{35}F_{12}N_5P_2Ru{\cdot}2H_2O$: C 39.55, H 4.46, N 7.95; found C 39.32, H 4.19, N 8.25.

**[Ru(η$^6$-p-cym)(imq)(pta)]($PF_6$)$_2$ (3).** Following the general procedure, the synthesis of complex **3** was carried out using compound **1** (29.5 mg, 0.045 mmol), $AgNO_3$ (7.1 mg, 0.055 mmol), pta (8.9 mg, 0.054 mmol) as the monodentate ligand, and $NH_4PF_6$ (39.2 mg, 0.225 mmol). The compound
was obtained as a yellow solid (36.5 mg, 87.9% yield). Crystals suitable for X-ray diffraction were obtained by crystallization in methanol at 298 K.
**$^1$H NMR** (400 MHz, Acetone-*d*$_6$) δ 8.86 (s, 1H), 8.16 (d, *J* = 9.0 Hz, 1H), 7.66 (ddd, *J* = 8.5, 7.2, 1.3 Hz, 1H), 7.58 (dd, *J* = 8.4, 1.2 Hz, 1H), 7.48 (ddd, *J* = 8.3, 7.2, 1.3 Hz, 1H), 6.31 (d, *J* = 6.2 Hz, 1H), 6.28 – 6.18 (m, 2H), 6.17 (d, *J* = 6.2 Hz, 1H), 4.65 (dd, *J* = 7.6, 2.5 Hz, 2H), 4.43 (s, 6H), 4.20 (s, 6H), 2.95 (hept, *J* = 7.1 Hz, 2H), 2.41 (s, 3H), 1.33 (d, *J* = 6.9 Hz, 3H), 1.29 (d, *J* = 6.9 Hz, 3H), 1.09 (dd, *J* = 6.6, 2.0 Hz, 6H). *A multiplet corresponds to the CH signal of imiquimod below the singlet at 2.41 ppm.* **$^{13}$C NMR** (126 MHz, Acetone-*d*$_6$) δ 206.19, 152.74, 148.78, 138.71, 136.44, 132.98, 131.34, 125.20, 123.53, 119.00, 113.65, 112.52, 104.14, 89.34, 89.32, 89.28, 89.25, 88.83, 88.81, 87.19, 87.16, 72.96 (d, $J_{PC}$ = 7.7 Hz), 56.39, 52.04 (d, $J_{PC}$ = 14.9 Hz), 32.08, 23.03, 22.99, 19.73, 19.66, 19.11. **$^{31}$P NMR** (126 MHz, Acetone-*d*$_6$) δ -33.55 (s, 1P, pta), -143.57 (hept, $J_{PF}$ = 708.8 Hz, $PF_6$) **MS** (ESI+): m/z calcd for $C_{30}H_{42}N_7PRu$ [M – H]$^+$: 632.2199; found: 632.2211. **EA** calcd for $C_{30}H_{42}F_{12}N_7P_3Ru{\cdot}0.5NH_4PF_6$: C, 35.88; H, 4.42; N, 10.46; found: C, 35.79; H, 4.47; N, 10.27.

**[Ru(η$^6$-p-cym)(imq)(im)]($PF_6$)$_2$ (4).** Following the general procedure, the synthesis of complex **4** was carried out using compound **1** (30 mg, 0.046 mmol), $AgNO_3$ (7.4 mg, 0.043 mmol), imidazole (3.7 mg, 0.0543 mmol) as the monodentate ligand and $NH_4PF_6$ (36.7 mg, 0.225 mmol). The compound was obtained as a light-yellow solid (35.2 mg, 93.8%). Crystals suitable for X-ray diffraction were obtained by crystallization in methanol at 298 K.

$^{1}$**H NMR** (300 MHz, Methanol-*d*$_4$) δ 8.99 (s, 1H), 7.90 (d, *J* = 8.1 Hz, 1H), 7.83 (s, 1H), 7.54 (t, *J* = 7.8 Hz, 1H), 7.45 (d, *J* = 8.3 Hz, 1H), 7.36 (t, *J* = 7.6 Hz, 1H), 7.09 (s, 1H), 6.97 (s, 1H), 6.19 (d, *J* = 5.9 Hz, 1H), 6.08 (d, *J* = 6.0 Hz, 1H), 5.84 (dd, *J* = 10.0, 6.2 Hz, 2H), 4.46 (dd, *J* = 7.6, 3.8 Hz, 2H), 2.61 (hept, *J* = 6.5 Hz, 1H), 2.21 (hept, *J* = 7.4 Hz, 1H), 1.90 (s, 3H), 1.15 – 1.01 (m, 10H), 0.96 (d, *J* = 6.6 Hz, 3H). $^{13}$**C NMR** (75 MHz, Methanol-*d*$_4$) δ 153.83, 147.64, 139.77, 139.12, 135.86, 132.19, 131.28, 130.62, 125.13, 123.33, 119.54, 118.97, 113.44, 111.39, 105.20, 103.55, 86.69, 85.45, 83.19, 81.15, 56.55, 32.42, 30.27, 22.76, 22.40, 19.62, 19.57, 18.26. **MS** (ESI+): m/z calcd for $C_{27}H_{34}N_6Ru$ [M – H]$^+$: 543.1810; found: 543.1826. **EA** calcd for $C_{27}H_{34}F_{12}N_6P_2Ru \cdot 2H_2O$: C 37.29, H 4.40, N 9.66; found C 37.11, H 4.00, N 9.87.

**[Ru(η$^6$-p-cym)(imq)(in)](PF$_6$)$_2$ (5).** Following the general procedure, the synthesis of complex **5** was carried out using compound **1** (20.0 mg, 0.030 mmol), $AgNO_3$ (4.7 mg, 0.028 mmol), indazol (5.32 mg, 0.045 mmol) as monodentate ligand and $NH_4PF_6$ (24.3 mg, 0.150 mmol). The compound was obtained as a light-yellow solid (26.5 mg, 87.5%).
$^{1}$**H NMR** (300 MHz, Methanol-*d*$_4$) δ 9.04 (s, 1H), 8.19 (s, 1H), 7.88 (d, *J* = 8.2 Hz, 1H), 7.68 (d, *J* = 8.3 Hz, 1H), 7.59 – 7.42 (m, 4H), 7.41 – 7.32 (m, 1H), 7.17 (t, *J* = 7.5 Hz, 1H), 6.36 (d, *J* = 5.8 Hz, 1H), 6.22 (d, *J* = 6.0 Hz, 1H), 6.00 (t, *J* = 6.1 Hz, 2H), 4.47 (d, *J* = 7.6 Hz, 2H), 2.69 – 2.51 (m, 1H), 2.32 – 2.10 (m, 1H), 1.91 (s, 3H), 1.13 (d, *J* = 6.9 Hz, 3H), 1.06 (d, *J* = 6.9 Hz, 3H), 1.02 (d, *J* = 6.6 Hz, 3H), 0.94 (d, *J* = 6.6 Hz, 3H). $^{13}$**C NMR** (75 MHz, Methanol-*d*$_4$) δ 148.07, 143.28, 140.40, 139.02, 135.67, 132.56, 131.46, 130.35, 125.38, 124.13, 123.70, 123.45, 121.53, 119.08, 113.42, 111.40, 110.79, 105.61, 104.27, 87.25, 86.07, 84.01, 81.94, 56.67, 32.50, 30.23, 22.77, 22.23, 19.55, 18.42. **MS** (ESI+): m/z calcd for $C_{31}H_{36}N_6Ru$ [M – H]$^+$: 593.1961; found: 593.1966. **EA** calcd for $C_{31}H_{36}F_{12}N_6P_2Ru \cdot 0.5NH_4PF_6$: C, 38.58; H, 3.97; N, 9.43; found: C, 38.81; H, 4.06; N, 9.68.

**[Ru(η$^6$-p-cym)(imq)(5-fu)]PF$_6$ (6).** Following the general procedure, the synthesis of complex **6**, was carried out using compound **1** (50.0 mg, 0.076 mmol), $AgNO_3$ (12.1 mg, 0.072 mmol) and the potassium salt of 5-fluorouracil (5-KHFU, 15.21 mg, 0.0812 mmol) as monodentate ligand. The compound was obtained as a yellow solid (33.3 mg, 82.7%).
Modifying the procedure described by X. Wang et al.,[24] the potassium 5-KHFU salt was also obtained by reacting equimolar amounts of 5-FU (100.0 mg, 0.768 mmol) and KOH (46.2 mg, 0.823 mmol) in MeOH (2 mL) under magnetic stirring at room temperature for 2 hours. After this time, a crystalline white solid was obtained, filtered, and washed three times with cold MeOH (~ 4 °C). Once dried under reduced pressure, it was ready for use without further purification.
$^{1}$**H NMR** (300 MHz, Acetone-*d*$_6$) δ 8.99 (s, 1H), 7.92 (d, *J* = 6.8 Hz, 1H), 7.56 – 7.42 (m, 2H), 7.32 (t, *J* = 7.2 Hz, 2H), 6.19 (d, *J* = 5.0 Hz, 1H), 6.01 (d, *J* = 6.0 Hz, 1H), 5.84 (t, *J* = 4.8 Hz, 2H), 4.59 – 4.29 (m, 2H), 2.90 – 2.74 (m, 1H), 2.20 (s, 3H), 2.18 – 2.10 (m, 1H), 1.19 (d, *J* = 6.9 Hz, 3H), 1.13 (d, *J* = 6.9 Hz, 3H), 0.90 (d, *J* = 6.6 Hz, 3H), 0.78 (d, *J* = 6.6 Hz, 3H). $^{13}$**C NMR** (75 MHz, Acetone-*d*$_6$) δ 158.92 (d, $J_{FC}$ = 25.2 Hz), 157.55, 152.95, 148.69, 142.82 (d, $J_{FC}$ = 225.9 Hz), 140.83 (d, $J_{FC}$ = 26.5 Hz), 138.50, 135.70, 130.95, 130.66, 124.72, 123.12, 118.75, 113.55, 104.47, 102.09, 84.76, 84.08, 83.89, 81.24, 55.76, 31.90, 22.99, 22.24, 19.47, 19.34, 18.76. *A multiplet corresponds to the CH signal of imiquimod below the singlet at 2.41 ppm.* $^{19}$**F NMR** (376 MHz, Methanol-*d*$_4$) δ 175.17 (d, $J_{FH}$ = 6.1 Hz). **MS** (ESI+): m/z calcd for $C_{28}H_{32}FN_6O_2Ru$ [M]$^+$ 605.1609; found 605.1621. **EA** calcd for $C_{28}H_{32}F_7N_6O_2PRu \cdot 2H_2O$: C 42.81, H 4.62, N 10.7; found C 42.36, H 4.27, N 10.84.

**Synthesis of complex 7**

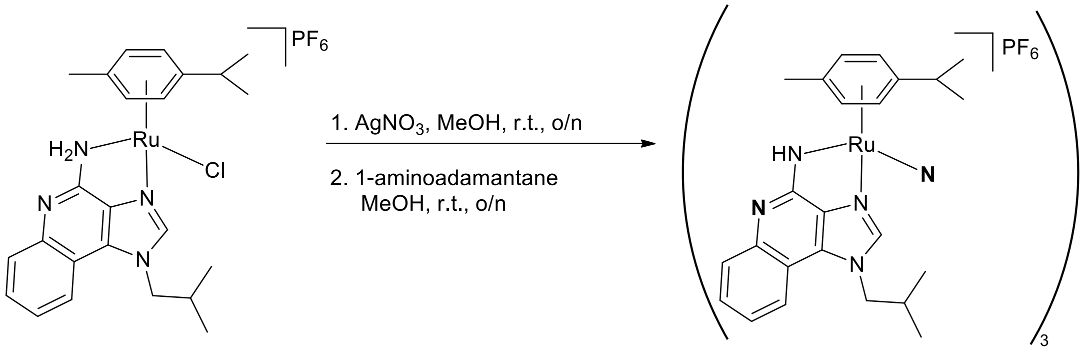


**Scheme 3.** Synthesis of complex **7**.

**[Ru(η$^6$-p-cym)(imq)]$_3$(PF$_6$)$_3$.** In a round-bottom flask, compound **1** (23.0 mg, 0.035 mmol) is dissolved in methanol (MeOH) under magnetic stirring. Next, a solution of silver nitrate ($AgNO_3$, 5.6 mg, 0.033 mmol) is added, and the reaction is stirred, protected from light at room temperature for 16–24 hours. After that time, the reaction mixture is filtered using a syringe filter (0.2 µm) to remove the silver salts (AgCl). The resulting solution is transferred to another round-bottom flask, and 1-aminoadamantane ligand (8.0 mg, 0.053 mmol) is added. The reaction mixture changed from orange to red instantaneously and, after 1 h stirring started to precipitated. The reaction mixture is left to stir for an additional 1.5 h and then is kept in the fridge overnight. The bright orange solid is isolated by decanting the supernatant, washed three times with diethyl ether, and dried under reduced pressure (9.2 mg, 34%). Crystals suitable for X-ray diffraction were obtained by crystallization in methanol at 298 K.
**$^1$H NMR** (300 MHz, Acetone-*d*$_6$) δ 9.28 (s, 1H), 8.94 (d, *J* = 8.9 Hz, 1H), 7.63 – 7.50 (m, 2H), 7.06 (t, *J* = 8.1 Hz, 1H), 6.92 (d, *J* = 7.0 Hz, 1H), 6.79 (d, *J* = 5.9 Hz, 1H), 6.71 (d, *J* = 5.7 Hz, 1H), 5.97 (d, *J* = 5.8 Hz, 1H), 4.37 (dd, *J* = 14.4, 6.4 Hz, 1H), 4.18 (dd, *J* = 14.4, 8.6 Hz, 1H), 1.98 (s, 3H), 1.96 – 1.89 (m, 1H), 1.07 (dd, *J* = 10.0, 7.0 Hz, 6H), 0.91 (d, *J* = 6.7 Hz, 3H), 0.49 (d, *J* = 6.6 Hz, 3H). *A multiplet corresponding to a C*H *p-cym is under the residual solvent peak.* **$^{13}$C NMR** (75 MHz, Acetone-*d*$_6$) δ 160.51(*C*), 147.88(*C*H), 147.81(*C*), 135.27(*C*), 130.01(*C*), 128.62(*C*H), 128.19(*C*H), 122.45(*C*H), 122.05(*C*H), 116.04(*C*), 88.63(*C*H), 86.08(*C*H), 84.68(*C*H), 79.31(*C*H), 56.08(*C*H$_2$), 32.36(*C*H), 29.41(*C*H, inside residual peak), 22.75(2x*C*H$_3$), 19.58(2x*C*H$_3$), 19.10(*C*H$_3$). *Two missing quaternary carbons are located between 100–106 ppm, assigned by correlations on $^1$H–$^{13}$C HMBC experiment.* **MS** (ESI+): m/z calcd for $C_{72}H_{86}N_{12}Ru_3$ $[M]^{3+}$ 475.1412; found 475.1437. **EA** calcd for $C_{72}H_{87}F_{18}N_{12}P_3Ru_3$: C 46.83, H 4.72, N 9.04; found C 46.21, H 4.76, N 9.59.

#### 2. Figures

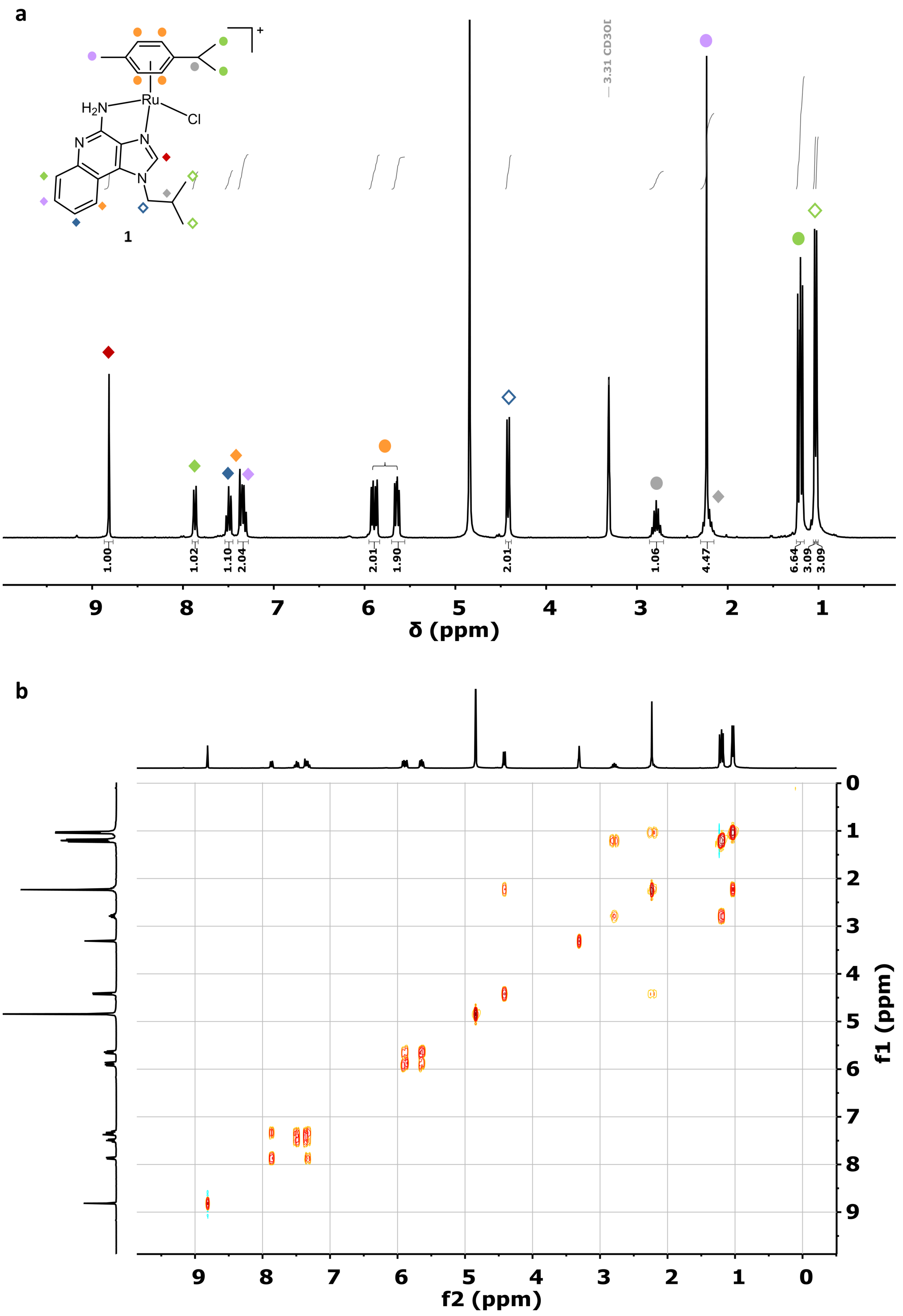


**Figure S1.** NMR spectra of complex [Ru(η[6]-*p*-cym)(imq)(Cl)]($PF_6$) (**1**). (a) [1]H (300 MHz, methanol-*d*$_4$, 298.5 K) and (b) [1]H-[1]H COSY (300 MHz, methanol-*d*$_4$, 298.5 K).

a

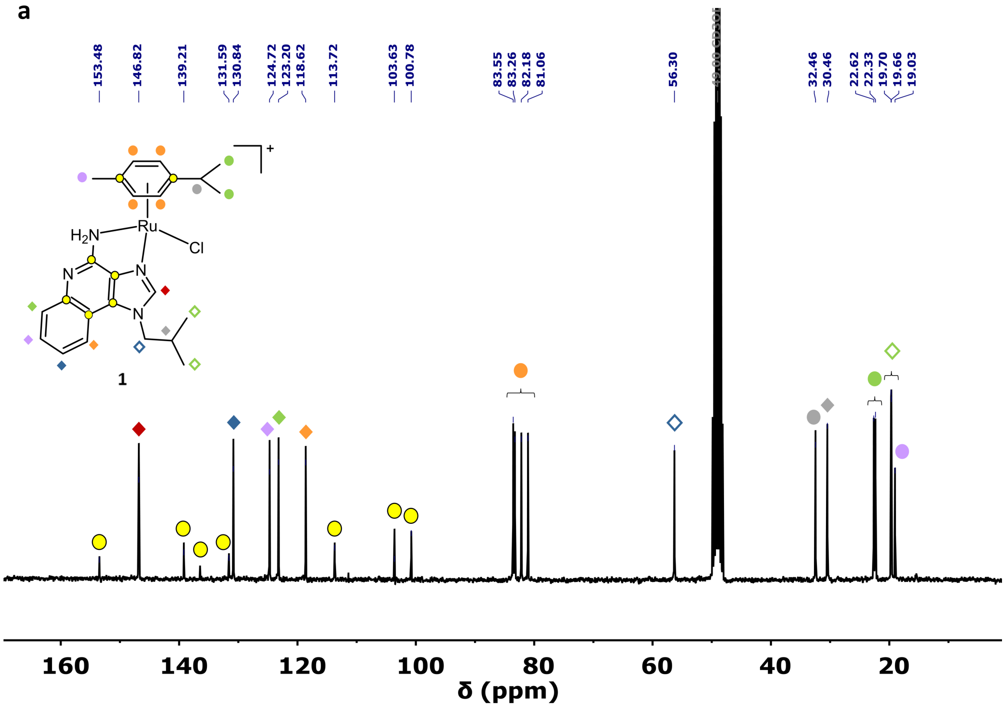

b

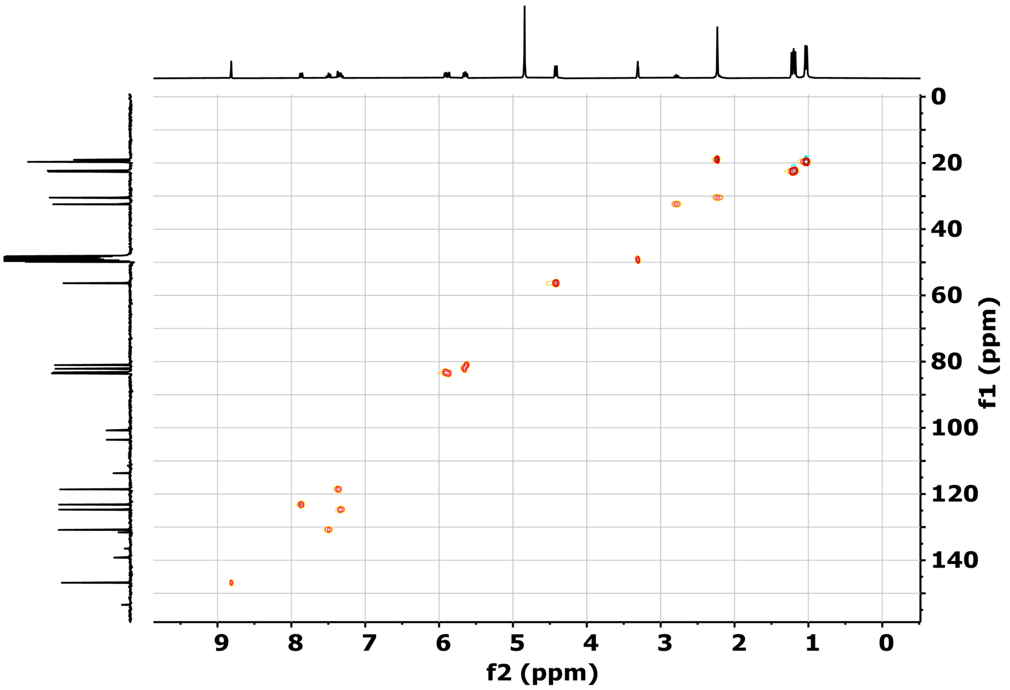

**Figure S2.** NMR spectra of complex [Ru(η$^6$-*p*-cym)(imq)(Cl)](PF$_6$) (**1**). (a) $^{13}$C (75 MHz, methanol-*d*$_4$, 298.5 K) and (b) $^1$H-$^{13}$C HSQC (300 and 75 MHz, methanol-*d*$_4$, 298.5 K).

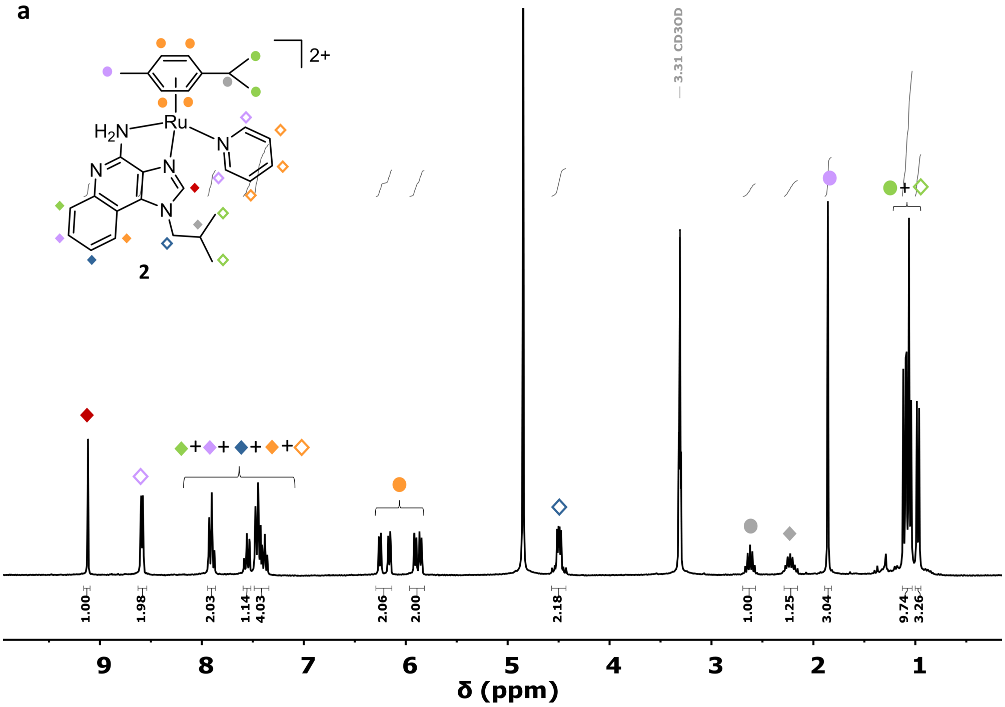


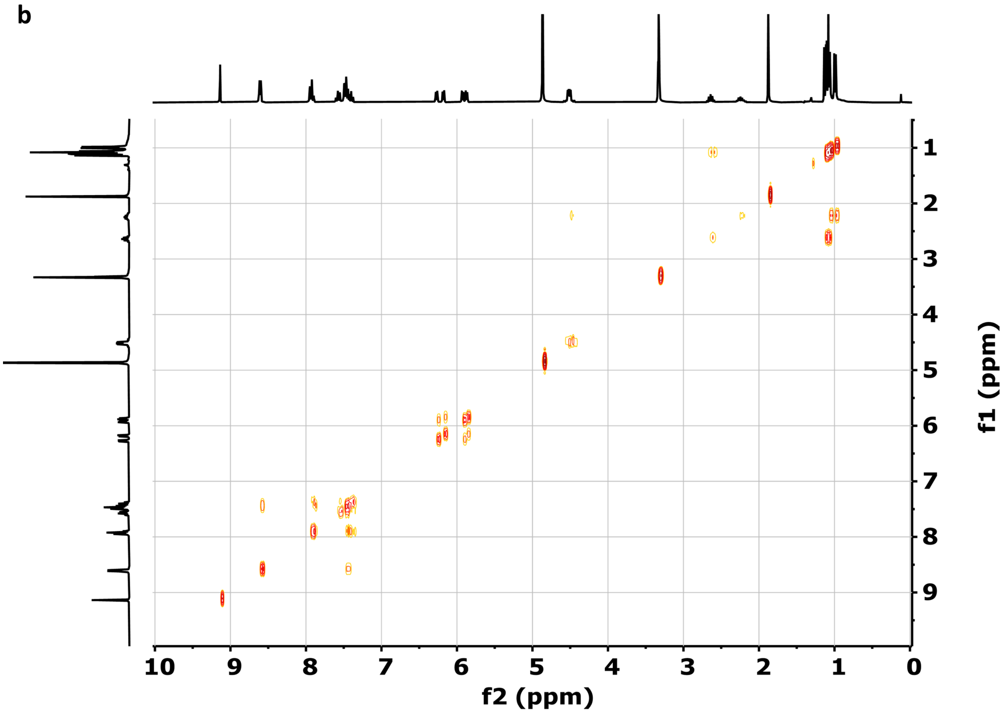


**Figure S3.** NMR spectra of complex [Ru($\eta^6$-*p*-cym)(imq)(py)]($PF_6$)$_2$ (**2**). (a) $^1$H (300 MHz, methanol-$d_4$, 298.5 K) and (b) $^1$H-$^1$H COSY (300 MHz, methanol-$d_4$, 298.5 K).

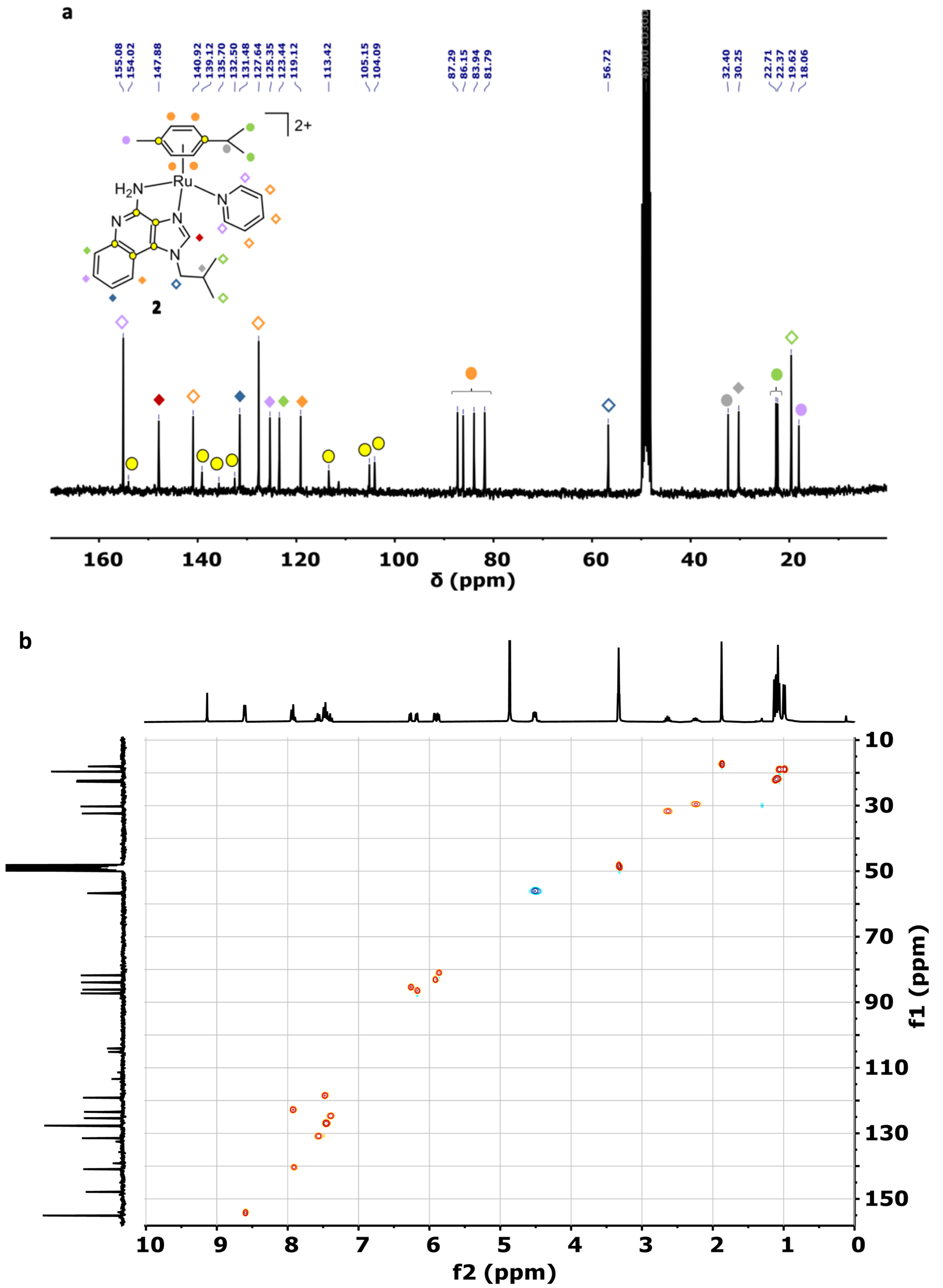


**Figure S4.** NMR spectra of complex [Ru(η$^6$-*p*-cym)(imq)(py)](PF$_6$)$_2$ (**2**). (a) $^{13}$C (75 MHz, methanol-$d_4$, 298.5 K) and (b) $^1$H-$^{13}$C HSQC (300 and 75 MHz, methanol-$d_4$, 298.5 K).

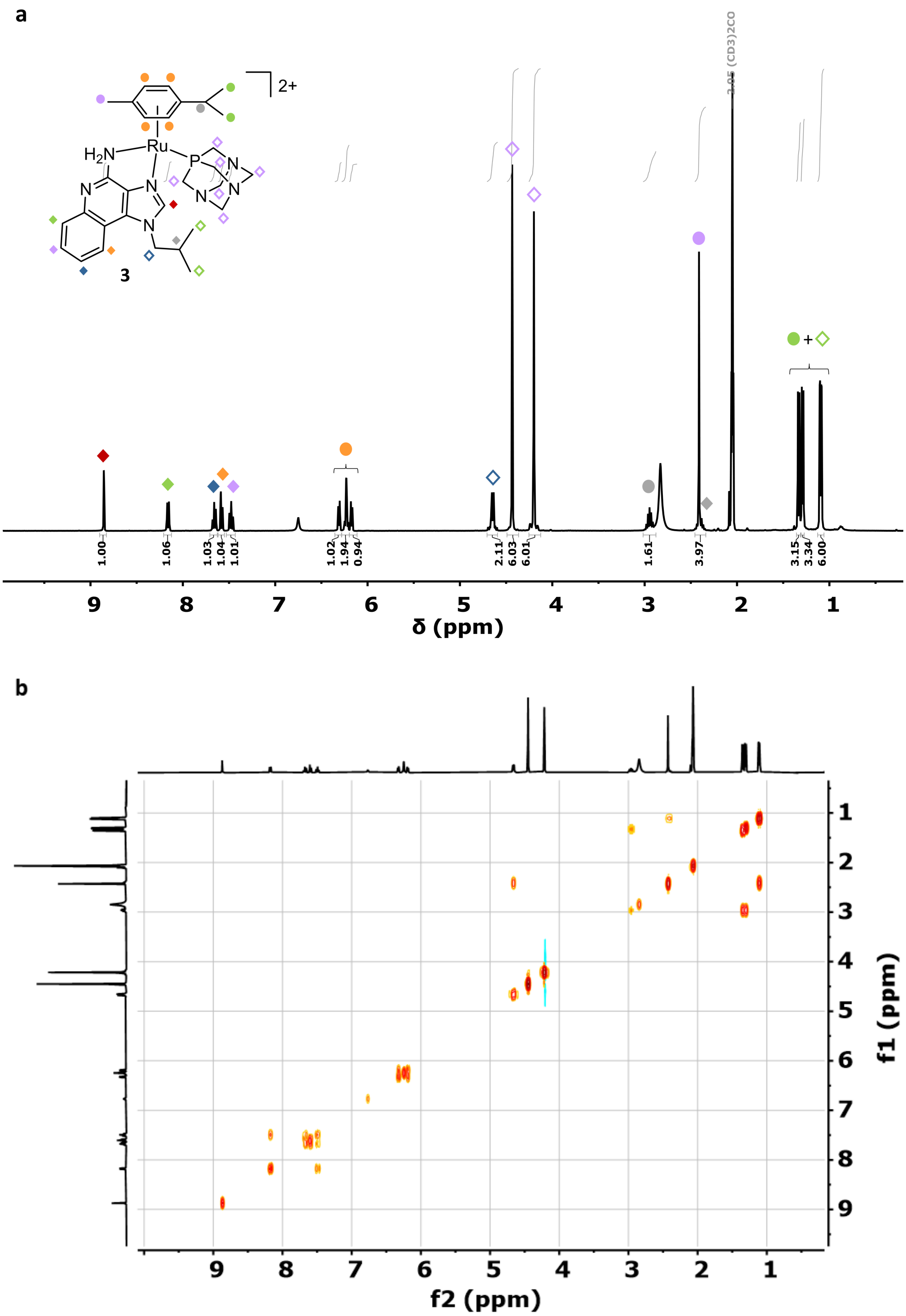


**Figure S5.** NMR spectra of complex [Ru($\eta^6$-*p*-cym)(imq)(pta)]$(PF_6)_2$ (**3**). (a) $^1H$ (400 MHz, acetone-$d_6$, 298.5 K) and (b) $^1H$-$^1H$ COSY (400 MHz, acetone-$d_6$, 298.5 K).

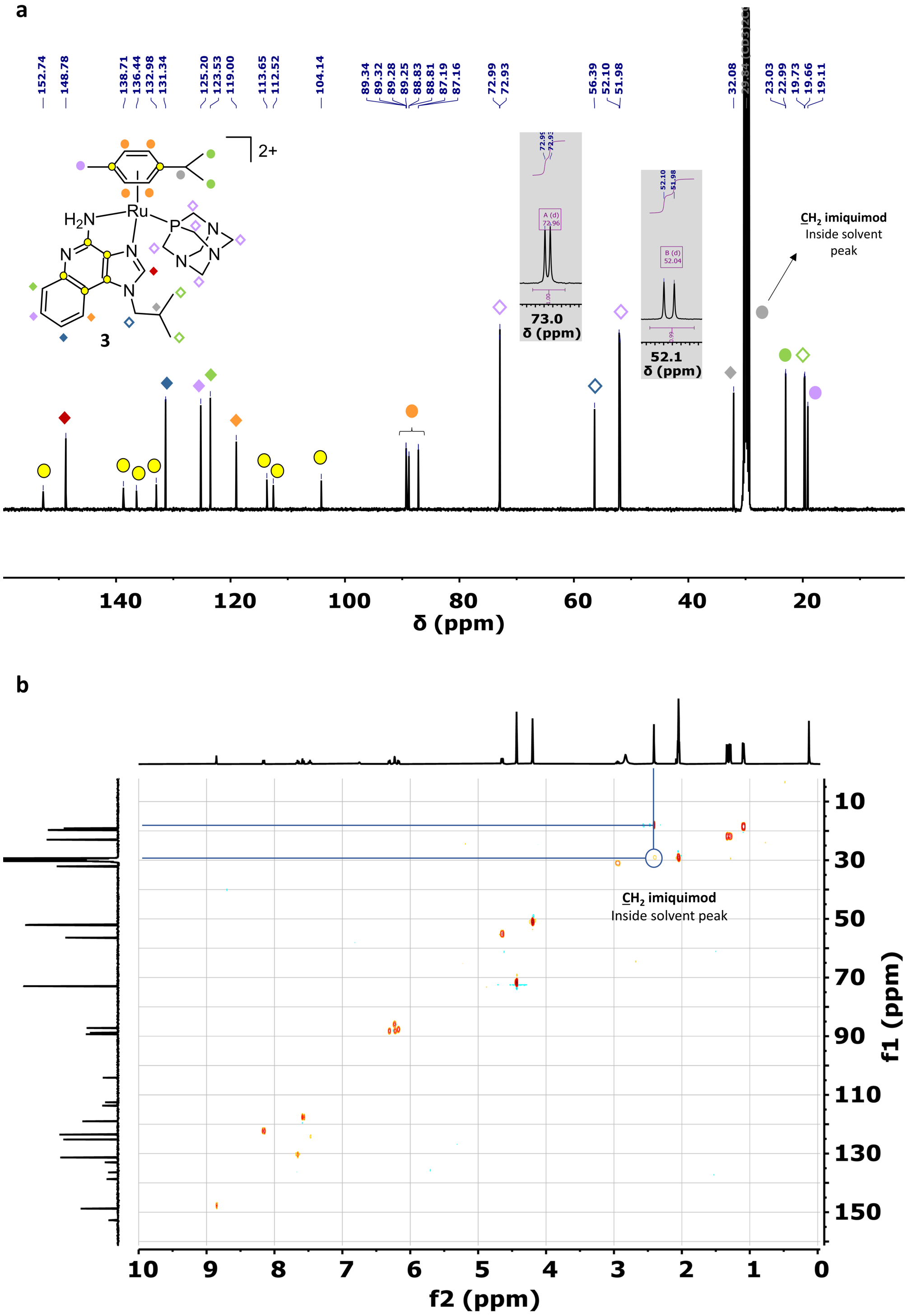


**Figure S6.** NMR spectra of complex [Ru(η$^6$-*p*-cym)(imq)(pta)]$(PF_6)_2$ (**3**). (a) $^{13}C$ (126 MHz, acetone-$d_6$, 298.5 K) and (b) $^1H$-$^{13}C$ HSQC (400 and 126 MHz, acetone-$d_6$, 298.5 K).

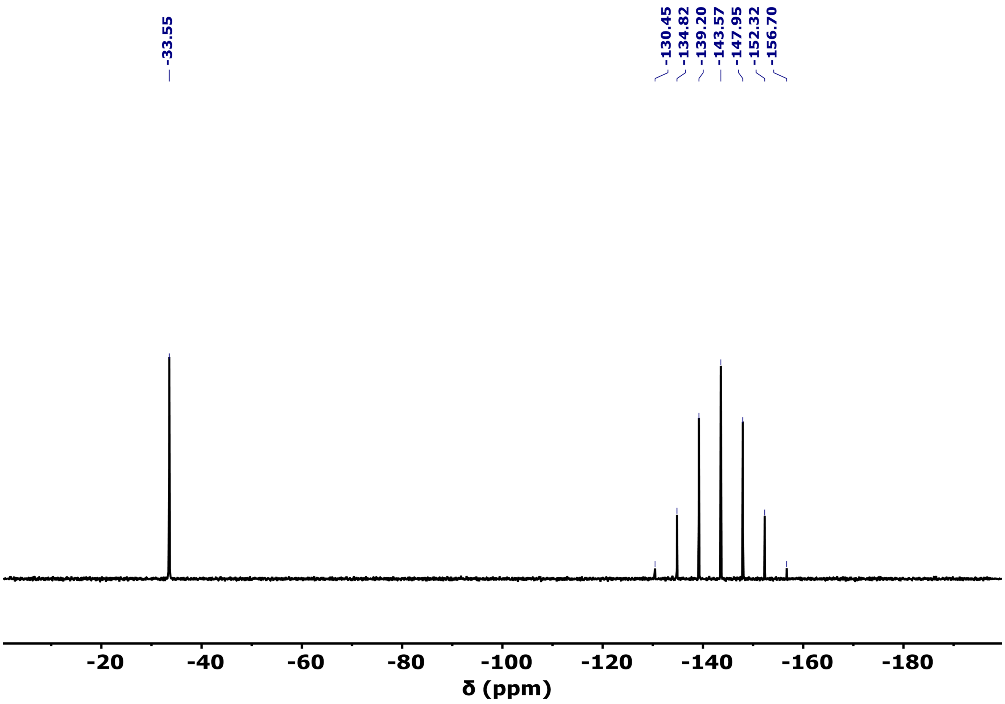


**Figure S7.** $^{31}P$ NMR spectrum of complex [Ru(η$^6$-*p*-cym)(imq)(pta)]$(PF_6)_2$ (**3**) (162 MHz, acetone-$d_6$, 298.5 K)

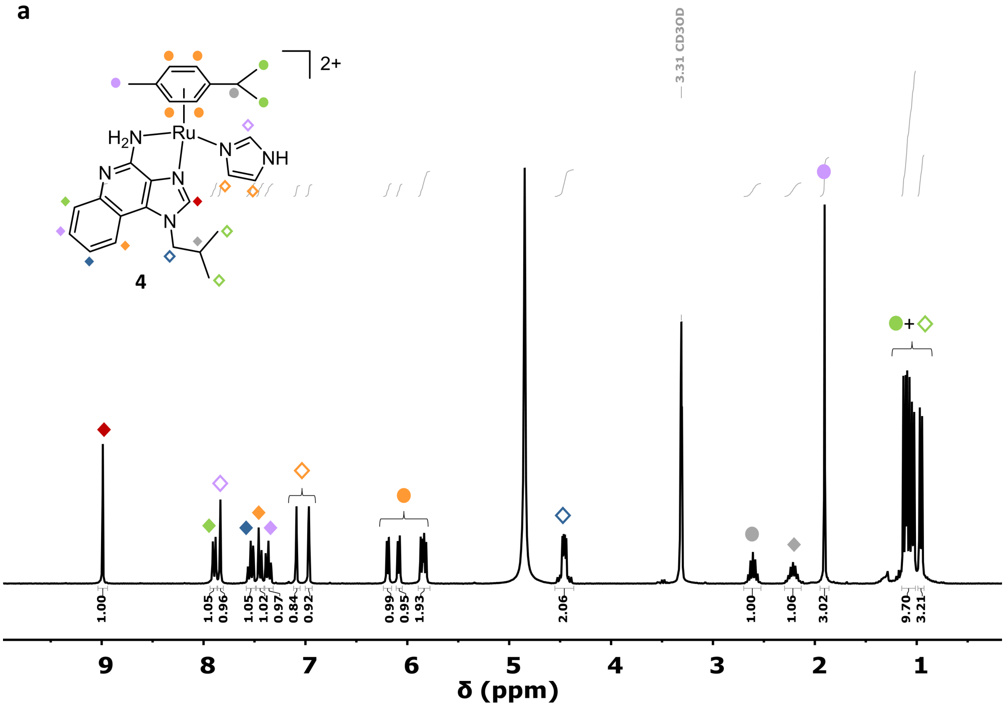


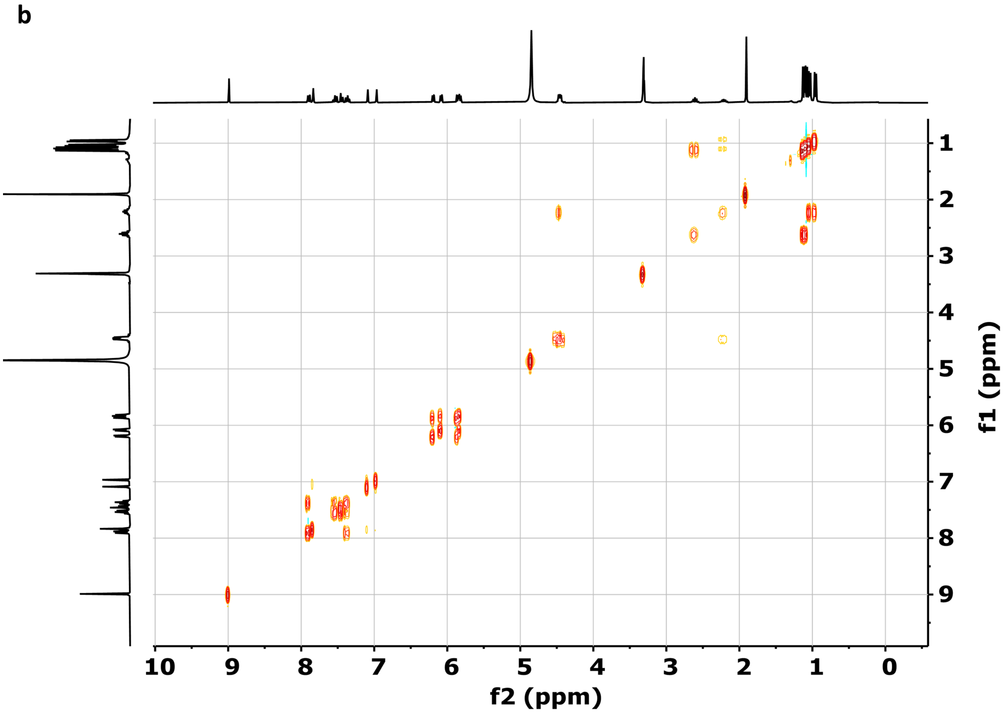


**Figure S8.** NMR spectra of complex [Ru(η$^6$-*p*-cym)(imq)(im)]($PF_6$)$_2$ (**4**). (a) $^1$H (300 MHz, methanol-$d_4$, 298.5 K) and (b) $^1$H-$^1$H COSY (300 MHz, metanol-$d_4$, 298.5 K).

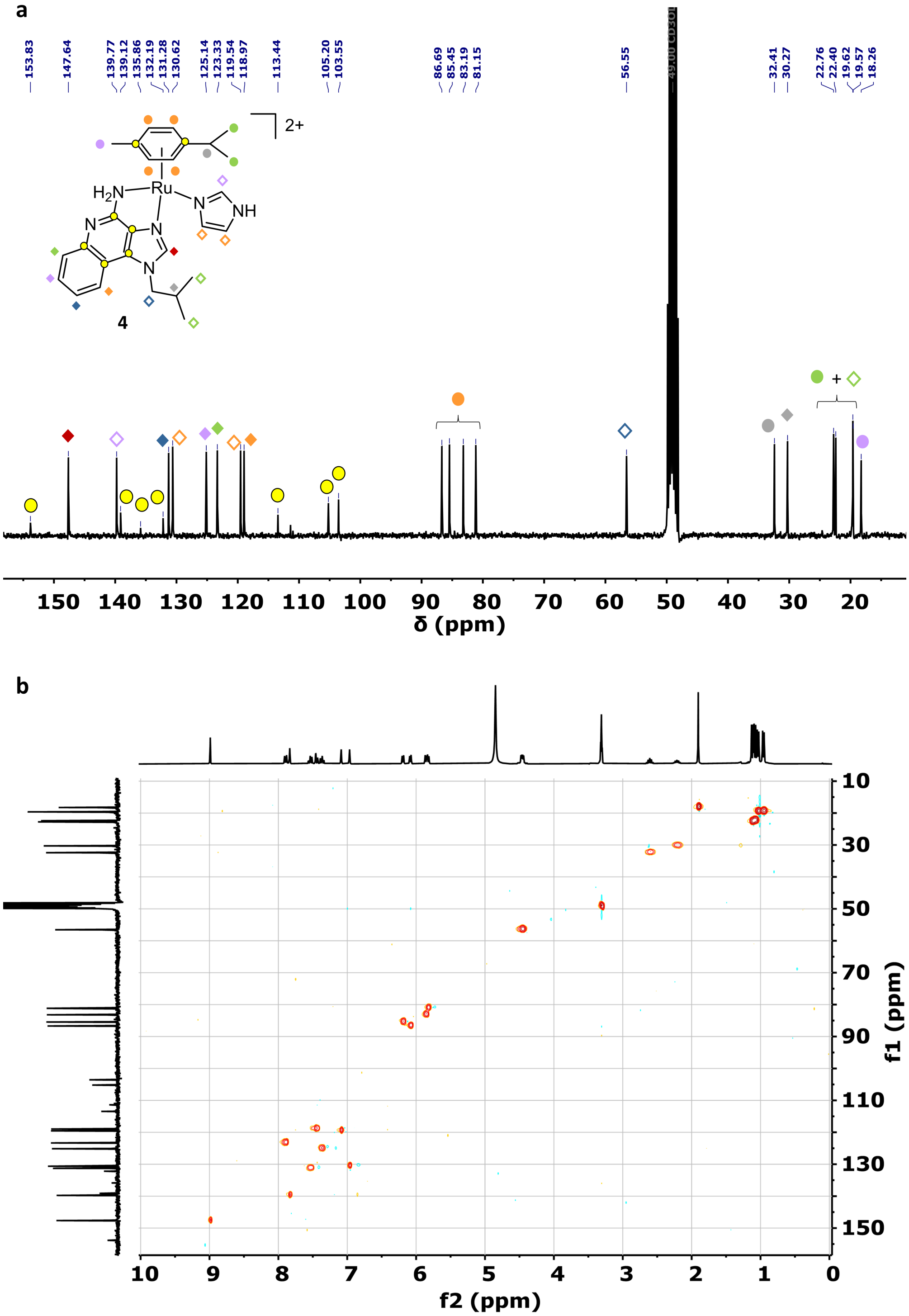


**Figure S9.** NMR spectra of complex [Ru(η$^6$-*p*-cym)(imq)(im)]$(PF_6)_2$ (**4**). (a) $^{13}$C (75 MHz, methanol-*d*$_4$, 298.5 K) and (b) $^1$H-$^{13}$C HSQC (300 and 75 MHz, methanol-*d*$_4$, 298.5 K).

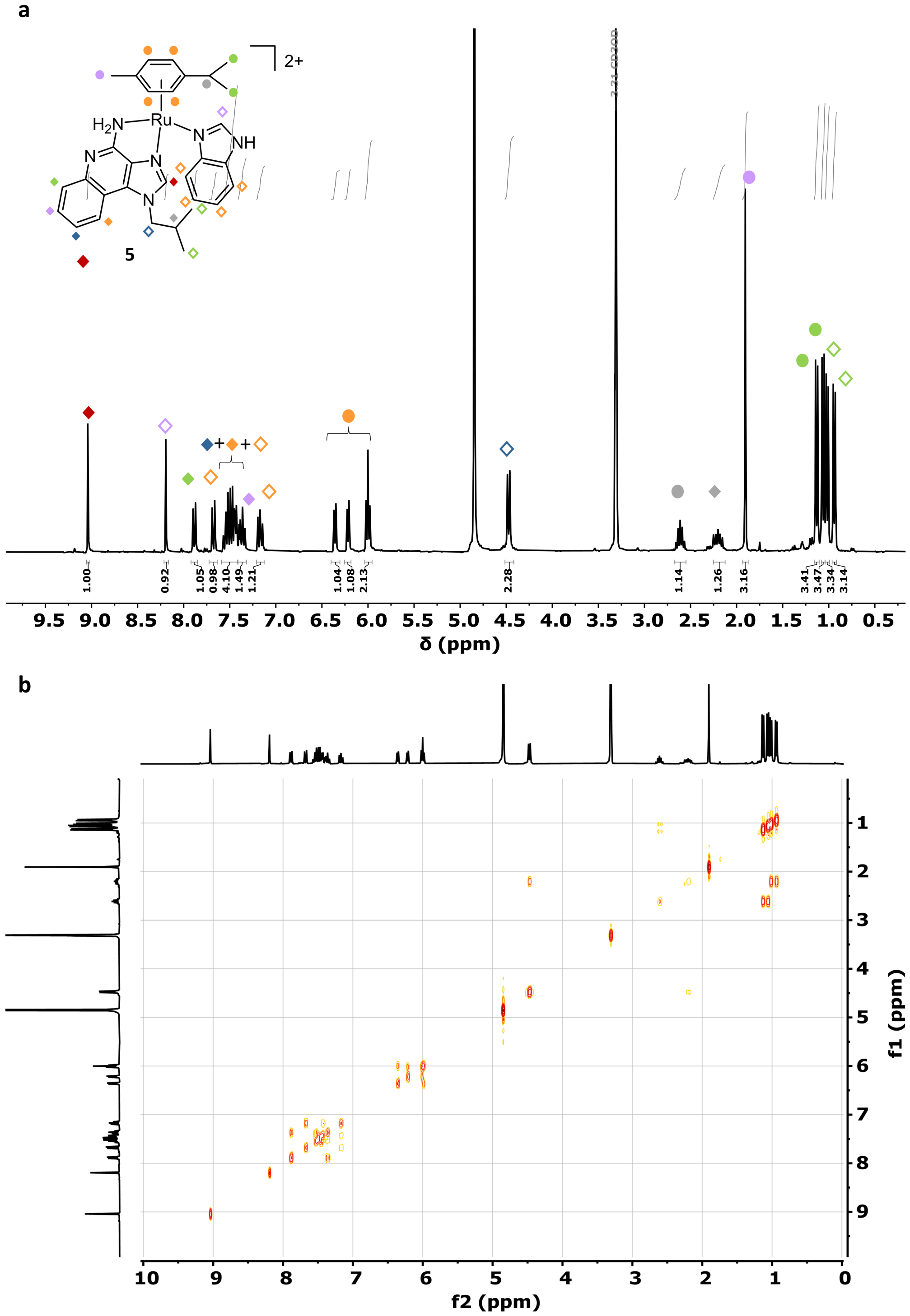


**Figure S10.** NMR spectra of complex [Ru(η$^6$-*p*-cym)(imq)(in)]($PF_6$)$_2$ (**5**). (a) $^1$H (300 MHz, methanol-*d*$_4$, 298.5 K) and (b) $^1$H-$^1$H COSY (300 MHz, methanol-*d*$_4$, 298.5 K).

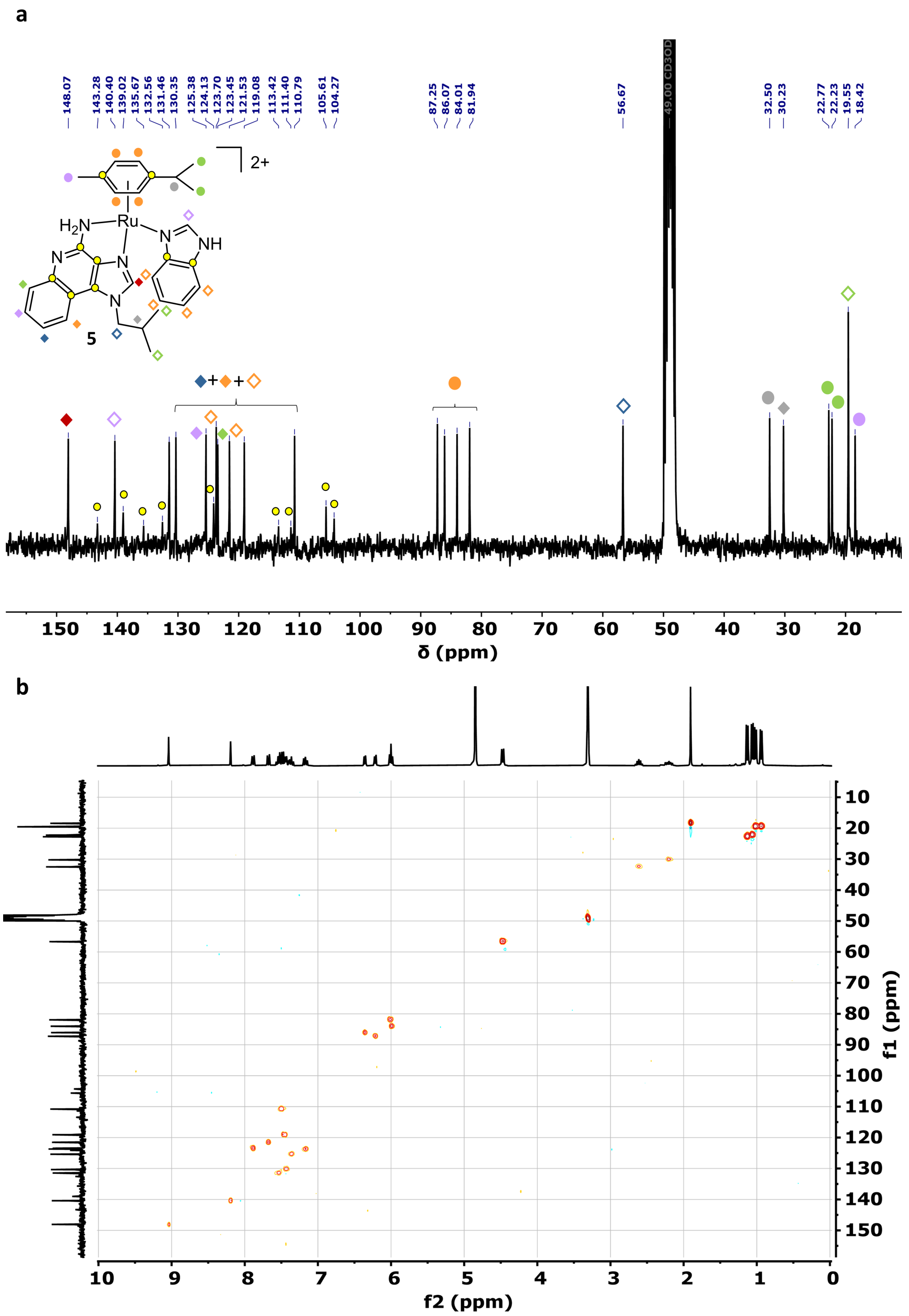


**Figure S11.** NMR spectra of complex [Ru(η$^6$-*p*-cym)(imq)(in)]($PF_6$)$_2$ (**5**). (a) $^{13}$C (75 MHz, methanol-$d_4$, 298.5 K) and (b) $^1$H-$^{13}$C HSQC (300 and 75 MHz, methanol-$d_4$, 298.5 K).

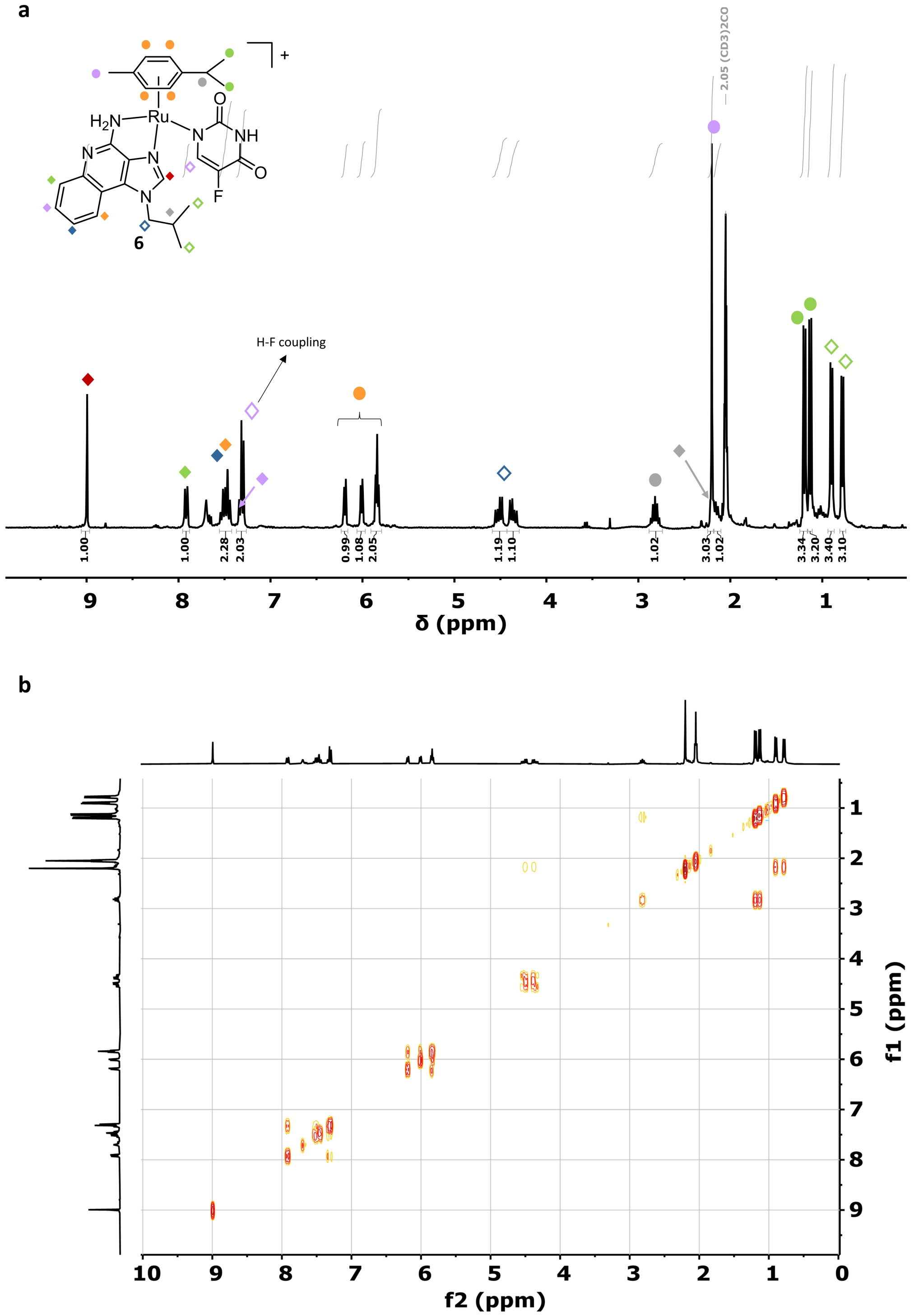


**Figure S12.** NMR spectra of complex [Ru(η$^6$-*p*-cym)(imq)(5-fu)]($PF_6$) (**6**). (a) $^1$H (300 MHz, acetone-$d_6$, 298.5 K) and (b) $^1$H-$^1$H COSY (300 MHz, acetone-$d_6$, 298.5 K).

a

b

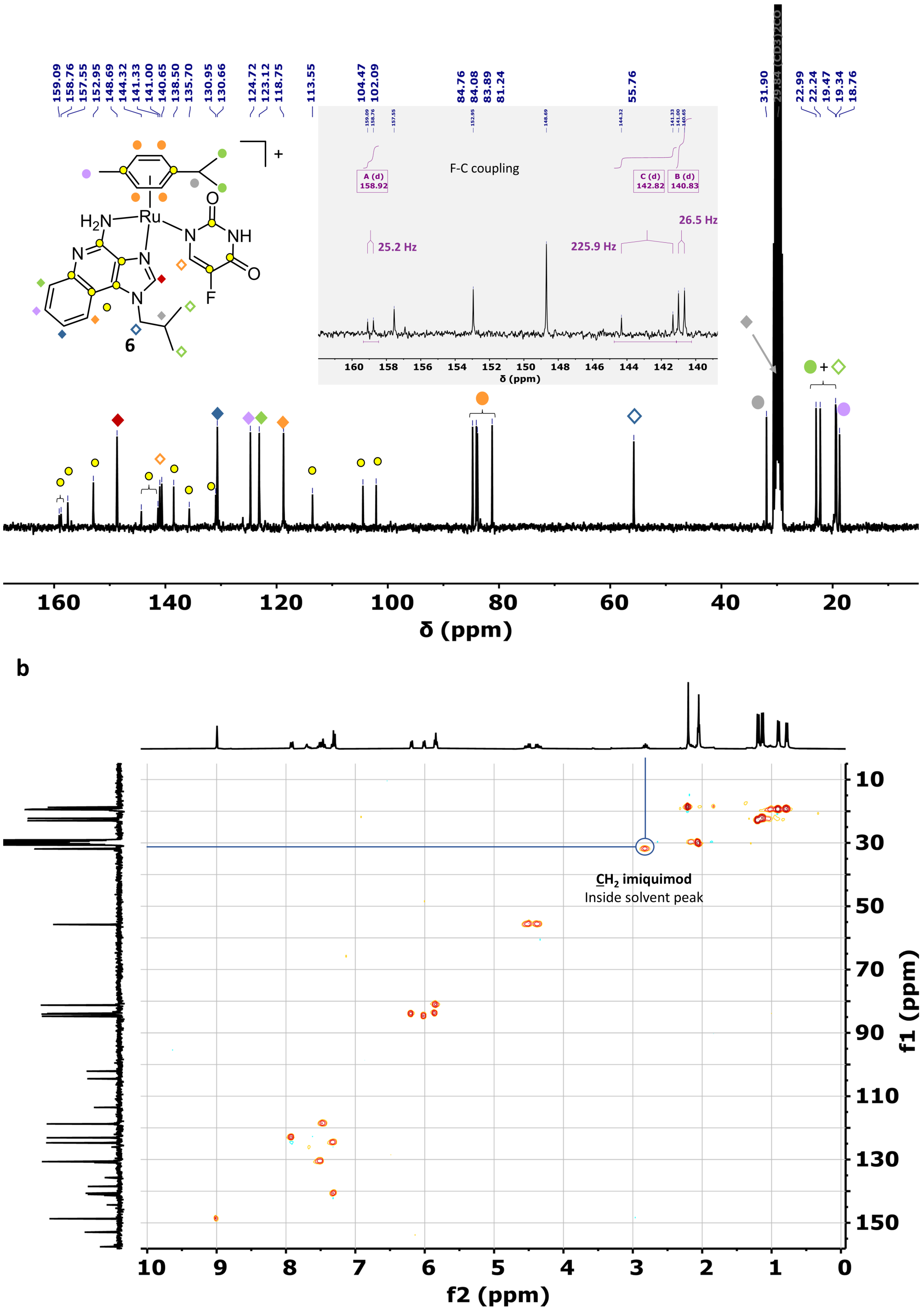

**Figure S13.** NMR spectra of complex [Ru(η[6]-*p*-cym)(imq)(5-fu)]($PF_6$) (**6**). (a) $^{13}C$ (75 MHz, acetone-$d_6$, 298.5 K) and (b) $^{1}H$-$^{13}C$ HSQC (300 and 75 MHz, acetone-$d_6$, 298.5 K).

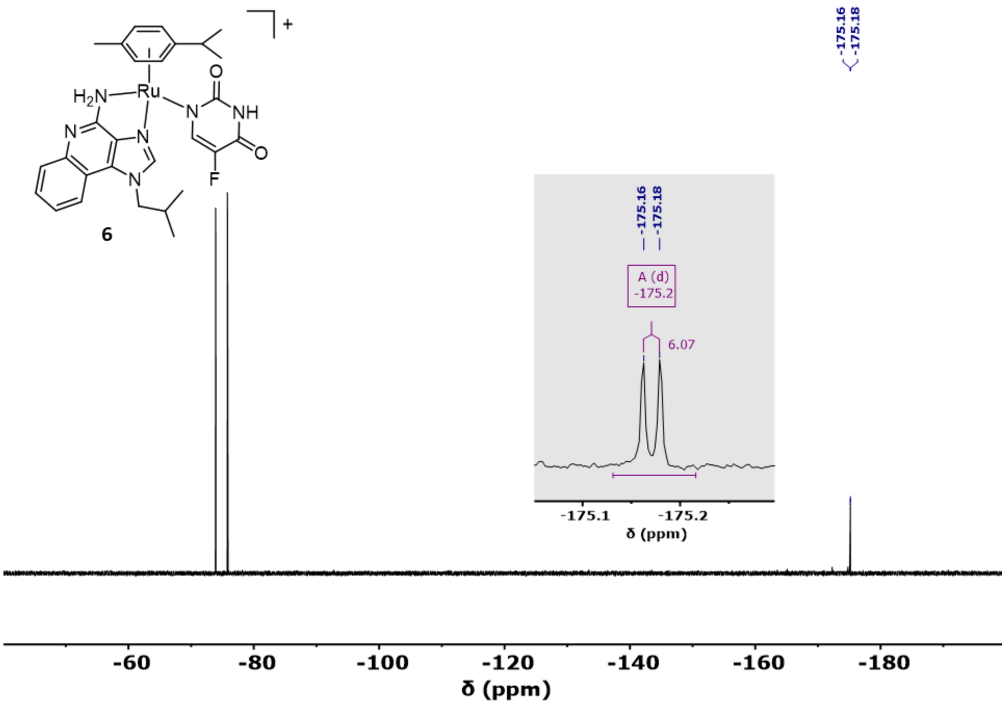


**Figure S14.** $^{19}$H NMR spectra of complex [Ru(η$^6$-*p*-cym)(imq)(5-fu)]($PF_6$) (**6**) (376 MHz, methanol-$d_4$, 298.5 K).

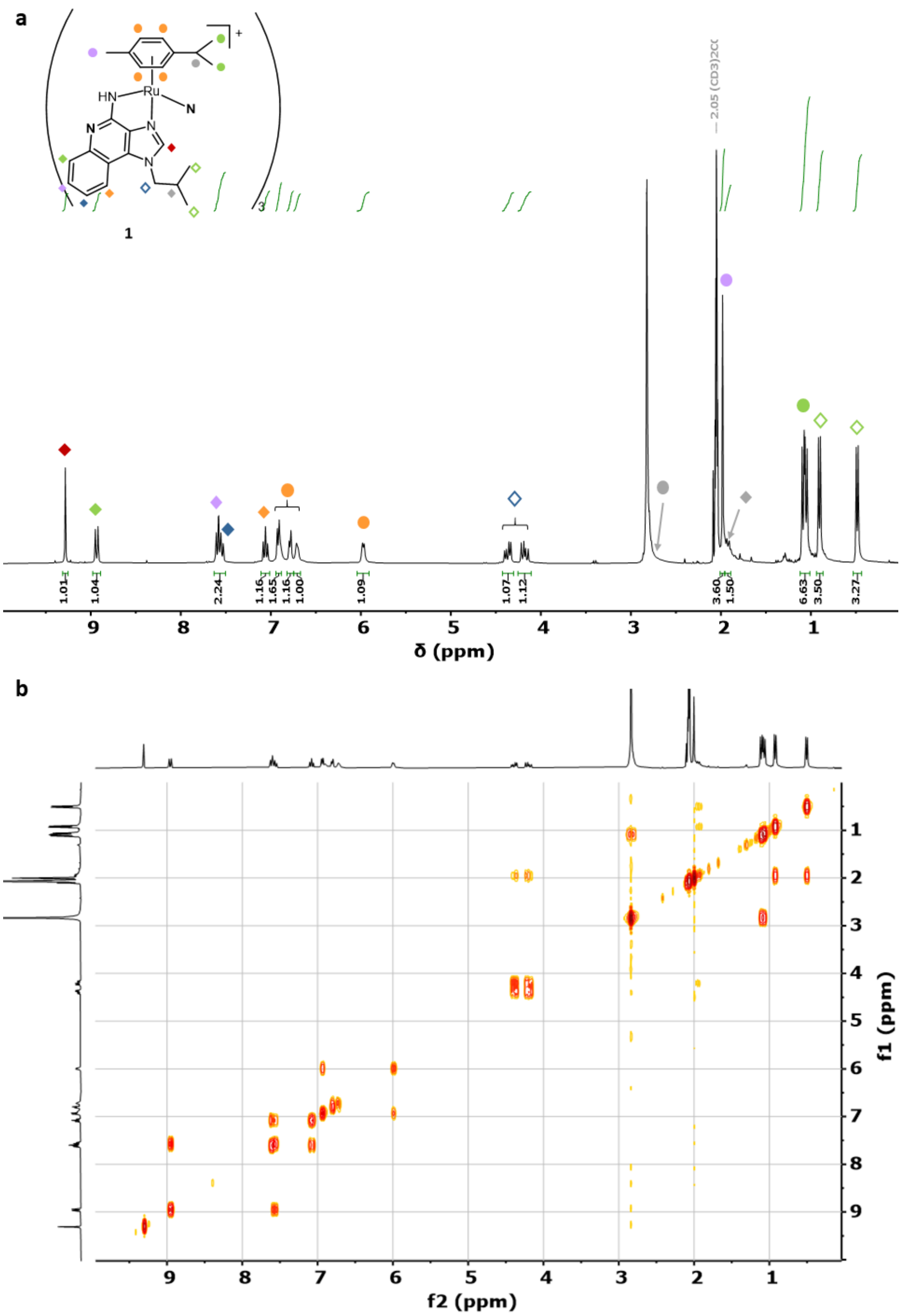


**Figure S15.** NMR spectra of complex [Ru($\eta^6$-*p*-cym)(imq)]$_3$(PF$_6$)$_3$ (**7**). (a) $^1$H (300 MHz, acetone-d6, 298.5 K) and (b) $^1$H-$^1$H COSY (300 MHz, acetone-d6, 298.5 K).

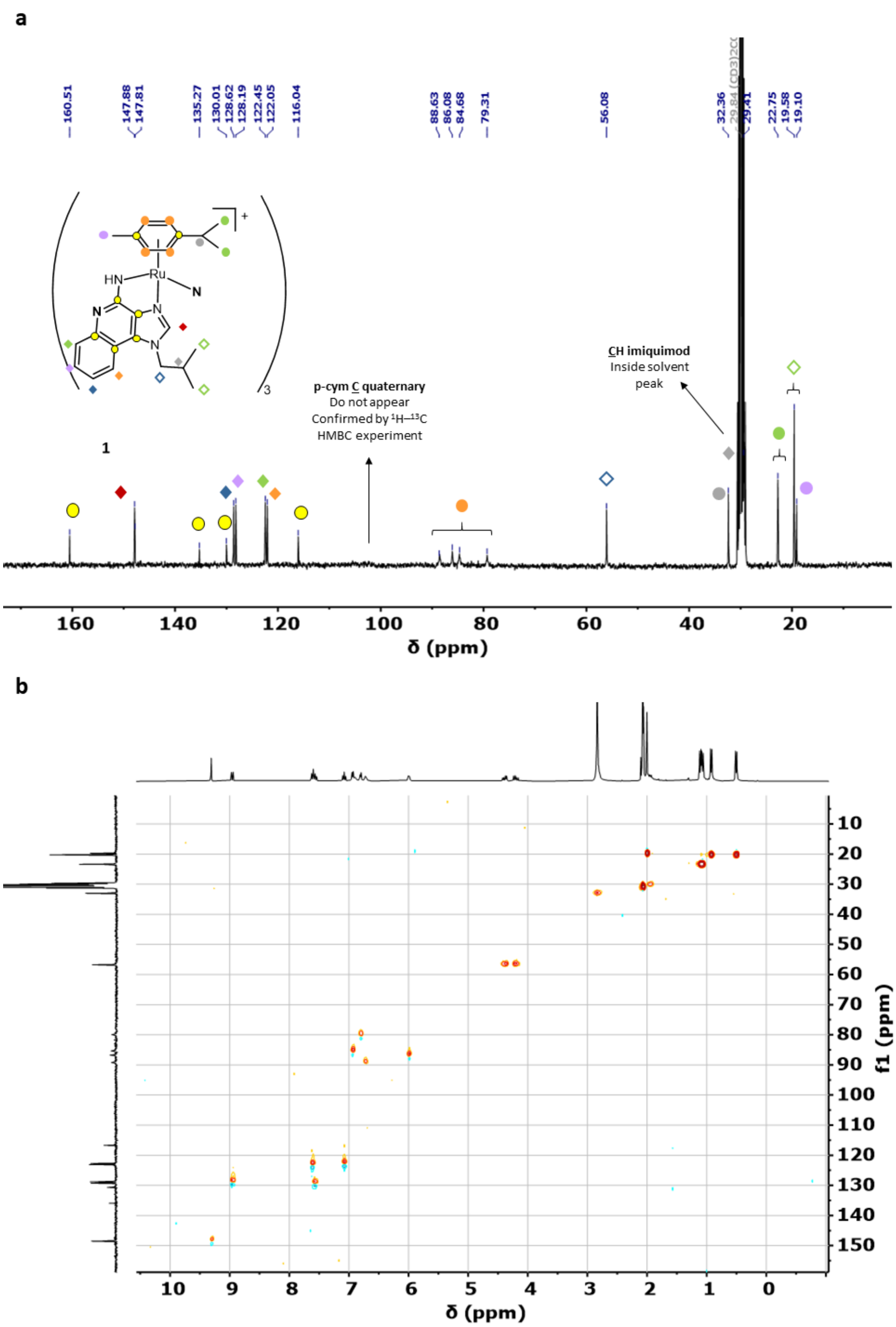


**Figure S16.** NMR spectra of complex [Ru(η$^6$-*p*-cym)(imq)]$_3$(PF$_6$)$_3$ (**7**). (a) $^{13}$C (75 MHz, acetone-*d*$_6$, 298.5 K) and (b) $^1$H-$^{13}$C HSQC (300 and 75 MHz, acetone-*d*$_6$, 298.5 K).

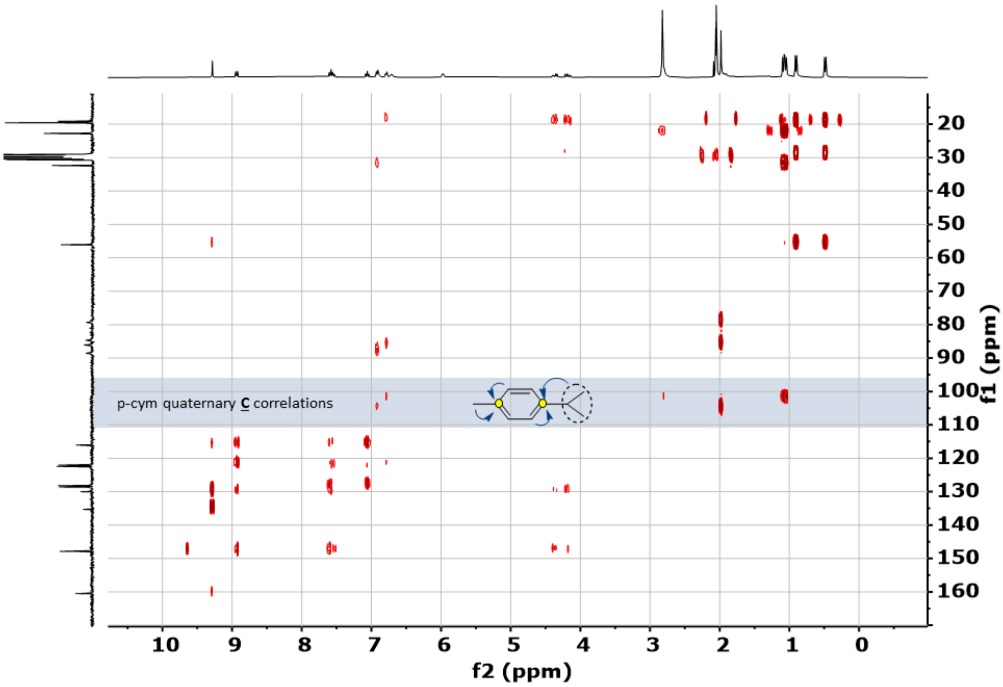


**Figure S17.** $^{1}H$-$^{13}C$ HMBC NMR spectra of complex [Ru(η$^6$-*p*-cym)(imq)]$_3$(PF$_6$)$_3$ (**7**) (300 and 75 MHz, acetone-$d_6$, 298.5 K).

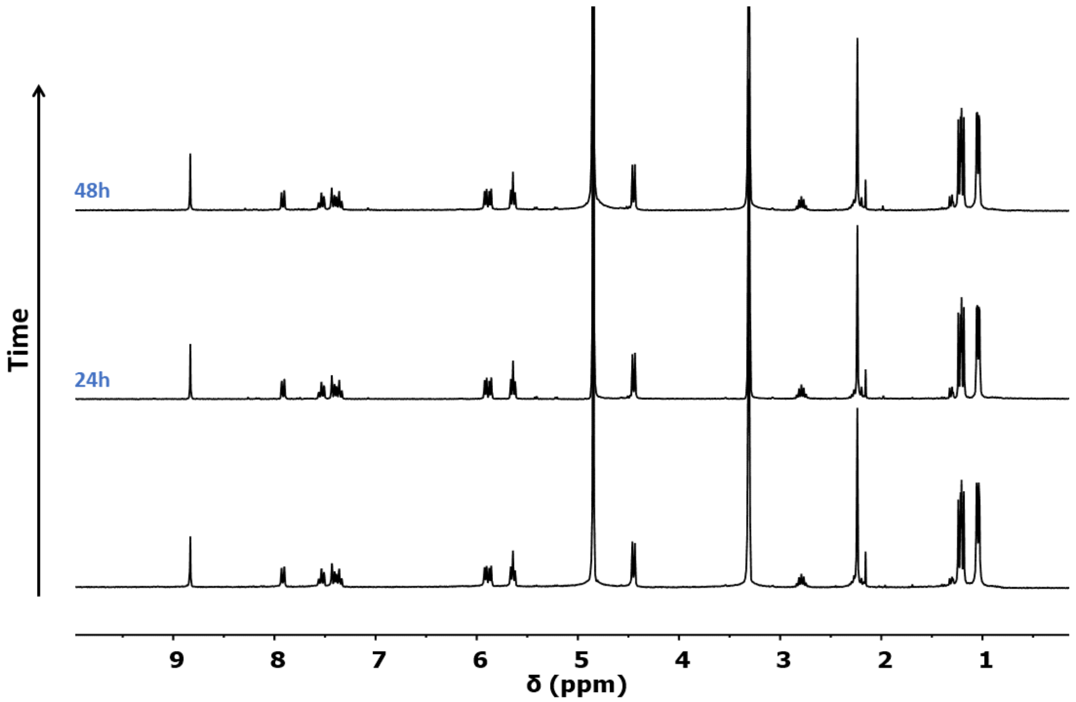


**Figure S18.** $^1$H NMR spectra of compound **1** in methanol-$d_4$ at different times (0, 24 and 48 h).

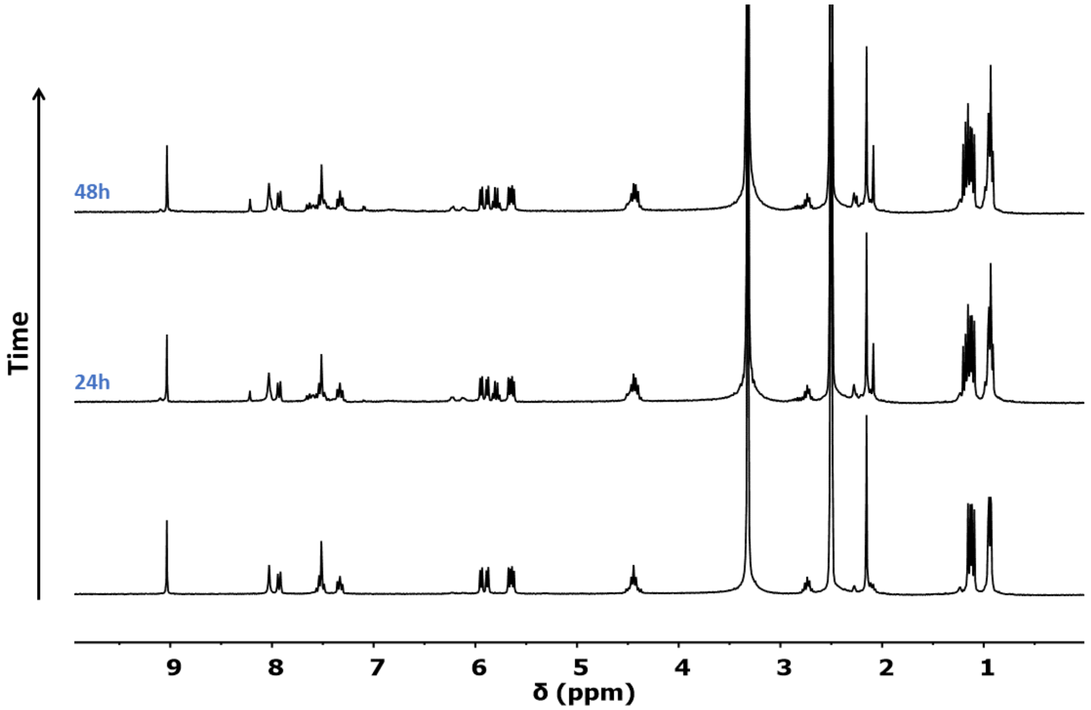


**Figure S19.** $^1$H NMR spectra of compound **1** in dimethyl sulfoxide-$d_6$ at different times (0, 24 and 48 h).

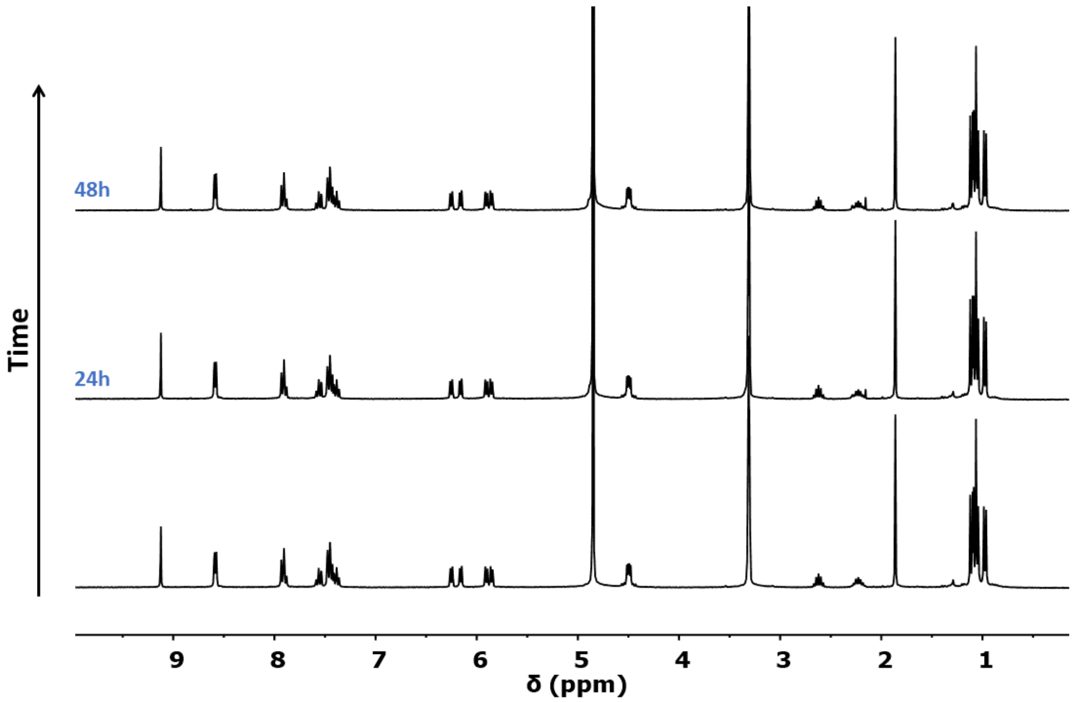


**Figure S20.** [1]H NMR spectra of compound **2** in methanol-$d_4$ at different times (0, 24 and 48 h).

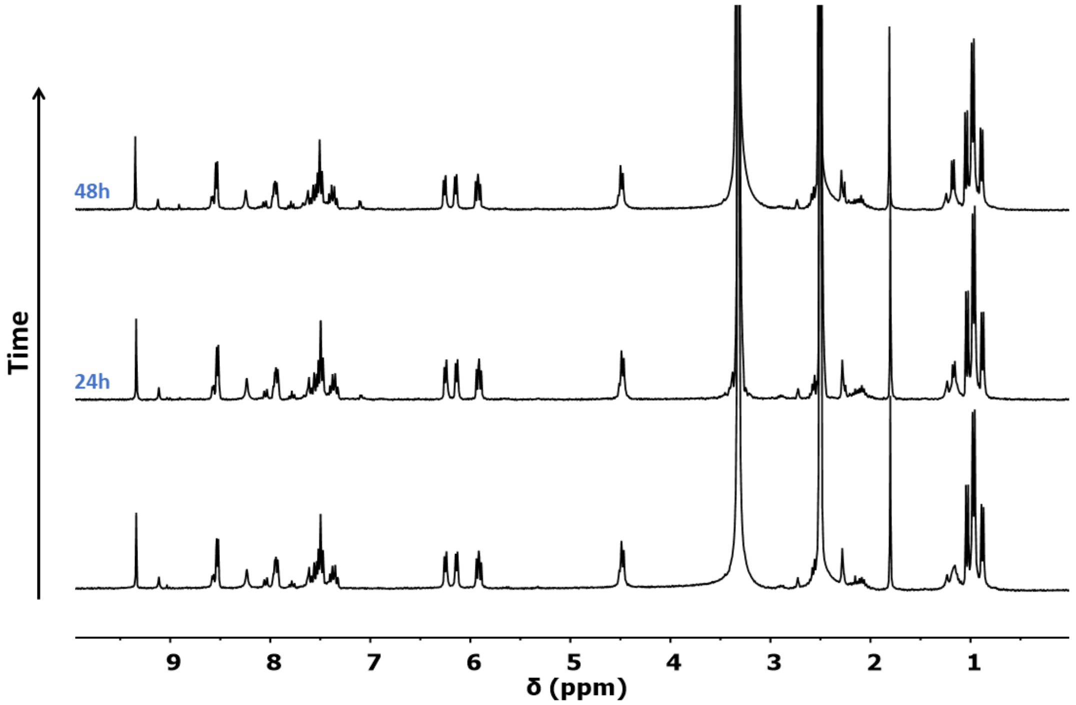


**Figure S21.** [1]H NMR spectra of compound **2** in dimethyl sulfoxide-$d_6$ at different times (0, 24 and 48 h).

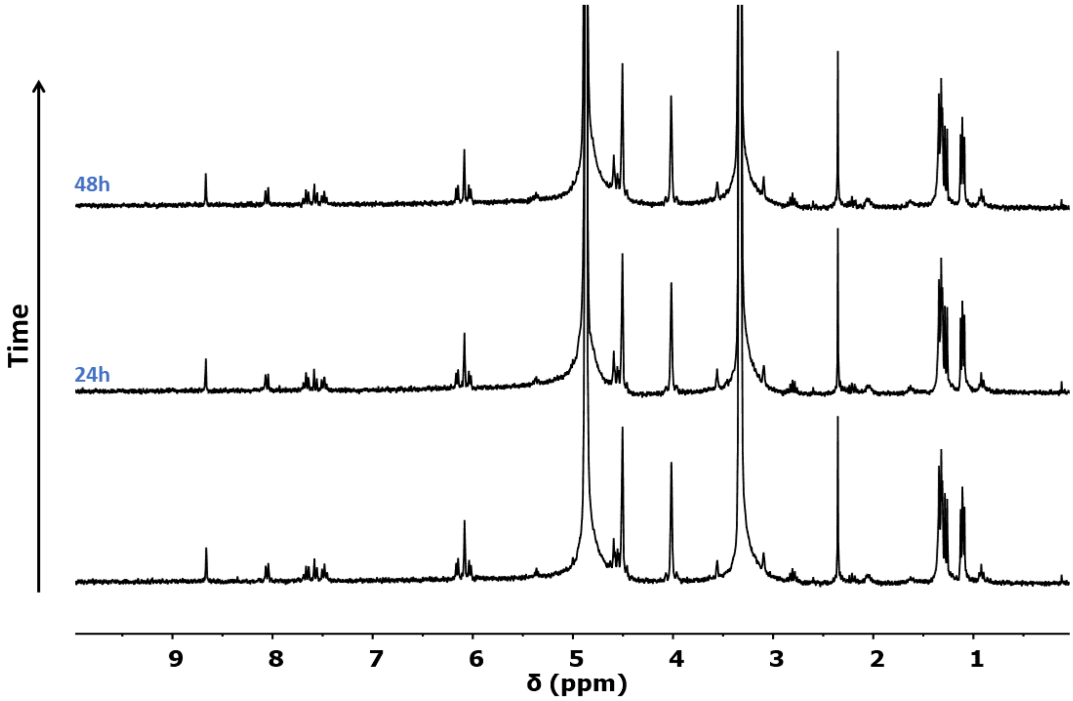


**Figure S22.** $^1$H NMR spectra of compound **3** in methanol-$d_4$ at different times (0, 24 and 48 h).

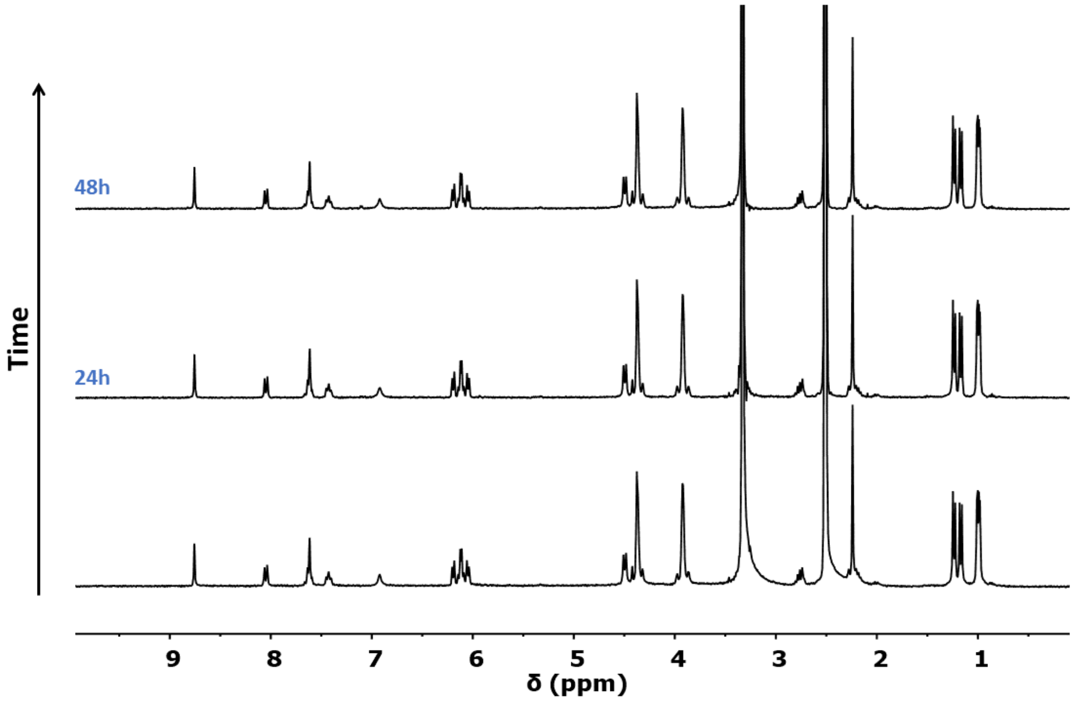


**Figure S23.** $^1$H NMR spectra of compound **3** in dimethyl sulfoxide-$d_6$ at different times (0, 24 and 48 h).

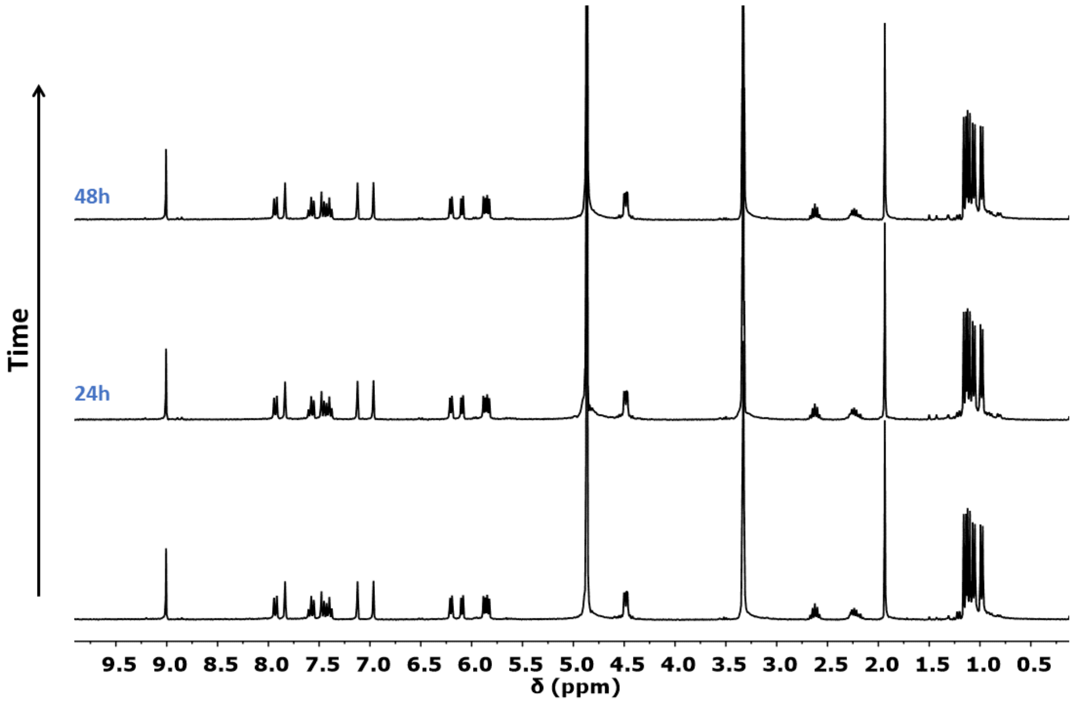


**Figure S24.** [1]H NMR spectra of compound **4** in methanol-$d_4$ at different times (0, 24 and 48 h).

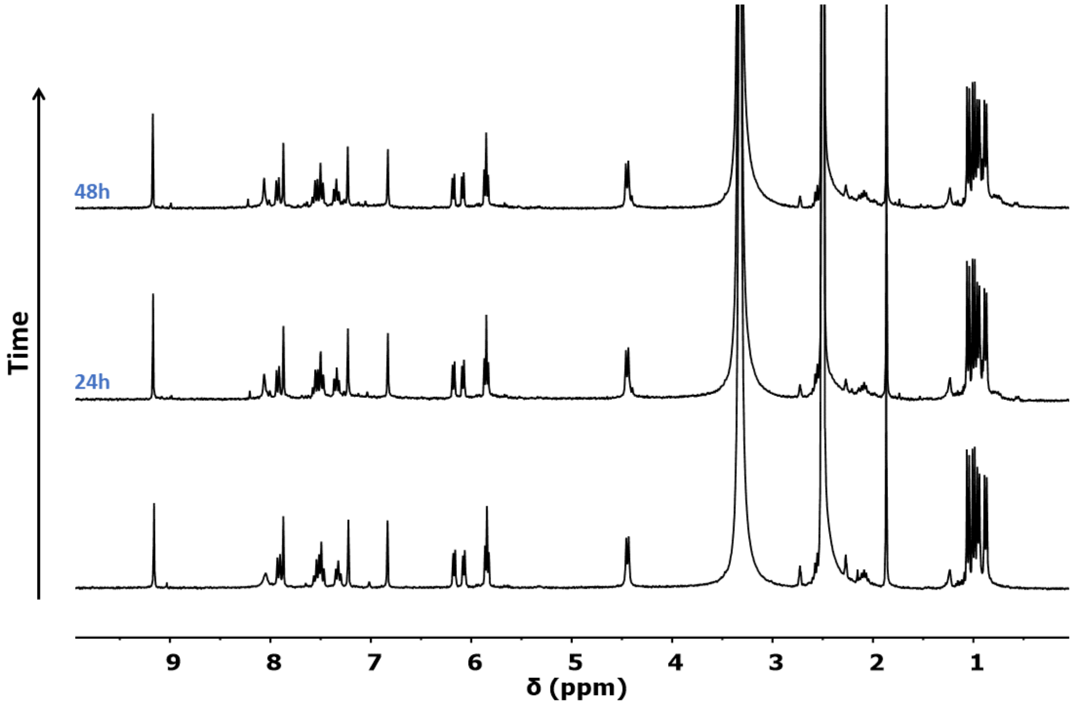


**Figure S25.** [1]H NMR spectra of compound **4** in dimethyl sulfoxide-$d_6$ at different times (0, 24 and 48 h).

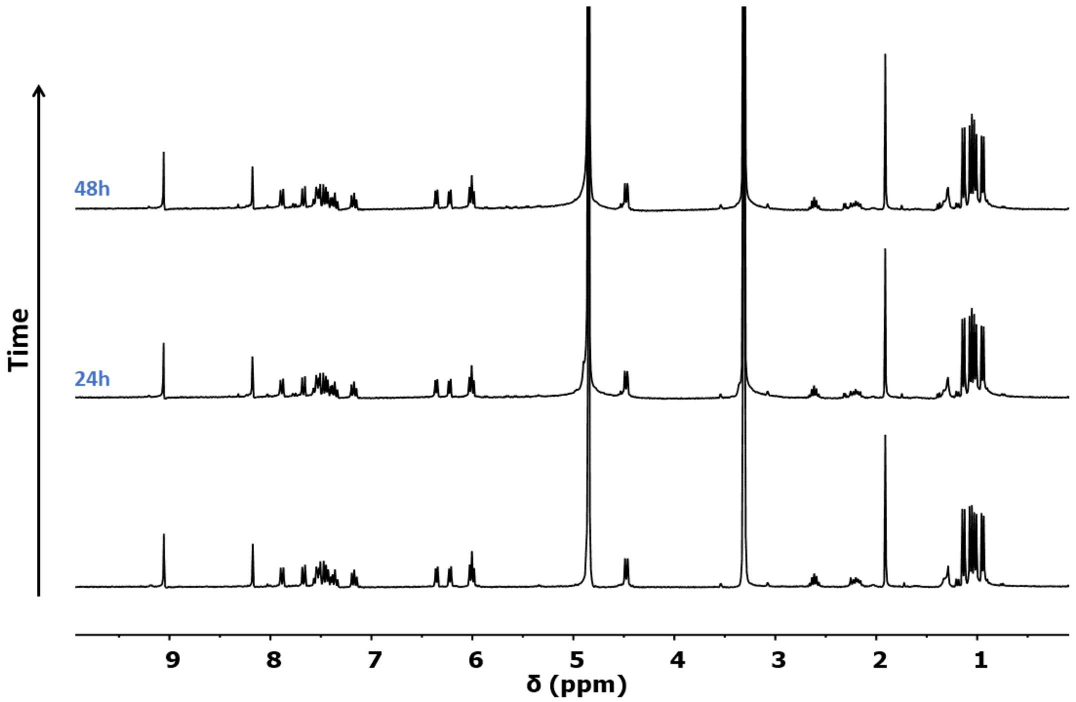


**Figure S26.** $^1$H NMR spectra of compound **5** in methanol-$d_4$ at different times (0, 24 and 48 h).

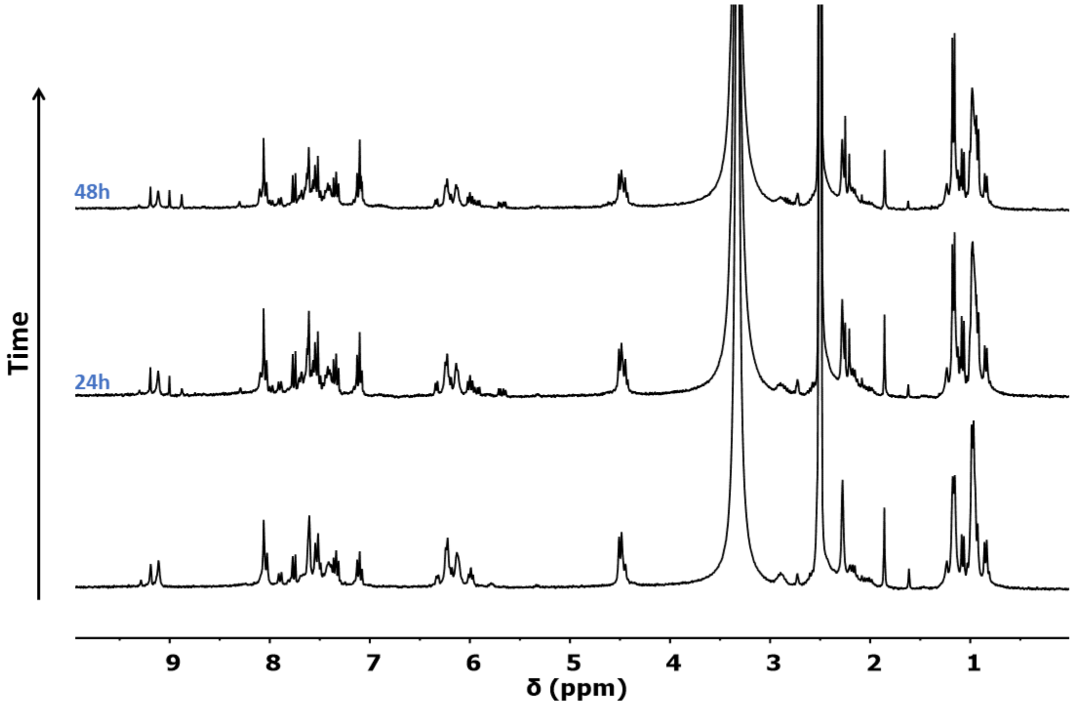


**Figure S27.** $^1$H NMR spectra of compound **5** in dimethyl sulfoxide-$d_6$ at different times (0, 24 and 48 h).

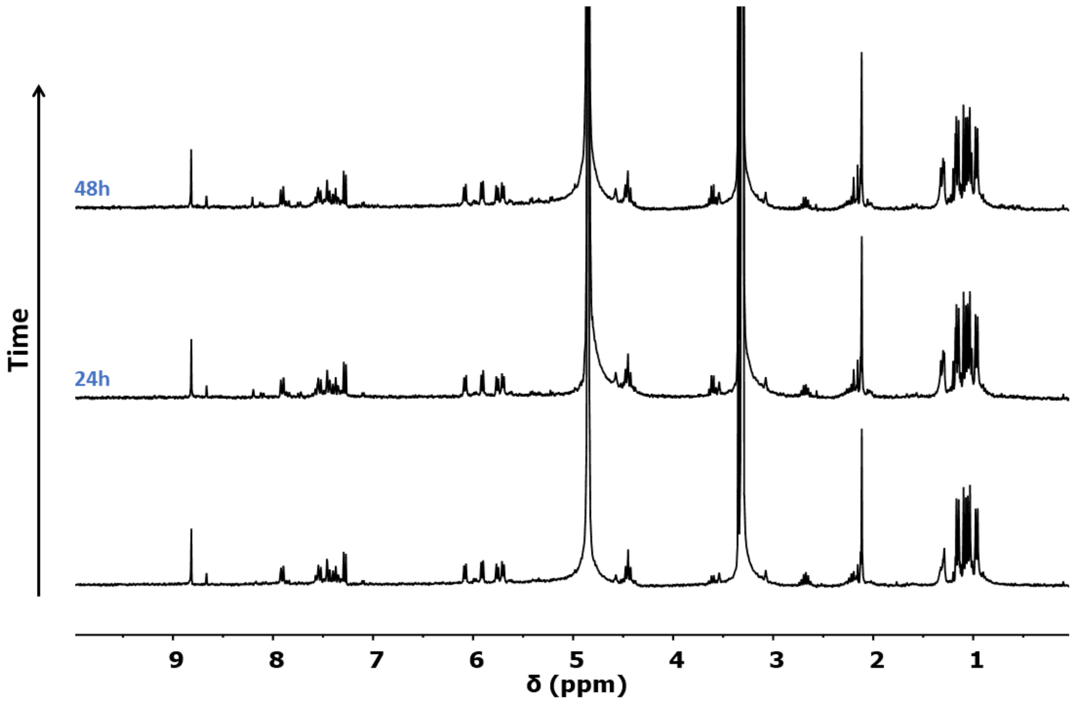


**Figure S28.** $^{1}$H NMR spectra of compound **6** in methanol-$d_4$ at different times (0, 24 and 48 h).

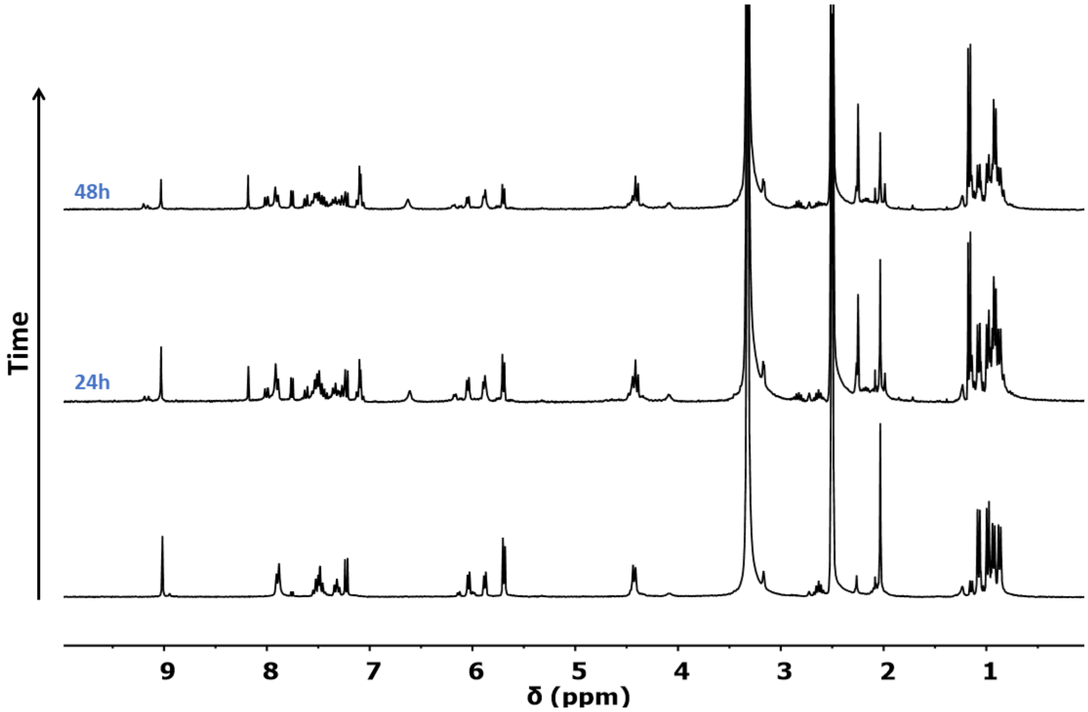


**Figure S29.** $^{1}$H NMR spectra of compound **6** in dimethyl sulfoxide-$d_6$ at different times (0, 24 and 48 h).

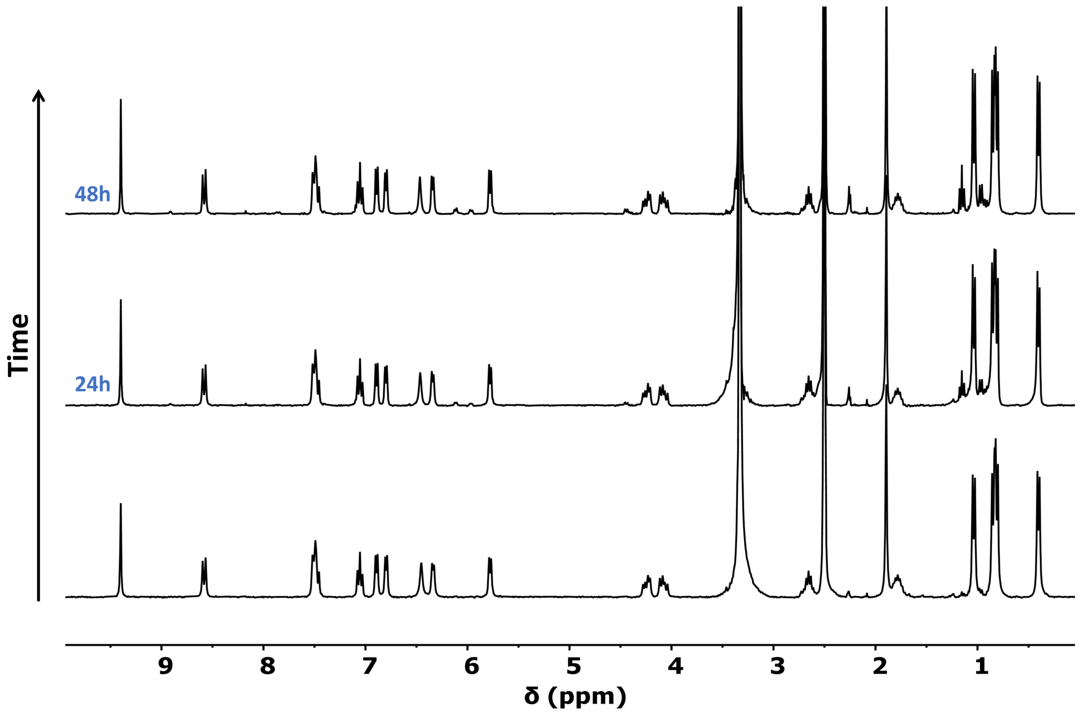

**Figure S30.** $^{1}$H NMR spectra of compound **7** in dimethyl sulfoxide-$d_6$ at different times (0, 24 and 48 h).

### 3. Tables

**Table S1.** Summary of the crystallographic data for complexes [Ru(η$^6$-*p*-cym)(imq)(Cl)]$PF_6$ (**1**), [Ru(η$^6$-*p*-cym)(imq)(py)]$(PF_6)_2$ (**2**), [Ru(η$^6$-*p*-cym)(imq)(im)]$(PF_6)_2$ (**4**), [Ru(η$^6$-*p*-cym)(imq)(5-fu)]$PF_6$ (**6**) and [Ru(η$^6$-*p*-cym)(imq)]$_3(PF_6)_3$ (**7**).

| | **1** | **2** | **4** | **6** | **7** |
|---|---|---|---|---|---|
| CCDC | 2583751 | 2583749 | 2583753 | 2583750 | 2583752 |
| Formula | $C_{48}H_{58}Cl_2F_{12}N_8P_2Ru_2$ | $C_{29}H_{35}F_{12}N_5P_2Ru$ | $C_{35}H_{52}F_{12}N_6O_2P_2Ru$ | $C_{28}H_{34}F_7N_6O_3P_2Ru$ | $C_{72}H_{87}F_{18}N_{12}P_3Ru_3$ |
| M | 1310.00 | 844.63 | 979.84 | 767.65 | 1859.04 |
| Crystal System | triclinic | triclinic | orthorhombic | monoclinic | Trigonal |
| Space group | P-1 | P-1 | Pbca | P21/n | P-3 |
| T [K] | 170 | 170 | 170 | 170 | 170 |
| a [Å] | 13.8085(4) | 10.6222(8) | 14.7277(1) | 12.9321(7) | 14.5859(2) |
| b [Å] | 13.8595(2) | 12.0711(6) | 18.3980(1) | 15.6713(11) | 14.5859(2) |
| c [Å] | 15.8550(4) | 13.6377(6) | 31.8961(2) | 16.0588(12) | 23.1498(3) |
| α [deg] | 94.846(2) | 83.159(4) | 90 | 90 | 90 |
| β [deg] | 109.858(2) | 74.055(5) | 90 | 92.345(6) | 90 |
| γ [deg] | 93.653(2) | 88.489(5) | 90 | 90 | 120 |
| V [Å$^3$] | 2829.83(14) | 1669.36(17) | 8642.58(9) | 3251.8(4) | 4264.49(10) |
| Z | 2 | 2 | 8 | 4 | 2 |
| Density [g·cm$^{-3}$] | 1.537 | 1.680 | 1.506 | 1.568 | 1.448 |
| μ [mm$^{-1}$] | 0.765 | 5.599 | 4.449 | 0.611 | 5.521 |
| Reflections collected | 13216 | 6795 | 8756 | 9353 | 5772 |
| $R_1$ [I > 2σ(I)] | 0.0888 | 0.0869 | 0.0391 | 0.0893 | 0.1308 |
| $wR_2$ [all data] | 0.2791 | 0.2587 | 0.1095 | 0.3012 | 0.3316 |
| GoF | 1.028 | 1.139 | 1.068 | 1.017 | 1.102 |

[a] $\sum||F_o|-|F_c||/\sum|F_o|$ [b] $\{\sum [w(F_o^2-F_c^2)^2]/ \sum [w(F_o^2)^2]\}^{½}$

**Table S2.** Cytotoxic effects of Ru complexes and imiquimod on HFF, HeLa, and VERO cells *in vitro*.[a]

| | $CC_{50}$[b] (μM) | | |
|---|---|---|---|
| **Compound** | **HFF** | **HeLa** | **VERO** |
| **1** | 84.94 | >100 | >100 |
| **2** | >100 | 67.33 | >100 |
| **3** | >100 | >100 | >100 |
| **4** | >100 | 57.07 | >100 |
| **5** | >100 | 53.78 | >100 |
| **6** | >100 | >100 | >100 |
| **7** | >100 | 17.95 | >100 |
| Imiquimod | 26.12 | 21.08 | >100 |

[a] The drug treatment period was 48h, and the cell viability was determined using the MTT cell viability assay. [b]$CC_{50}$ is defined as the compound concentration producing 50% cytotoxicity compared to the vehicle. Imiquimod was used for comparison. Experiments were performed in triplicate, and data are displayed as an average of three independent experiments.

**Table S3.** Cellular accumulation of Ru complexes (as ng Ru/$10^6$ cells) in different cell lines and different incubation times.

| Compound | HFF | | | | HeLa | | | |
|---|---|---|---|---|---|---|---|---|
| | 25 µM | | 50 µM | | 12.5 µM | | 25 µM | |
| | 8 h | 24 h | 8 h | 24 h | 8h | 24 h | 8 h | 24 h |
| **1** | - | - | 6.0 ±0.5 | 9.1±0.2 | - | - | 7.6±0.3 | 10.4±0.2 |
| **6** | - | - | - | - | 7.1±0.3 | 5.7±0.3 | - | - |
| **7** | 485±21 | 819±14 | - | - | 245±7 | 442±15 | - | - |
